# Promoter repression and 3D-restructuring resolves divergent developmental gene expression in TADs

**DOI:** 10.1101/2021.10.08.463672

**Authors:** Alessa R. Ringel, Quentin Szabo, Andrea M. Chiariello, Konrad Chudzik, Robert Schöpflin, Patricia Rothe, Alexandra L. Mattei, Tobias Zehnder, Dermot Harnett, Verena Laupert, Simona Bianco, Sara Hetzel, Mai Phan, Magdalena Schindler, Daniel Ibrahim, Christina Paliou, Andrea Esposito, Cesar A. Prada-Medina, Stefan Haas, Peter Giere, Martin Vingron, Lars Wittler, Alexander Meissner, Mario Nicodemi, Giacomo Cavalli, Frédéric Bantignies, Stefan Mundlos, Michael I. Robson

## Abstract

Cohesin loop extrusion facilitates precise gene expression by continuously driving promoters to sample all enhancers located within the same topologically-associated domain (TAD). However, many TADs contain multiple genes with divergent expression patterns, thereby indicating additional forces further refine how enhancer activities are utilised. Here, we unravel the mechanisms enabling a new gene, *Rex1,* to emerge with divergent expression within the ancient *Fat1* TAD in placental mammals. We show that such divergent expression is not determined by a strict enhancer-promoter compatibility code, intra-TAD position or nuclear envelope-attachment. Instead, TAD-restructuring in embryonic stem cells (ESCs) separates *Rex1* and *Fat1* with distinct proximal enhancers that independently drive their expression. By contrast, in later embryonic tissues, DNA methylation renders the inactive *Rex1* promoter profoundly unresponsive to *Fat1* enhancers within the intact TAD. Combined, these features adapted an ancient regulatory landscape during evolution to support two entirely independent *Rex1* and *Fat1* expression programs. Thus, rather than operating only as rigid blocks of co-regulated genes, TAD-regulatory landscapes can orchestrate complex divergent expression patterns in evolution.

**HIGHLIGHTS:**

- New genes can emerge in evolution without taking on the expression pattern of their surrounding pre-existing TAD.
- Compartmentalisation can restructure seemingly evolutionarily stable TADs to control a promoter’s access to enhancers.
- Lamina-associated domains neither prevent transcriptional activation nor enhancer-promoter communication.
- Repression rather than promoter-specificity refines when genes respond to promiscuous enhancer activities in specific tissues.

## INTRODUCTION

During development, enhancers with diverse activities drive extraordinarily complex transcription at target genes in time and space (Long et al., 2016). Remarkably, such enhancers can activate target genes lying hundreds of kilobases away by physically contacting promoters in three-dimensional space via chromatin folding (Bonev and Cavalli, 2016; Furlong and Levine, 2018). Collectively, this allows many developmental loci to be regulated by complex modular ensembles of enhancers distributed within large gene regulatory landscapes (Robson et al., 2019). However, how distal-acting enhancers are directed to only selected target promoters within regulatory landscapes has remained a central question.

In recent years, the 3D organisation of the genome has emerged as one such critical regulator of enhancer activities. Regulatory landscapes are partitioned into preferentially self-interacting blocks termed topologically-associated domains (TADs) by cohesin and the zinc-finger transcription factor CTCF (Dixon et al., 2012; Nora et al., 2012; Rao et al., 2014). Cohesin is thought to form TADs by progressively extruding chromatin loops until blocked by CTCF-boundaries, thereby bringing distant loci into spatial proximity (Fudenberg et al., 2016; Sanborn et al., 2015). In this way, TADs support gene regulation by continuously driving promoters to preferentially sample all enhancers within the same but not neighbouring domains (Kane et al., 2021; Symmons et al., 2014; Zuin et al., 2021). As such, TADs and their enhancer landscapes are frequently conserved across cell types and species to coordinate transcription in development and evolution, respectively (Dixon et al., 2012; Fraser et al., 2015; Harmston et al., 2017; Krefting et al., 2018). Moreover, large structural variants (SVs) that break TAD boundaries can generate ectopic enhancer-promoter contacts that drive altered gene expression in both disease and evolution (Acemel et al., 2017; Real et al., 2020; Spielmann et al., 2018). Consequently, TADs provide a framework to understand the partitioning of regulatory information and to identify expression-altering genomic rearrangements in evolution and human disease.

Nonetheless, this simple framework cannot explain crucial features of gene regulation alone. Indeed, TADs seemingly buffer the effects of the extreme distances in regulatory landscapes, thereby allowing enhancer activities to be transmitted throughout a domain independently of position (Anderson et al., 2014; Zuin et al., 2021). However, many TADs contain multiple genes with non-overlapping expression despite all promoters contacting the same enhancers (Dixon et al., 2016). Moreover, mutations that create novel ectopic enhancer-promoter contacts within rearranged TADs frequently do so without driving corresponding gene misexpression or disease (Despang et al., 2019; Laugsch et al., 2019). Finally, many potentially beneficial genome configurations could likely not be explored in evolution if genes universally adopted the entire regulatory activities of rearranged TADs. As a result, though facilitating and delimiting enhancer-promoter contacts, additional mechanisms are proposed to govern how and when enhancer activities are utilised within TADs. For example, strict enhancer-promoter compatibility or rendering promoters inert through repression may enable their divergent expression within multi-gene TADs (Furlong and Levine, 2018). Alternatively, isolation at the nuclear envelope (NE) in repressive lamina-associated domains (LADs) may sequester specific promoters away from enhancers within shared TADs (van Steensel and Belmont, 2017).

Here, we systematically test mechanisms enabling differential expression within the model *Rex1/Fat1* multi-gene TAD during mouse development *in vivo*. Specifically, we comprehensively mapped enhancer usage and chromatin structure in multiple vertebrate species cell-types when *Fat1* is expressed with or without *Rex1*. We demonstrate *Rex1* emerged within the ancient *Fat1* TAD landscape in placental mammals but is independently regulated through two mechanisms. First, in mouse embryonic stem cells (ESCs), the TAD is restructured into smaller separated domains with differing NE-attachment and this is driven by compartmentalisation overriding cohesin loop extrusion. Consequently, *Rex1* and *Fat1* are independently activated by separate clusters of nearby enhancers. By contrast, in embryonic limbs, DNA methylation renders *Rex1* inert to functionally compatible *Fat1* enhancer activities despite their transmission throughout the intact NE-attached TAD. Collectively, these data demonstrate that mammalian gene expression is not necessarily controlled by strict enhancer-promoter compatibility within invariant TAD scaffolds.

Rather, multiple elaborate and non-overlapping patterns of gene expression can emerge within the same landscape during evolution through structural changes and selective promoter silencing.

## RESULTS

### Divergent gene expression is common within multi-gene TADs

Multiple studies have produced conflicting results concerning coordinated gene expression in TADs (Despang et al., 2019; Flavahan et al., 2016; Laugsch et al., 2019; Ribeiro et al., 2021; Shen et al., 2012; Zhan et al., 2017). Consequently, we employed available HiC to map the distribution of genes in TADs in mouse E11.5 limbs, cortical neurons (CN) and ESCs (Figure 1) (Bonev et al., 2017; Kraft et al., 2019). This revealed ∼12% of the ∼2400 TADs found in each tissue contained only a single gene that collectively were enriched in developmental GO-terms (Figures S1A and C) (Eden et al., 2009). Thus, as previously suggested, a fraction of developmental loci are isolated alone within dedicated TAD regulatory landscapes (Wu et al., 2021). Nevertheless, ∼88% of TADs contained multiple genes. By classifying these promoters into ubiquitous (Ubiq.) or non-ubiquitous (non-Ubiq.) expression classes, we found TADs on average contain 2.4 non-Ubiq. and 3.6 Ubiq. genes (Figures 1B and S1B-C) (see STAR methods). Thus, multi-gene TADs dominate in the genome and frequently contain multiple non-Ubiq. “developmental” and/or Ubiq. “housekeeping” genes.

**Figure 1.**
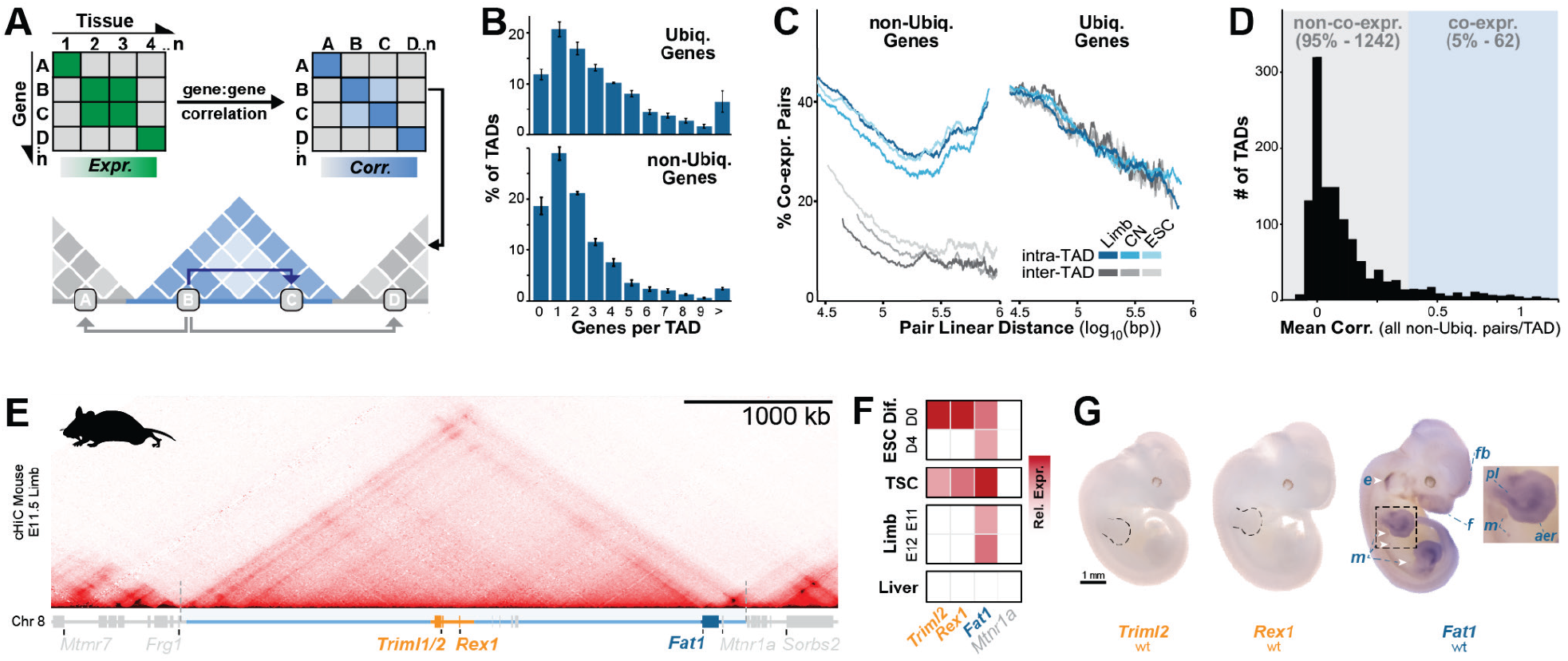
Promoters are poorly coregulated in most multi-gene TADs. **A.** Summary of method to determine correlation in expression between pairs of intra- or inter-TAD promoters. Promoter activity patterns across 329 mouse cell types and developmental stages were extracted from FANTOM5 CAGE data and the similarities between promoter pairs was determined by correlation analysis (Consortium et al., 2014; Lizio et al., 2015). TADs were identified from available HiC in embryonic E11.5 limbs, cortical neurons (CN) and ESCs (Bonev et al., 2017). **B.** Average frequency distribution of the number of non-Ubiq. and Ubiq. genes in limb, CN and ESC TADs. **C.** Fraction of co-expressing intra-TAD and inter-TAD gene pairs according to their linear separation. Pairs were considered co-expressing when correlation was in the top 10% for all genes (rho=0.38). Lines representing a moving window average of 2000 gene pairs. **D.** Frequency distribution of mean expression correlation for non-Ubiq. genes in a domain for all multi-gene TADs. **E.** cHiC at the *Rex1/Fat1* locus in E11.5 limb buds. Genes are shown as bars (*Fat1*, blue), *Mtnr1a* (grey), *Triml1*, *Triml2* and *Rex1* (orange). The ∼3.8 Mb *Rex1/Fat1* TAD (light blue) and the 293kb sub- *Rex1* region (*Rex1R*, orange) are also highlighted. Note that *Triml1* and *2* possess a shared bidirectional promoter (see Figure S1I). **F and G.** Gene activity overview from Fantom5 CAGE expression (F) and WISH (G). *Rex1* and *Triml2* are restricted to ESCs and trophoblast stem cells (TSCs). *Fat1* is expressed widely, including in the ear (e), mammary glands (m), face (f), forebrain (fb), proximal limb (pl) and apical ectodermal ridge (aer). Further HiC and scRNA-seq expression is available in Figures S1 and S2, respectively.

We next determined if multi-gene TADs support coordinated or divergent expression of their hosted genes in FANTOM5 nascent transcription datasets (Consortium et al., 2014). Specifically, we identified co-regulated pairs of Ubiq. or non-Ubiq. genes from their correlated expression changes across 329 mouse cell types and developmental stages (see STAR methods). Interestingly, pairs of non-Ubiq. genes were more frequently co-expressed when located within the same rather than different TADs at all length scales examined (Figure 1C) (Flavahan et al., 2016; Rennie et al., 2018; Zhan et al., 2017). By contrast, Ubiq. gene pairs remained equally co-expressed regardless of TAD co-occupancy. Thus, so-called “developmental” non-Ubiq. genes, but not their “housekeeping” Ubiq. counterparts, are more likely to share common regulatory instructions when located within shared TADs.

Nevertheless, despite this, TADs do not generally function to drive gene co-regulation. Most non-Ubiq. genes sharing a TAD are not co-regulated, and so only 5% of TADs display high mean co-regulation between *all* their constituent non-Ubiq. gene pairs (Figure 1D) (see STAR methods). Thus, multi-gene TADs largely do not behave as coordinated regulatory domains. Moreover, only ∼15% of the co-expressed promoter-pairs found along chromosomes share TADs, indicating most co-regulation is instead driven in *trans* (Figure S1D). Hence, though TADs restrict the regulatory information genes are exposed to, they are far from necessary or sufficient to determine co-regulation. Rather, other mechanisms must determine the subset of enhancers that most promoters utilise within a majority of TADs.

### Rex1 and Fat1 are differentially expressed despite sharing the same TAD

From the analysis above, we chose the representative *Rex1/Fat1* TAD to comprehensively dissect mechanisms enabling divergent expression in a multi-gene domain (Figure 1E). As confirmed by multiple cHiC and HiC datasets, this large TAD stably contains five genes in E11.5 limbs and multiple other mouse tissues (Figures 1E and S1E-H). Specifically, *Rex1*, *Triml1* and *Triml2* are positioned within a central 300 kb region (*Rex1R*) flanked by two ∼1.5 Mb gene deserts (D1 and D2). By contrast, *Fat1* and *Mtnr1a* genes are positioned near the TAD’s telomeric boundary. Thus, the mouse *Rex1*/*Fat1* locus fits the criteria of an apparently stable multi-gene TAD.

Despite occupying the same TAD, the *Rex1R, Fat1*, and *Mtnr1a* genes reportedly possess distinct transcriptional and functional properties. *Rex1* (*Zfp42*) is an extensively-studied transcription factor expressed during pluripotency (Masui et al., 2008). Similarly, *Triml1* and *Triml2* encode E3 ubiquitin ligase-like proteins with activity in pluripotency and a potential placental function (Zhang et al., 2020). By contrast, *Fat1* encodes an atypical cadherin possessing elaborate later embryonic expression and pleiotropic roles in tissue morphogenesis, cell growth and migration, and cancer (Peng et al., 2021). Finally, the melatonin receptor-encoding *Mtnr1a* contributes to circadian rhythm through highly restricted expression in, for example, the Suprachiasmatic nucleus and *pars tuberalis* (Klosen et al., 2019).

To confirm the previously reported divergent expression in this multi-gene TAD, we reanalysed available CAGE and single cell RNA-seq (scRNA-seq) datasets spanning mouse development (Figures 1F-G and S2) (Cao et al., 2019; Consortium et al., 2014; Lizio et al., 2015; Marsh and Blelloch, 2020; Pijuan-Sala et al., 2019). This revealed *Rex1R* genes and *Fat1* are co-transcribed in ESCs, placental trophoblasts and the extraembryonic ectoderm and endoderm (Figures 1F and S2A and B). Nevertheless, *Rex1R* genes are inactive after gastrulation despite continued *Fat1* transcription in a variety of tissues, including E11 limb buds (Figures 1F and S2C). Confirming this, whole mount *in situ* hybridisation (WISH) demonstrated elaborate *Fat1* activity in the E11.5 limb, ear, snout and mammary glands while *Rex1R* genes were undetectable, as previously reported (Figure 1G) (Ciani et al., 2003; Helmbacher, 2018; Kim et al., 2011; Zhang et al., 2020). Thus, though co-transcribed in some early developmental cell types, *Fat1* is largely independently expressed without *Rex1R* genes in most later tissues within their shared and stable TAD. By contrast, *Mtrn1a* expression was absent in all analyzed tissues, thereby excluding it from further analyses.

Collectively, these features made the *Rex1/Fat1* TAD an ideal candidate to study how highly functionally distinct genes achieve divergent expression in the same TAD.

### Independent Rex1R regulation emerged within Fat1’s ancient TAD landscape

TADs are frequently conserved as structural and regulatory units across species (Dixon et al., 2012; Fraser et al., 2015; Harmston et al., 2017; Krefting et al., 2018). We thus reasoned that divergent regulation of the functionally different *Fat1* and *Rex1R* genes emerged to accommodate a unique evolutionary history. As such, we examined the TAD and its enhancer landscape across the vertebrate evolutionary tree. HiC in multiple vertebrate species identified a conserved TAD that has been maintained at a largely constant length relative to diploid genome size despite frequent flanking synteny breaks (Figure 2A-D and S3). However, only *Fat1* universally occupies the TAD in all tested vertebrate species while *Triml1/2* and *Rex1* uniquely appear in eutherian placental mammals, including mice, humans and pigs (Figure 2A-C and S3) (Kim et al., 2007; Sadeqzadeh et al., 2014; Zhang et al., 2020). In particular, the *Rex1* gene itself reportedly emerged from a retroposion-driven duplication of the *Yin Yang 1* (*Yy1)* transcription factor in the eutherian lineage (Kim et al., 2007). Thus, the functionally distinct *Rex1R* genes emerged in eutherians long after *Fat1* and its conserved mono-gene TAD co-evolved in ancestral vertebrates.

**Figure 2.**
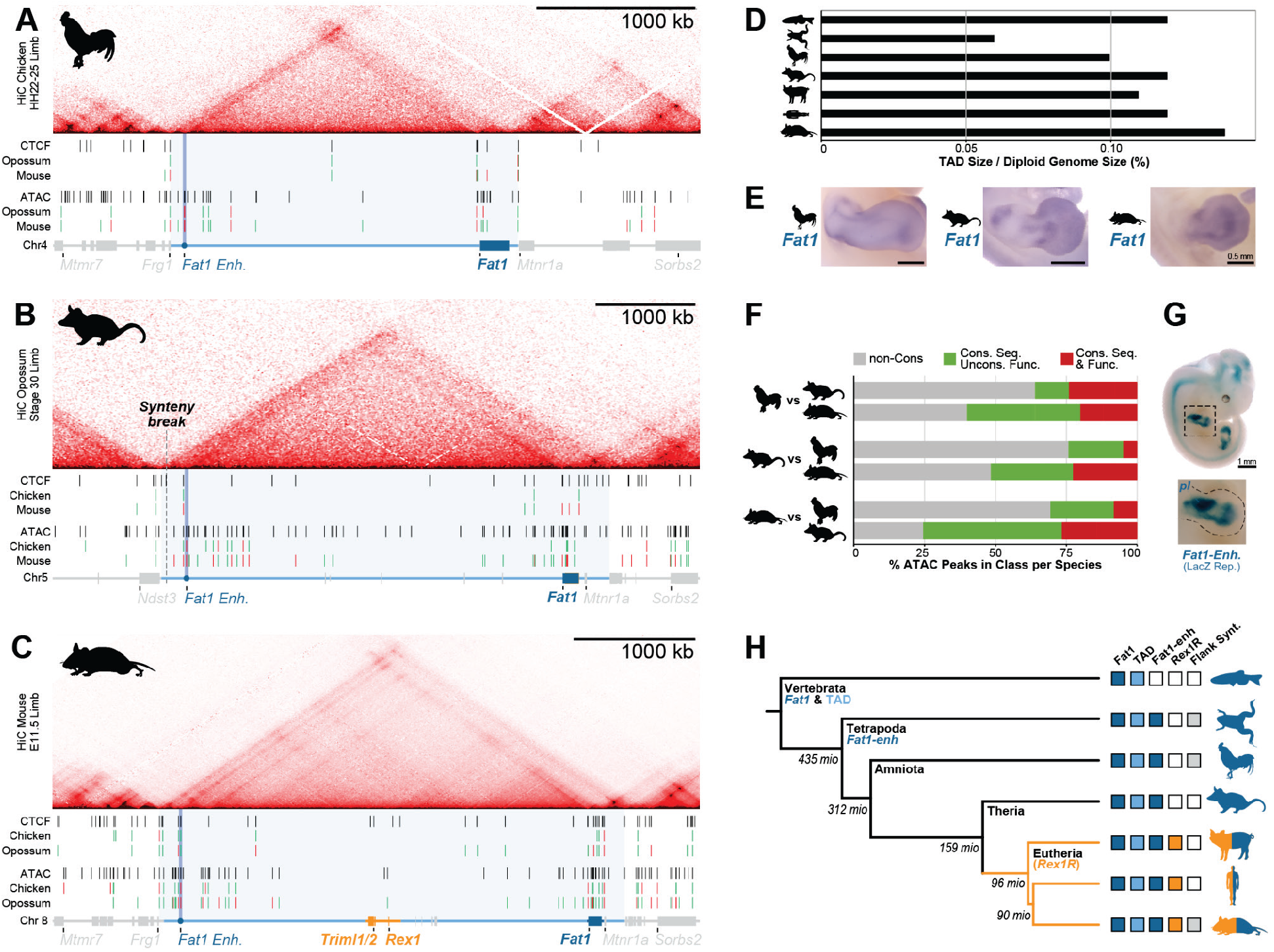
*Rex1R* genes emerged within *Fat1’s* ancient TAD-regulatory landscape. **A-C.** HiC from morphologically stage-matched chicken (A), opossum (B), and mouse (C) limb buds with matching ATAC-seq and CTCF ChIP-seq peaks shown below. ATAC and CTCF peaks are coloured according to their conservation of sequence with or without corresponding signal in the indicated species; red (seq.+, signal+); green (seq+, signal-); grey (seq-). An example ultra-conserved *Fat1* enhancer (*Fat1-enh.*) is highlighted (circle and dark blue box). **D.** Quantification of TAD size as fraction of diploid genome size in indicated species. **E.** Species-specific *Fat1* WISH in chicken, opossum and mouse embryonic limbs. **F.** Quantification of conservation of chicken, mouse or opossum ATAC-seq peaks in indicated species. Peaks are classified in each species comparison as with or without sequence conservation with or without matching functional conservation. **G.** LacZ reporter assay of cloned mouse *Fat1-enh* in E11.5 embryos when integrated at an ectopic locus. **H.** Phylogenetic tree with presence of *Fat1,* the TAD, *Fat1-enh, Rex1R* or flanking synteny outside the TAD indicated. See Figure S3.

We now assessed if the TAD originally evolved to solely regulate *Fat1*. Matching mouse development, *Fat1* displayed conserved expression in limbs from morphologically stage-matched chicken and opossum embryos (Figure 2E). Consequently, we further mapped putative enhancers likely driving this expression in the limbs of all three species by chromatin accessibility (ATAC-seq) (Figure 2A-C). This identified 25, 62 and 49 putative cis-regulatory elments in chicken, opossum and mouse, respectively, which clustered in the TAD’s distal arm or *Fat1*’s gene body (Figures 2F). Of these, 12-49% lacked ATAC signal in comparison species despite the presence of direct or indirect sequence conservation (Baranasic et al., 2021) (see STAR methods). Likewise, 24-76% of ATAC-peaks displayed no sequence conservation and were entirely species-specific. Thus, significant enhancer turnover occurred at the locus over time, perhaps to sustain or modify *Fat1* activity which has diverse critical physiological functions (Peng et al., 2021). Nevertheless, between 5-27% of putative elements were functionally conserved between each species and this universally included the *Fat1* promoter. Of these, a tetrapod-specific conserved enhancer located ∼3 Mb from *Fat1* drove *Fat1*-like activity in the proximal limb and neural tube when tested in a mouse lacZ reporter assay (Figure 2G) (see STAR methods). Thus, *Fat1* co-evolved with a conserved TAD and enhancers that likely drive its limb and, presumably, wider embryonic expression patterns (Figure 2H). By contrast, *Rex1R* appeared within the locus much later in placental mammals where it was able to evade *Fat1* enhancers and become independently regulated.

This demonstrates that novel genes and transcriptional programs can emerge in pre-established regulatory landscapes without compromising their existing functions. However, collectively, our genome-wide and evolutionary analyses indicate additional mechanisms must further refine how enhancer activities are used within multi-gene TADs.

### TAD restructuring in ESCs drives Fat1 and Rex1 to independently utilise local enhancers

We sought to identify the mechanisms adapting the ancient TAD landscape for independent *Fat1* and *Rex1R* gene regulation in placental mammals. Thus, we comprehensively mapped active enhancers and chromatin structure in mouse tissues where *Fat1* and *Rex1R* genes are divergently expressed (E11.5 limbs) or active together (ESCs) (Figure 3). Significantly, both *Rex1* and *Fat1* are dispensable for pluripotency and limb development, with the latter possessing functional redundancy with *Fat2, 3 or 4* (Ciani et al., 2003; Masui et al., 2008; Sadeqzadeh et al., 2014). As such, alterations to their regulation can be studied in ESCs and limbs without additional confounding effects.

**Figure 3.**
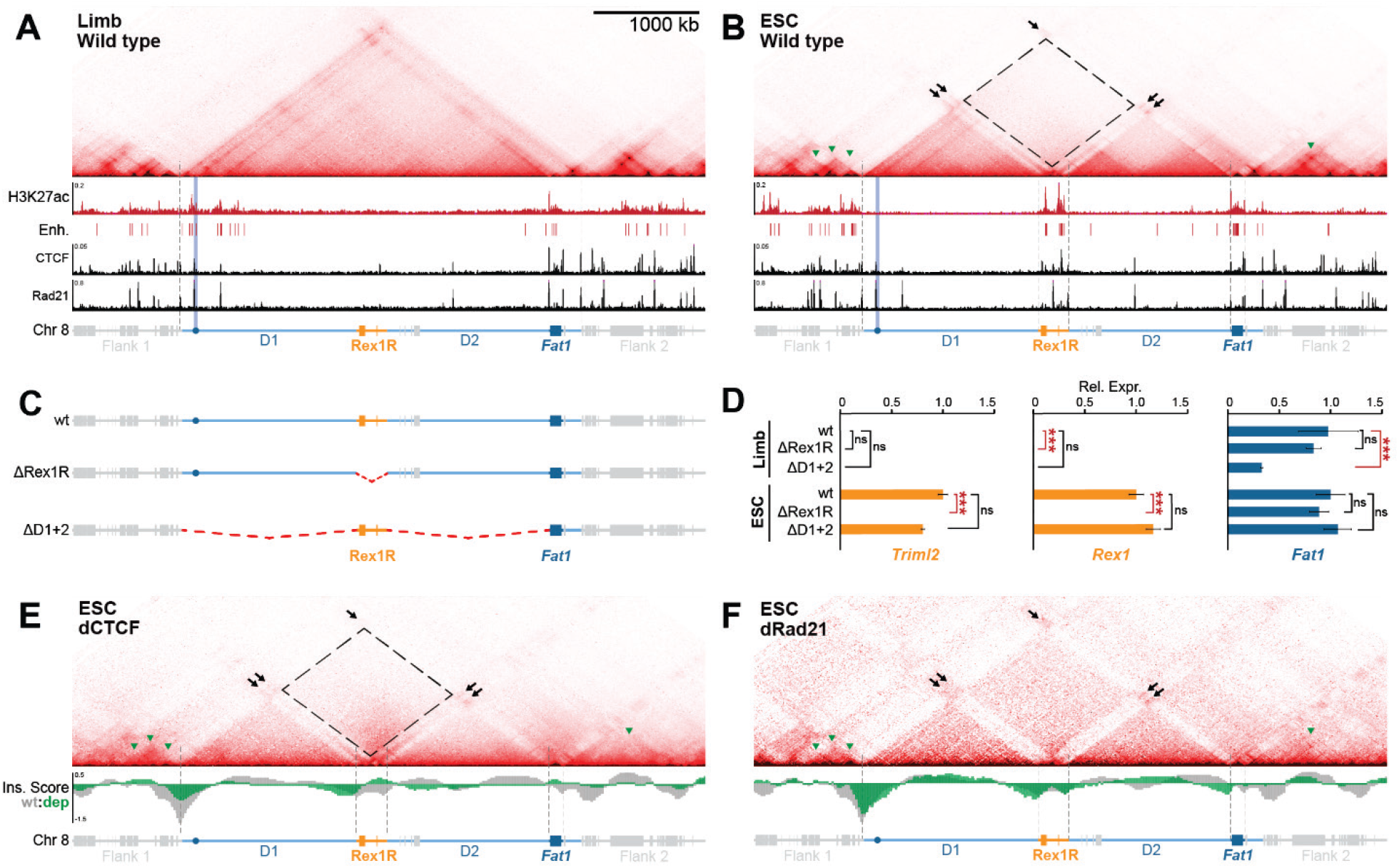
TAD-restructuring in ESCs isolates *Rex1R* and *Fat1* with proximal enhancers independently from cohesin and CTCF. A and B. cHiC from E11.5 limb buds (A) and ESCs (B) with H3K27ac, CTCF and Rad21 ChIPseq shown below. Black arrows indicate interactions between active H3K27ac-marked regions and dotted rectangle displays lost interaction between inactive H3K27ac-free gene deserts 1 and 2 (D1 and D2). **C and D.** Schematic of deletion mutants (C) with gene expression effects analyzed by RNA-seq (D). *Fat1* and *Rex1* require only local *Rex1R* or *Fat1* enhancers for expression in ESCs, respectively. However, *Fat1*, but not *Rex1*, utilises distal D1 enhancers in the embryonic limb. n=2-4 biological replicates per sample. *** p<0.001, * p<0.05 and ns p>0.001. **E and F.** cHiC from dCTCF (E) or dRad21 (F) ESCs. Corresponding insulation score profiles for wildtype (grey) or depletion (green) ESCs are shown below. The *Rex1*/*Fat1* locus partitioning persists following disrupted loop extrusion despite the loss of surrounding TADs (green arrows). See Figure S4.

In E11.5 limbs, this confirmed active H3K27ac-marked enhancers cluster near the intact TAD’s centromeric boundary and within *Fat1*’s gene body (Figures 3A and S4A) (Andrey et al., 2017). However, in ESCs, a radically different TAD structure and underlying enhancer landscape emerged. Here, ESC-enhancer activities are redistributed into two distinct clusters found locally within *Rex1R* and *Fat1’*s gene body (Figures 3B and S4A) (Bauer et al., 2021). Correspondingly, the TAD is organised into four domains separated according to activity. Specifically, *Rex1R* and *Fat1* eliminate interactions with the flanking D1 and D2 regions and become separately isolated with their local enhancers in individual active domains. In parallel, lost interactions between D1 and D2 create two additional separated inactive domains. Collectively, these alterations reproducibly restructure the TAD in cell types where *Fat1* and *Rex1R* genes are both simultaneously active, such as 8-cell mouse embryos and human ESCs (Figure S4B-D) (Bonora et al., 2020; Zhang et al., 2019). Thus, though evolutionarily stable, the ancient TAD has a flexible structure in pluripotent placental mammal cells that physically restricts *Fat1* and *Rex1R* with separate local enhancers.

We thus tested if this locus restructuring grants *Rex1R* and *Fat1* functional independence by generating E11.5 embryos and ESCs harbouring a series of deletions (Kraft et al., 2015). Specifically, we eliminated the placental mammal-specific *Rex1R* (ΔRex1R) or the ancient D1 and D2 regions (ΔD1, ΔD2 or ΔD1+2) (Figures 3C and S4E). RNA-seq in mutant E11.5 limb buds revealed *Fat1* expression was severely disrupted by deletion of the ancient gene deserts but not the more recently-emerged *Rex1R.* Specifically, eliminating putative centromeric limb enhancers in ΔD1 and ΔD1+2 mutants reduced limb-wide *Fat1* expression by 56-67% (Figures 3D and S4E). By contrast, *Rex1R* genes remained inactive in wildtype and all mutant limbs. Hence, in later development, elaborate *Fat1* expression is driven by an ancient TAD regulatory landscape and distal enhancers that have no effect on *Rex1R* gene expression.

In contrast, in ESCs, *Fat1* expression remained universally unaffected in ΔD1, ΔD2, ΔD1+2 and ΔRex1R mutants (Figures 3D and S4E). Similarly, *Rex1R* genes were unaffected by single ΔD1/ΔD2 or combined ΔD1+2 deletions, except *Triml1/2* that showed mildly decreased activity in ΔD2 ESCs. Thus, *Fat1* and *Rex1R* genes utilise only local enhancers within their physically isolated domains in the dismembered TAD for activity in ESCs. As such, during pluripotency, *Fat1* and *Rex1R* genes are functionally independent from one another and the majority of the surrounding ancient regulatory landscape.

### The Rex1/Fat1 TAD is partitioned in ESCs in a CTCF and cohesin-independent manner

We further searched for the mechanism equipping the ancient conserved TAD with such structural flexibility in ESCs. The current prevailing model is that TADs are formed by cohesin progressively extruding chromatin loops until blocked at CTCF boundaries (Fudenberg et al., 2016; Sanborn et al., 2015). As previously reported, binding sites for CTCF and the cohesin subunit Rad21 are enriched within *Rex1R* specifically in ESCs (Figure 3A-B and S4A) (Bonev et al., 2017). From this we speculated that ESC-specific CTCF binding in *Rex1R* blocks cohesin extrusion inside the center of the domain, thereby driving locus restructuring.

To test this, we employed available AID degron-tagged ESCs to globally deplete CTCF or Rad21 (Figures S4F and G) (see STAR methods) (Liu et al., 2021; Nora et al., 2017). Similar to previous reports, cHiC demonstrated most surrounding TAD structures and insulation collapsed once loop extrusion was either unconstrained (dCTCF) or eliminated entirely (dRad21) (Figures 3E and F) (Liu et al., 2021; Nora et al., 2017; Rao et al., 2017). However, surprisingly, in dCTCF ESCs, the *Rex1/Fat1* locus continues to partition into four discrete domains despite loop extrusion now proceeding across *Rex1R* unimpeded. Likewise, the four-domained structure continued to persist after complete ablation of loop extrusion following cohesin-depletion. Therefore, *Rex1/Fat1* TAD partitioning in ESCs occurs independently of CTCF and loop extrusion and must instead be driven by one or several other dominant forces.

### Compartmentalisation dominates in ESCs to partition the Rex1/Fat1 TAD

Beyond loop extrusion, chromatin is also antagonistically structured by the tendency of active or repressed chromatin to physically separate into mutually-exclusive A and B compartments, respectively (Nuebler et al., 2018). Many B compartments then further interact with the NE to form repressive LADs (Falk et al., 2019; Rao et al., 2014; Robson et al., 2017). As the *Rex1/Fat1* TAD restructures into active and inactive domains independently of cohesin, we reasoned that altered compartmentalisation at the NE could drive its partitioning in ESCs.

To examine this possibility, we comprehensively mapped ESC compartments by HiC and corresponding NE-attachment by DamID-seq (Figure 4A) (Vogel et al., 2007). To further directly link altered 3D structure and NE-attachment simultaneously at single loci, we additionally applied polymer modelling and 3D-structured illumination microscopy (3D-SIM) (see Figure S5 and STAR methods for summary) (Barbieri et al., 2012; Beliveau et al., 2015; Gustafsson et al., 2008; Nicodemi and Prisco, 2009; Schermelleh et al., 2008; Szabo et al., 2020; Szabo et al., 2018). For the latter, chromatin was visualised through Oligopaint fluorescence *in situ* hybridisation (FISH) and the NE through Lamin B1 immunolabeling (Figure S5D). Through this modelling and microscopy, we successfully measured simulated and observed structural features, including object NE-proximity, intermingling, and geometric shape (sphericity) (Figures 4C and S6). In all cases, trends extracted from modelling and microscopy closely overlapped and so will be described below interchangeably. However, FISH and modelling measurements can be viewed together simultaneously for comparison in Figures S6.

**Figure 4.**
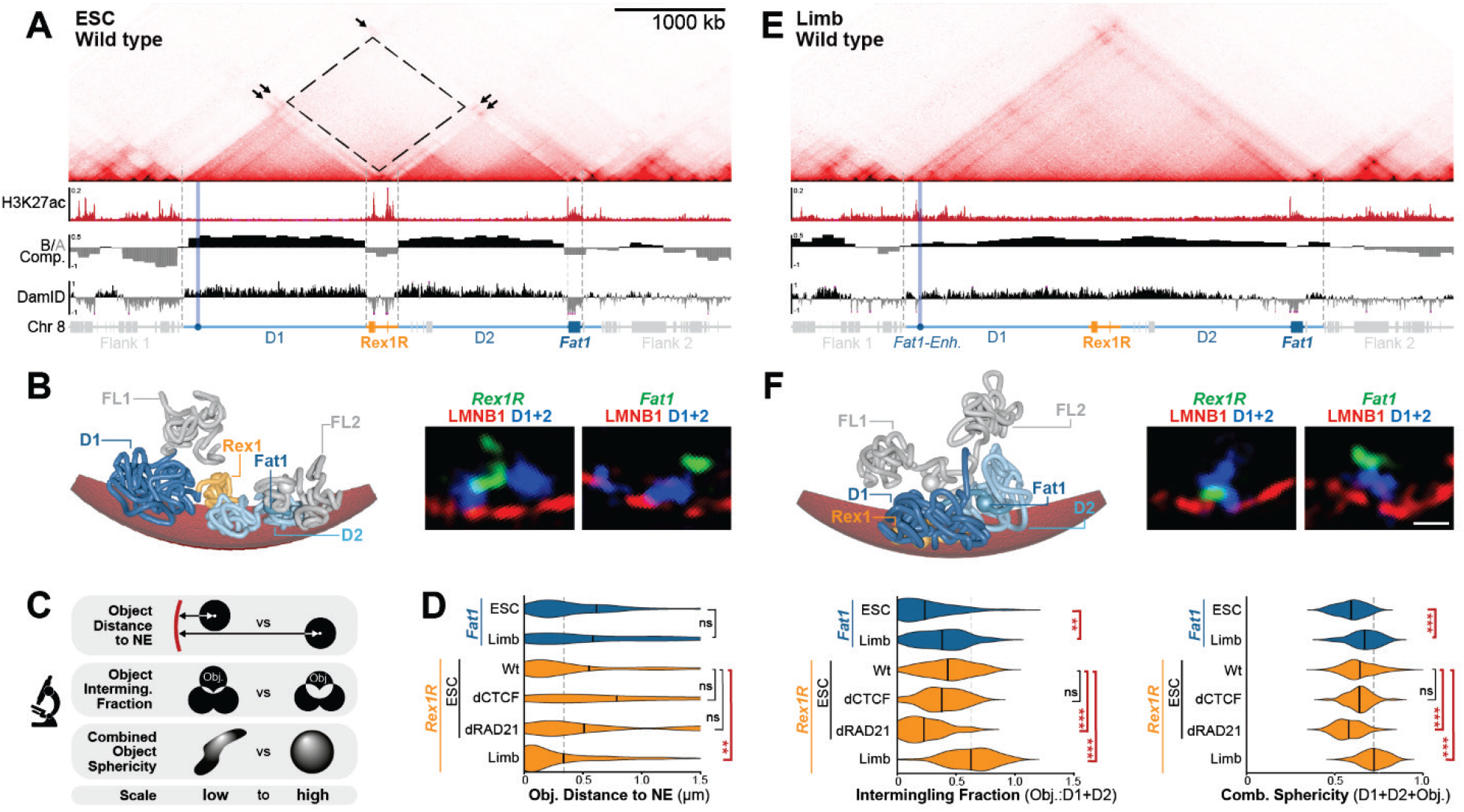
The *Rex1/Fat1* TAD is restructured into discrete compartments in ESCs but accommodates different chromatin environments in limb. A. cHiC from ESCs with H3K27ac-ChIP-seq, compartments and Lamin B1 DamID tracks below. B. *Left*. Modified strings-and-binders polymer model in ESCs with simulated NE (red). *Right*. Representative Z-slice of Lamin B1 immunostaining (red) with Oligopaint-stained D1+D2 (blue) and *Rex1R* or *Fat1* (green) in ESCs. Scale bar is 500 nm. **C.** Summary of FISH measurements for object (i) centroid distance to the NE, (ii) fraction of intermingling with D1+D2, and (iii) combined sphericity of D1+D2 with *Fat1* or *Rex1R*, respectively. **D.** Quantification of FISH measurements in wildtype, CTCF-depleted (dCTCF) and Rad21-depleted (dRad21) ESCs or wildtype limb. Grey line highlights median limb values for reference. **E and F.** cHiC and matching genome browser views (E), polymer modelling (F, left) and FISH images (F, right) in E11.5 limbs. *** p<0.001, * p<0.05 and ns p>0.001 from Welch’s *t*-test comparisons between indicated samples. FISH; n=16 -138 alleles of at least two biological replicates. See Figures S5 and S6 for summaries of modelling optimisation or its quantitative comparison with FISH, respectively.

Collectively, this demonstrated TAD partitioning in ESCs directly corresponds with underlying compartmentalisation and NE-attachment. Specifically, in ESC cHiC, active *Fat1* and *Rex1R* both occupy separated A compartments with their proximal enhancers while D1 and D2 represent B compartments (Figure 4A). Matching this, *Rex1R* and *Fat1* each possess low NE-proximity and intermingle poorly with D1+D2 (Figures 4B-D and S6A-B). Conversely, D1 and D2 themselves remain NE-attached and also poorly intermingle together (Figures S6A and D). As a result, in ESCs, collective *Rex1R+*D1+D2 or *Fat1+D1+D2* sphericity remains low, thereby indicating the objects exist as separated structures in a non-spherical elongated state (Figures 4B-D and S6C). Similar local NE-disassociation of *Fat1* and *Rex1* also occurs in human ESCs (Figure S4C and D) (van Schaik et al., 2020). Thus, *Fat1* and *Rex1R* genes occupy separate compartments away from the NE that match their independent utilisation of local enhancers.

We thus aimed to confirm that compartmentalisation is the dominant force driving TAD partitioning by applying immunoFISH to CTCF- and Rad21-depleted ESCs. Significantly, *Rex1R* intermingling and combined sphericity with D1+D2 were unaffected when loop extrusion proceeds unrestricted in CTCF-depleted ESCs (Figure 4D). However, both measurements decrease following cohesin depletion, thereby indicating the region’s partitioning into compartments further intensifies when loop extrusion is eliminated. *Rex1R’*s NE-proximity was not affected following either depletion (Figure 4D). Hence, in ESCs compartmentalisation overrides loop extrusion to disassemble the TAD. Supporting this, deleting the active *Rex1R* compartment in ΔRex1R ESCs restored D1:D2 intermingling to partially reassemble the TAD (Figure S6E and F). However, *Fat1’*s continued association in a separate active compartment maintains its separation from D1 and D2 in this partially restored TAD structure.

Collectively, this demonstrates compartmentalisation restructures the TAD in ESCs to isolate *Rex1R* genes and *Fat1* with separate local enhancer clusters. Consequently, *Rex1R* and *Fat1* operate as independent entities when simultaneously active within their shared regulatory landscape in placental mammals.

### LADs neither directly silence nor indirectly insulate Rex1R genes

We now sought to dissect the later embryonic limb situation where *Rex1R* genes remain inactive despite contacting *Fat1* and its distal limb enhancers within an intact TAD. We thus repeated our compartment, DamID, modelling and FISH analyses in the limb. In limbs the inactive *Rex1R* is now incorporated with D1 and D2 in a large B compartment LAD that spans most of the intact TAD, as reported in other differentiated cell types (Figure 4E) (Takebayashi et al., 2012). By contrast, the active *Fat1* still locates within an A compartment and, together with its limb *Fat1-enh,* remains locally detached from the NE. Modelling and FISH demonstrate this differential NE-attachment confers *Fat1* and *Rex1R* with distinct positions within the intact TAD (Figure 4F). *Rex1R* now displays higher intermingling with D1+D2 and is buried on average within 330 nm of the repressive NE (Figures 4D and S6A-B). By contrast, reduced D1-D2 intermingling and NE-proximity preferentially positions *Fat1* at the TAD’s nucleoplasmic surface. Thus, unlike in ESCs, the intact limb TAD simultaneously supports multiple inactive LAD and active non-LAD compartments. Accordingly, active *Fat1* transcription is driven preferentially at the intact TAD’s nucleoplasmic face by locally detaching distal enhancers despite intervening inactive chromatin remaining NE-associated (Figure 4E-F).

LADs are compacted heterochromatin domains that frequently repress transcription (Leemans et al., 2019; Ou et al., 2017; Robson et al., 2016). Accordingly, we reasoned that the LAD environment surrounding *Rex1R* inactivates its genes in limb by direct repression or by indirectly blocking *Fat1* enhancer activities. To discriminate these possibilities, we (i) eliminated surrounding LADs or (ii) mapped the enhancer activities received at *Rex1R’*s NE-attached position.

We first tested the effects of eliminating the LADs surrounding *Rex1R*. Examining ΔD1, ΔD2 or ΔD1+2 mutant limbs by cHiC revealed *Rex1R* and *Fat1* continue to co-occupy an increasingly reduced but still insulated TAD once surrounding LADs are removed (Figure 5A and Figure S7A-D). However, FISH measurements demonstrate this significantly alters *Rex1R’*s nuclear environment. *Rex1R* displays reduced NE-attachment in combined ΔD1+2 LAD deletion-limbs, but not their single ΔD1 or ΔD2 counterparts (Figure 5B). This released *Rex1R* in ΔD1+2 limbs now increases intermingling with flanking chromatin (Flank 1) lying outside the TAD (Figure 5B). Thus, the surrounding D1 and D2 LADs together stabilise *Rex1R* at the NE and contributes to physically blocking its interactions with chromatin outside the TAD. Nevertheless, despite this greater association with active chromatin, *Rex1R* genes are not ectopically activated in ΔD1+2 mutant limbs (Figure 3D). As such, NE-attachment can be uncoupled from transcriptional activity and is not necessary to directly drive or maintain *Rex1R* gene silencing.

**Figure 5.**
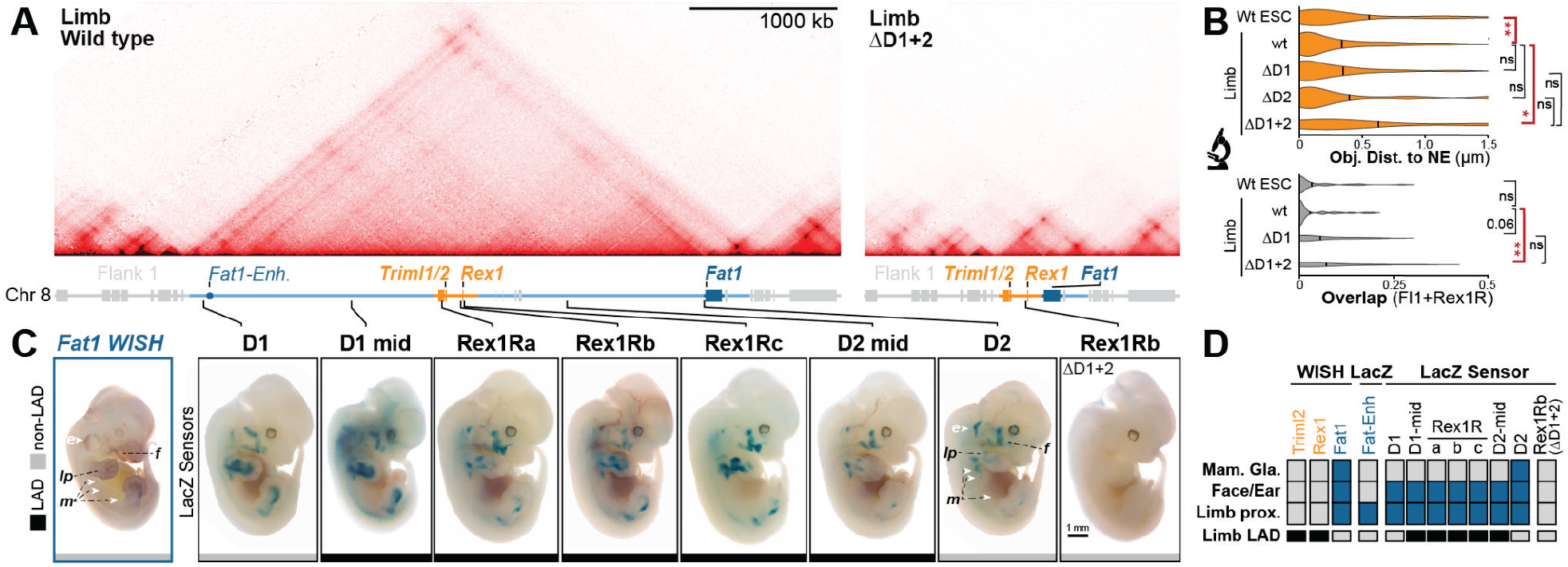
Surrounding LADs stabilise *Rex1R’*s NE-attachment but do not silence its genes or block their communication with *Fat1* enhancers. A. HiC from wildtype and ΔD1+2 E11.5 limb buds. **B.** FISH quantification of *Rex1R-*distances to NE and *Rex1R-*overlap with neighbouring Flank 1 (Fl1) chromatin. In limb buds, *Rex1R* requires adjacent LADs for consistent NE-attachment and isolation from flanking chromatin. *** p<0.001, ** p<0.01, * p<0.05 and ns p>0.05 from Welch’s *t*-test comparisons between indicated samples. WISH; n= 28 - 44 alleles of at least two biological replicates. **C.** Staining of endogenous *Fat1* (WISH, *left*) or integrated *β-globin* LacZ sensors (LacZ, *right*) in E12.5 embryos. Sensor integration sites are indicated by lines and their NE-attachment in limb by black (LAD) or grey (non-LAD) boxes. Staining is indicated in the ear (e), mammary glands (m), face (f), and proximal limb (pl). See Figure S7E for additional wildtype *Rex1* and *Triml2* WISH. **D.** Summary of gene, enhancer and sensor activities with LAD-status indicated. Promoters activate and sample *Fat1* information found within D1 and D2 regardless of lamina-association or proximity to *Rex1R*. See Figure S7.

Accordingly, we next determined if surrounding heterochromatic LADs instead indirectly facilitate *Rex1R* gene inactivity by blocking their communication with *Fat1* enhancers. Hence, we mapped the availability of *Fat1* regulatory activity by integrating minimal promoter-LacZ reporter constructs at seven positions throughout the TAD (Symmons et al., 2014). LacZ staining of E12.5 embryos revealed all such “sensor” locations within the TAD recapitulated the *Fat1* expression pattern, though subtle positional differences were observed (Figure 5C-D). For example, E12.5 mammary gland-staining was only observed near the *Fat1* promoter while proximal limb-signal proportionally increased at positions closer to the limb *Fat1-enh*. Nevertheless, *Fat1-*like ear, face and proximal limb LacZ staining was observed in E12.5 embryos at all three *Rex1R* sensor positions lying within 3-20 kb of the *Rex1* or *Triml1*/*2* promoters. Moreover, sensor staining from the *Rex1Rb* position was absent when integrated in ΔD1+2 embryos where the majority of the TAD and *Fat1* embryonic enhancers are eliminated (Figure 5C-D). Thus, the genomic positions of *Rex1R* genes successfully sample *Fat1* enhancers despite extensive surrounding and intervening heterochromatic LADs.

Collectively, this demonstrates that LADs neither directly silence *Rex1R* genes nor indirectly block their communication with *Fat1* enhancers.

### Enhancer-promoter specificity is not responsible for Rex1R gene inactivity

As regulatory information is sampled throughout the intact limb TAD, we postulated that strict functional incompatibility of *Rex1R* promoters with *Fat1* enhancers maintains their later embryonic inactivity (van Arensbergen et al., 2014). We therefore exchanged the *Rex1*, *Triml1/2* or *Fat1* core promoters into the LacZ regulatory sensor and positioned these constructs at Rex1Rb, 20 kb from the endogenous *Rex1* promoter (Figure 6A). Moreover, as a control, these modified sensor constructs were first integrated at the *Rosa26* safe harbour locus to confirm their lack of autonomous, enhancer-independent transcription (Figure S7F). In all cases, no LacZ signal was observed at the enhancer-free *Rosa26* locus (Figure S6G). By contrast, the *Triml1/2*, *Rex1* and *Fat1* promoters integrated at Rex1Rb all recapitulated the *Fat1-*like limb, face and ear LacZ activity pattern observed with the previous *β-globin* sensor (Figure 5A). Thus, remarkably, *Rex1R* and *Fat1* promoters are fully compatible with active *Fat1* enhancers in the TAD in later embryos. Nevertheless, the *Fat1* promoter generated additional *Fat1-*expression domains, including the forebrain and limb apical ectodermal ridge (AER), thereby indicating some degree of selectivity exists (Figure 5A). Regardless, these data indicate endogenous promoter silencing, rather than a strict enhancer-promoter compatibility code, likely maintains *Rex1R* gene inactivity in later embryos.

**Figure 6.**
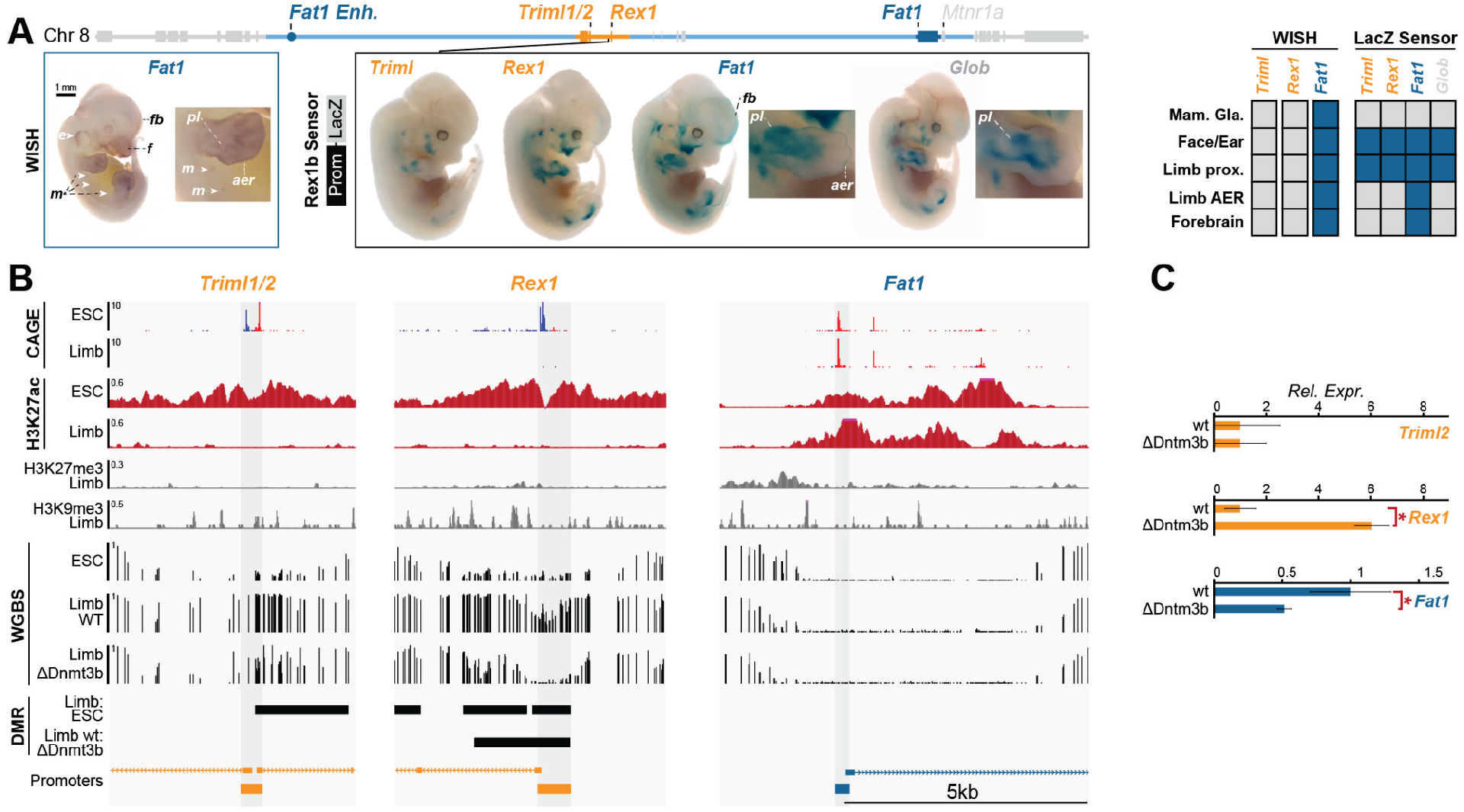
DNA methylation and not enhancer compatibility renders *Rex1* insensitive to *Fat1* regulatory information. **A.** E12.5 embryos stained for *Fat1* WISH (*left*) or LacZ sensors (*middle)* driven at Rex1Rb by the *Triml1/2*, *Rex1, Fat1* or *β-globin* (*Glob*) core promoters with summary (*right*). Staining is indicated in the ear (e), mammary glands (m), face (face), forebrain (fb), proximal limb (pl) and apical ectodermal ridge (aer). In all cases, core promoters were selected to incorporate at least 250 bp upstream and 50 bp downstream of the major endogenous TSS-defined in FANTOM5 CAGE transcriptomes (see panel B). **B.** CAGE, H3K27ac, H3K27me3, H3K9me3 and WGBS tracks from ESCs and/or E11.5 limb buds. Cloned minimal promoters are highlighted in grey. Differentially methylated regions (DMRs) between wildtype and DNMT3B KO limbs are denoted by black bars. DMRs were calculated with a minimum methylation difference of 0.2 containing at minimum 10 CpGs not further than 300 bp apart from each other filtered using a Q-value < 0.05 (2D-KS test, Bonferroni correction). **C.** RNA-seq expression effects of DNMT3B KO on *Triml2, Rex1 and Fat1* expression in wildtype and mutant limbs. RNA-seq was performed in at least biological duplicates. *** p<0.001, * p<0.05 and ns p>0.001. See Figure S7.

### DNA methylation desensitises Rex1 to limb enhancers

We thus sought to determine which repressive mechanisms could drive silencing of the placental-mammal specific *Rex1R* genes in the embryonic limb. Analysis of published ChIP-seq identified no enrichment of H3K27me3 or H3K9me3 at *Rex1R* promoters in E11.5 limbs, thereby ruling out both polycomb and classical heterochromatization as silencing mechanisms (Figure 6B) (Gorkin et al., 2020). By contrast, whole genome bisulfite sequencing (WGBS) identified differentially methylated regions (DMRs) between limb buds and ESCs that surround the *Rex1* and *Triml1*/*2* promoters. Specifically, the DMRs at the *Rex1* or *Triml1/2* promoters go from 13-25% DNA methylation in ESCs to 57-93% methylation in limb buds. Conversely, matching its on-going transcription, the *Fat1* promoter remains permanently unmethylated in both cell types.

Consequently, we reasoned that DNA methylation renders *Rex1R* genes permanently insensitive to ancient *Fat1*-enhancer activities in later embryonic tissues.

We thus generated E11.5 embryos lacking the *de novo* DNA methyltransferase 3B (DNMT3B) (Figure S7H). As reported previously, WGBS in DNMT3B^-/-^ embryonic limbs confirmed a DMR denoting a 71% loss of methylation at the *Rex1*, but not *Triml1/2* or *Fat1* promoters (Figure 6B) (Borgel et al., 2010). Unfortunately, further reductions to ESC methylation-levels in limb were not possible as embryos lacking both DNMT3A and DNMT3B died before E11.5 as previously reported (data not shown) (Okano et al., 1999). Nevertheless, *Rex1* displayed 6-fold upregulation when partially unmethylated in DNMT3B^-/-^ embryonic limbs (Figure 6C). By contrast, *Triml1/2*’s still methylated promoter was unaffected while *Fat1’s* consistently unmethylated promoter displayed ∼50% reduced expression. Collectively, this suggests the endogenous *Rex1* promoter is rendered insensitive to *Fat1* limb enhancers by DNA methylation-driven silencing.

## DISCUSSION

Here, we show that two mechanisms allowed an ancient TAD to incorporate new independently-regulated genes during evolution, namely promoter repression and 3D-restructuring (Figure 7). In later embryonic limbs, DNA methylation renders the eutherian *Rex1* promoter unresponsive to compatible *Fat1* enhancers found in the same TAD. By contrast, in pluripotent stem cells, independent *Fat1* and *Rex1R* regulation is achieved by partitioning the ancient conserved TAD into four discrete compartments. Hence, no single feature explains divergent *Rex1* and *Fat1* expression alone, thereby necessitating our simultaneous analysis of several mechanisms operating in parallel in embryos *in vivo*.

**Figure 7.**
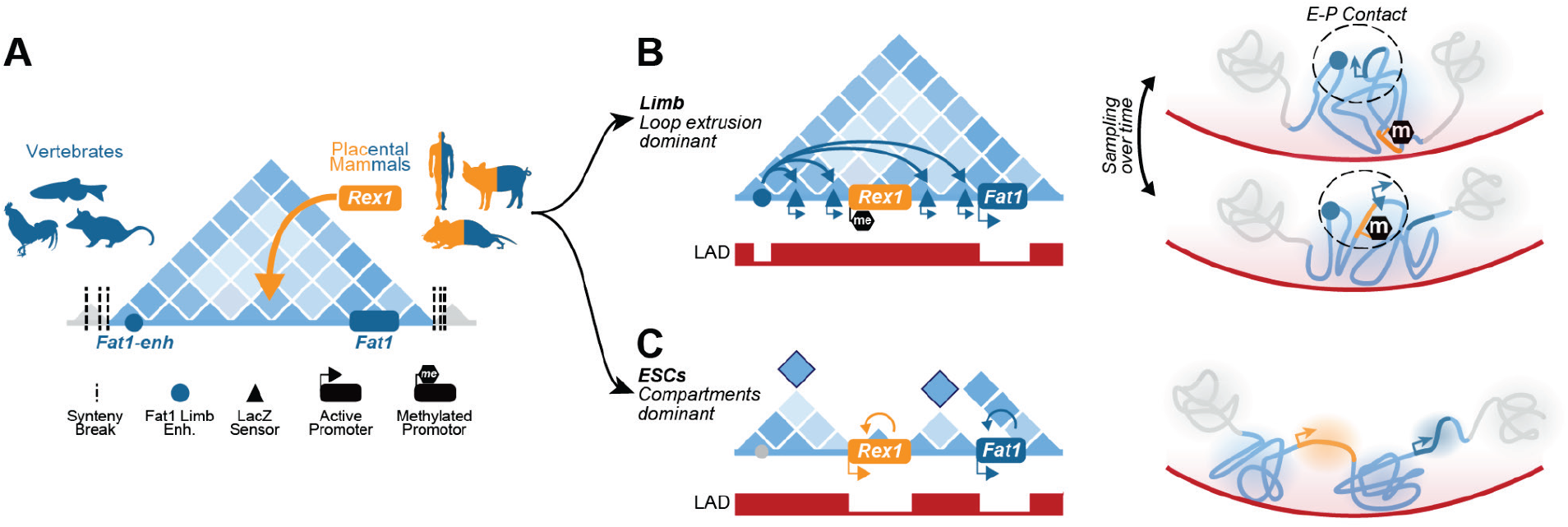
Model for independent *Rex1R* expression within *Fat1*’s ancient TAD. **A.** *Fat1*, its enhancer landscape and TAD existed together as a regulatory unit in all vertebrates despite frequent flanking synteny breaks. *Rex1* and *Triml1/2* emerged with divergent expression within this domain in later placental mammals. **B.** In the limb and embryo, *Fat1* enhancers emerge from LADs and promiscuously sample promoters throughout the domain’s both active and NE-attached inactive compartments. However, despite this and its functional compatibility with *Fat1* enhancers, methylation of *Rex1’*s promoter prevents its activation. **C.** In ESCs, increased compartmentalisation and/or weakened loop extrusion restructures the TAD, thereby driving the *Rex1R* and *Fat1* genes to independently utilise only local enhancers.

TADs are frequently described as stable structural scaffolds that ensure transmission of enhancer activities to promoters found within a domain’s boundaries (Andrey and Mundlos, 2017). In this paradigm, genes with similar functions can be controlled together in a shared TAD while those requiring divergent regulation must be placed alone in separated domains (Wu et al., 2021). However, we and others observe that this framework is too simplistic. We find most multi-gene TADs in the genome display poor average co-regulation of their collective hosted developmental genes. Accordingly, many regulatory landscapes are reported to support multiple independent expression programs within the same genomic region (Andrey et al., 2013; Huang et al., 2017; Palstra et al., 2003; Soshnikova and Duboule, 2009). This capacity furnishes the genome with enormous regulatory complexity and evolutionary flexibility. For example, the divergent spatiotemporal expression of *Hox* promoters from their multi-gene TADs faciliates proper embryo morphogenesis in development (Andrey et al., 2013; Noordermeer et al., 2011). Likewise, we show that the entirely new *Rex1R* genes and their enhancers could emerge without disrupting the preexisting *Fat1* regulatory landscape and its diverse physiological functions. However, perhaps more significantly, the ability to host multiple non-overlapping regulatory programs likely increases the genome’s tolerance for structural rearrangement by preventing gene misexpression. Indeed, many structural variant mutations that combine genes and enhancers in shuffled TADs do not drive gene misexpression or disease (Despang et al., 2019; Laugsch et al., 2019). Consequently, regulatory landscapes must not be viewed as rigid TAD blocks but rather as flexible entities that employ multiple mechanisms to refine how enhancers are utilised. Understanding these mechanisms will be critical when predicting how rearranged regulatory landscapes behave in disease and evolution.

Our results indicate enhancer activities can be further refined by modulating their physical contacts with genes through alterations to chromatin structure and NE-attachment. Similar structural flexibility has previously been observed to modulate enhancer activities at other loci. For example, a largely unknown molecular mechanism drives a topological switch that ensures the *Pen* enhancer only contacts and activates the *Pitx1* promoter in the hindlimb (Kragesteen et al., 2018). Similarly, during erythrogenesis, interactions between the Ldb1 and GATA1 transcription factors drive LCR enhancers to sequentially contact and activate distinct *β-globin* genes within their shared TAD (Deng et al., 2014; Huang et al., 2017; Palstra et al., 2003). Here, TAD structural disassembly isolates *Rex1* and *Fat1* with separate enhancers in ESCs within epigenetically-defined compartments that ensure their independent activation away from the NE. Mechanistically, this disassembly occurs independently from loop extrusion and is instead seemingly a product of compartmentalisation, i.e. the antagonistic tendency of active and inactive chromatin to spatially separate (Rao et al., 2017; Schwarzer et al., 2017). This raises the intriguing possibility that chromatin structure, and thus enhancer-promoter contacts, could be controlled and self-reinforced by their underlying epigenetic state. Supporting this, eliminating H3K27ac-marked enhancers and promoters was reported to collapse compartment-separation at *Rex1* as well as *Dppa2* (Sima et al., 2019). However, more generally, it highlights that TADs can be flexibly re-organised by compartmentalisation and NE-attachment when required despite their apparent stability across cell types and species.

Such NE-dynamics make it important to consider the functional role of LADs in regulating *Rex1* expression within its TAD. Indeed, LADs are generally viewed as repressive entities due to their higher compaction, lower mobility and frequent repression of promoters incorporated within them (Chubb et al., 2002; Finlan et al., 2008; Leemans et al., 2019; Ou et al., 2017; Reddy et al., 2008). However, artificial activation of genes in LADs drives the locally restricted detachment of their promoters and gene bodies from the NE (Brueckner et al., 2020; Therizols et al., 2014). Thus, though LADs may be a barrier to optimal transcription, they can be locally restructured to allow gene escape and activation when needed. Supporting this, we observe the *Fat1-enh* undergoes local NE-detachment when active in limbs. Likewise, our integrated promoter-lacZ reporters are successfully activated in limbs despite their integration in LADs. Moreover, this activation was driven by *Fat1* enhancers despite up to 3.5 Mb of LADs separating them. As such, LADs are not fundamentally incompatible with either gene activation or enhancer-promoter communication. Rather, intact TADs provide an environment to mix active and inactive chromatin through loop extrusion. In this way, TADs can act to facilitate widespread enhancer-promoter communication across the diverse LAD and non-LAD environments of regulatory landscapes.

Strict functional compatibilities between distinct enhancers and promoters have been proposed to instead define how regulatory activities are utilised (van Arensbergen et al., 2014). Indeed, several examples of such strict compatibilities have been observed in drosophila (Li and Noll, 1994; Merli et al., 1996). However, in mammals the few tested developmental promoters faithfully recapitulate surrounding enhancer activities when integrated into ectopic TADs (Marinic et al., 2013; Shima et al., 2016; Symmons et al., 2014). Similarly, we find that the *Triml1/2, Rex1,* and *Fat1* promoters all drove a similar *Fat1-*like expression pattern in later embryos when integrated as LacZ sensors. Thus, at least at this locus, enhancers can promiscuously activate developmental promoters but are prevented from doing so by additional mechanisms. Nevertheless, we acknowledge the significant caveat that lacZ staining does not quantitatively measure transcription. We also observe novel AER and forebrain lacZ expression domains in the sensor employing *Fat1*’s promoter. Thus, though largely functionally compatible, differences in promoter sensitivity to distinct enhancers could drive significant quantitative differences in their overall regulation. Accordingly, though challenging, it will be critical to systematically determine the extent and causes of different promoter responsiveness to enhancers (Long et al., 2016; van Arensbergen et al., 2014). However, at least here, such quantitative differences cannot account for the complete inactivity of *Rex1R* genes observed in embryos.

Promoter repression presents a solution to the problem of enhancer promiscuity within the highly communicative environments of TADs. For example, the sequential release of polycomb repression allows the co-linear activation of *HoxD* genes by enhancers located within their shared TAD (Noordermeer et al., 2011; Soshnikova and Duboule, 2009). Likewise, during differentiation, both X chromosomes activate enhancers within the X-inactivation center TAD in differentiating female cells. However, *Xist* becomes marked by H3K9me3 on one allele, thereby rendering it insensitive to enhancers and enabling its chromosome to escape random X inactivation (Gjaltema et al., 2021). Here, we find DNA methylation provides another mechanism that can control gene susceptibility to enhancers. Importantly, this model likely explains the only minor gene expression defects in development observed when DNA methylation is eliminated entirely in early embryos (Grosswendt et al., 2020; Yagi et al., 2020). In this TAD view, misexpression would be limited to only unmethylated genes exposed to enhancers within shared landscapes and, even then, only in the specific cell types where those enhancers are active. As such, DNA methylation forms part of a larger ecosystem of repressive mechanisms that further refine highly promiscuous enhancer activities. Combined with dynamic changes to chromatin structure, such promoter repression enables divergent transcriptional programs to be encoded within the same overlapping genomic locus in evolution.

From this a more refined view of TADs is emerging. While TADs partition regulatory interactions, other mechanisms govern where and when these interactions activate promoters. Consequently, all levels of regulation - from promoter-state to flexible 3D structure - must be considered to successfully predict the genome’s transcriptional outputs. As such, we believe such cell-type-specific measurements of promoter state and 3D structure should be incorporated into recent enhancer-promoter models (Fulco et al., 2019; Nasser et al., 2021; Zuin et al., 2021). With this, these tools will better predict the benign or pathological effects of structural variant mutations in human patients.

---

**Figure S1.**
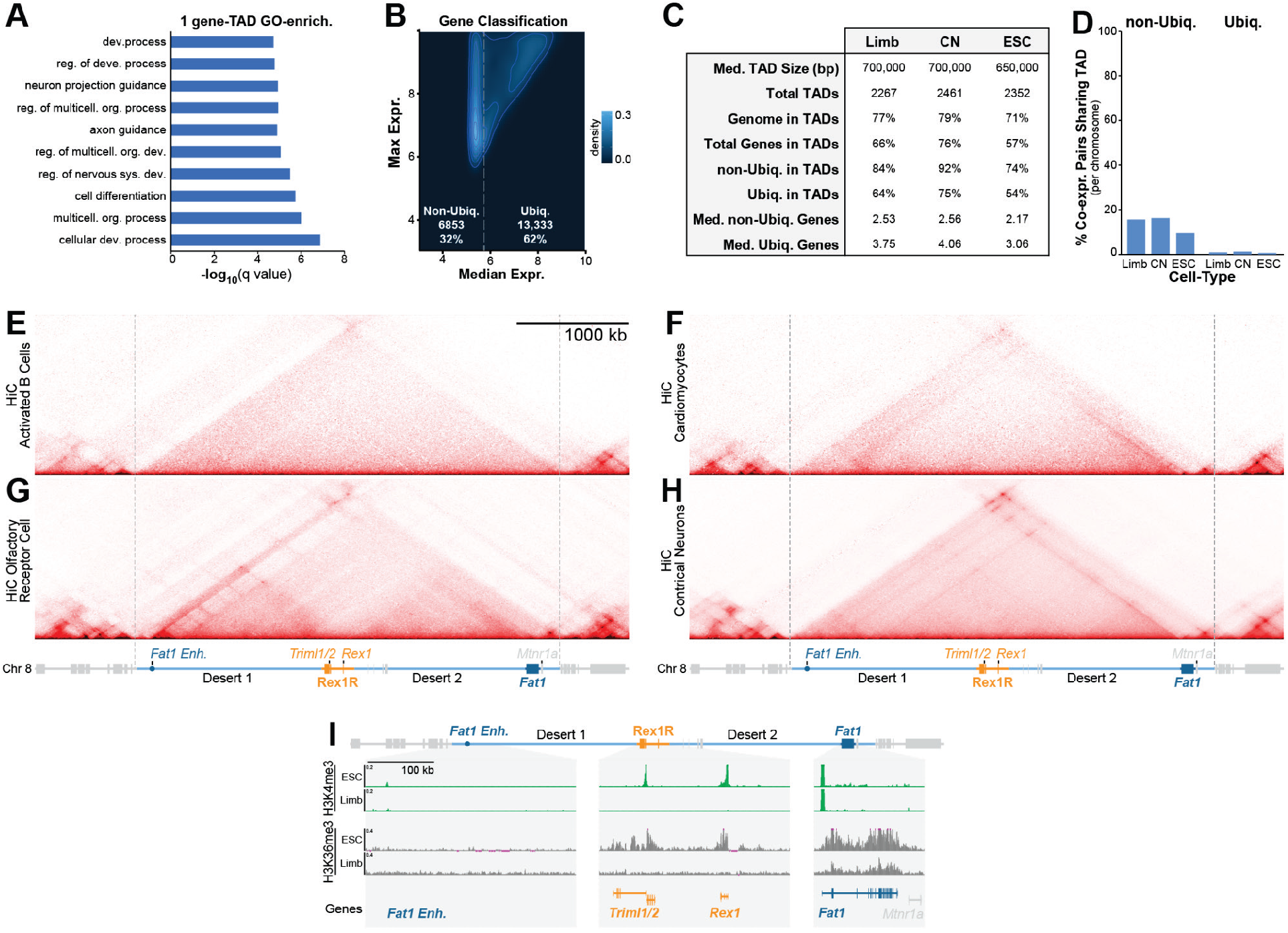
Summaries of co-expression analysis and confirmation of *Rex1/Fat1* TAD maintenance across multiple cell types. **A.** GO-term enrichment for genes within single-gene TADs (Eden et al., 2009). **B.** Classification of genes into non-ubiquitously-(non-Ubiq.) and ubiquitously- (Ubiq.) expressed classes according to their maximum and median expression across FANTOM5 CAGE samples. **C.** TAD and gene statistics in limb, CNs and ESCs. **D.** Fraction of co-expressing gene pairs found on the same chromosome that share TADs. Most co-regulation on each chromosome occurs *in trans* outside of TADs. **E-H.** Published HiC from activated B cells (E), cardiomyocytes (F), olfactory receptor cells (G), and cortical neurons (H) demonstrating TAD boundary stability across multiple lineages. **I.** Zoom of the centromeric TAD arm, *Rex1R* and *Fat1* gene body with H3K4me3 and H3K36me3 ChIP-seq shown. Note that *Triml1* and *Triml2* are transcribed from a single shared bidirectional promoter as indicated by a single peak of H3K4me3 and broad H3K36me3 marking the transcribed gene body. See Figure 1.

**Figure S2.**
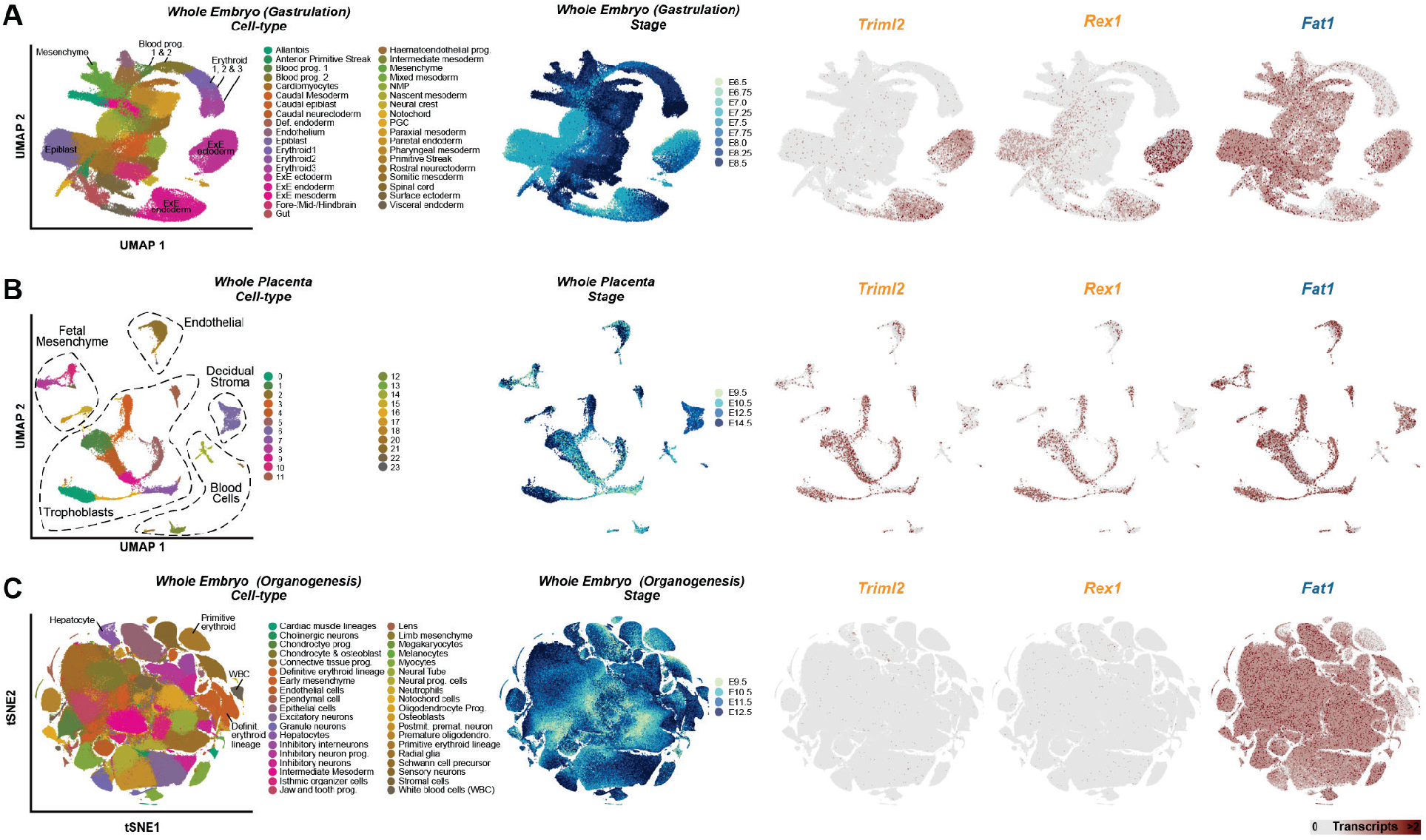
*Rex1R* genes and *Fat1* are independently expressed during gastrulation, organogenesis and placental development. **A-C.** UMAPs from re-processed scRNA-seq from whole gastrulating embryos (A), the developing placenta (B), and whole embryos during organogenesis (C) (Cao et al., 2019; Marsh and Blelloch, 2020; Pijuan-Sala et al., 2019). UMAP embedding is coloured according to cell type (*left*), developmental stage (*middle*), or expression of *Triml2, Rex1* or *Fat1 (right)*. *Rex1R* genes (*Triml2* and *Rex1)* are expressed in the extraembryonic ectoderm and endoderm (A) and placental trophoblasts (B). *Rex1* is also expressed in the E6.5 epiblast (A). *Fat1* is expressed widely in many tissues (A-C) but is absent, for example, in blood progenitors and erythroid cells (A and C).

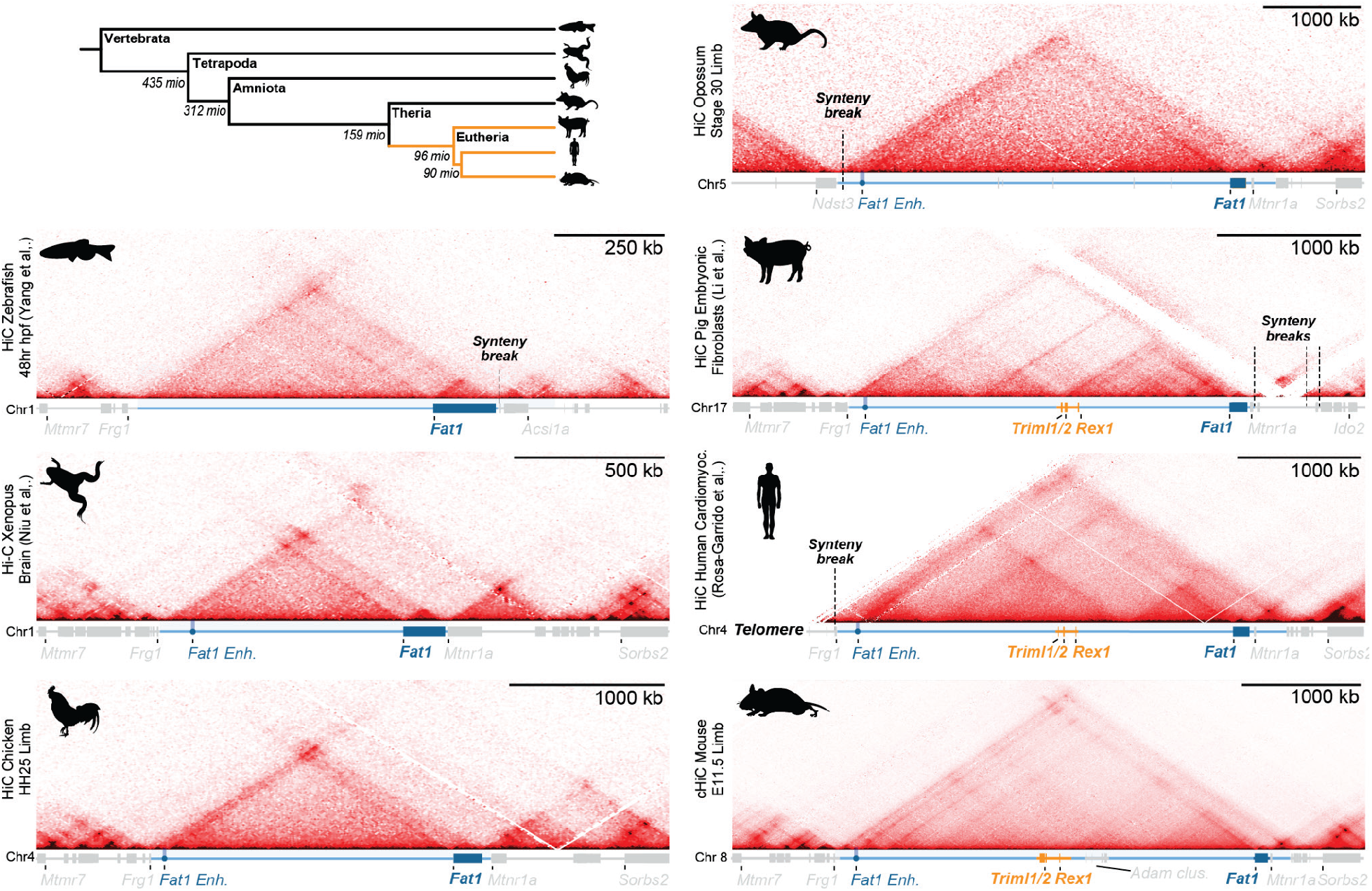

**Figure S4.**
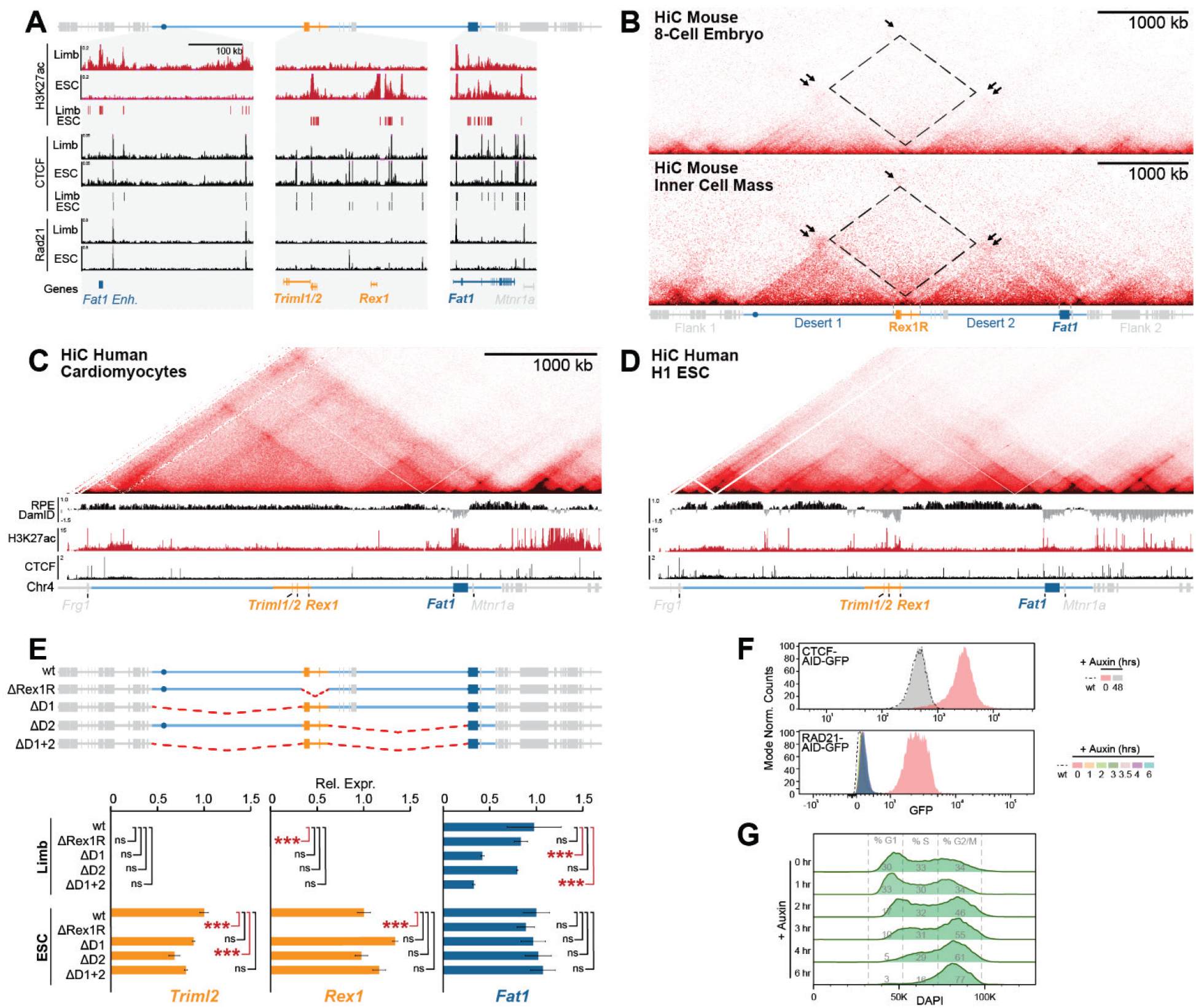
*Rex1*/*Fat1* TAD disassembly is a common feature of pluripotency in placental mammals. **A.** Zooms of E11.5 limb and ESC H3K27ac, CTCF and RAD21 ChIP-seq with called enhancers or CTCF peaks below. **B.** Low input HiC from mouse 8-cell embryos (*top*) and pluripotent cells from the inner cell mass (*bottom*). **C and D.** cHiC from human cardiomyocytes (C) and H1 ESCs (D) with corresponding H3K27ac and CTCF ChIP-seq and DamID shown below. Note DamID from retinal pigment epithelium (RPE) cells was used to define locus lamina-association when *Rex1R* is inactive in differentiated cells. **E.** Schematic of deletion mutants (*top*) with effects on gene expression determined by RNA-seq (*bottom*). n=2-4 biological replicates per sample. *** p<0.001, * p<0.05 and ns p>0.001. **F.** FACs distributions of GFP signal in CTCF-AID-GFP (top) and Rad21-AID-GFP (bottom) ESCs following indicated auxin treatments. **G.** Distribution of cell-cycle phases in Rad21-AID-GFP ESCs showing rapid accumulation in S and G2M within 6 hours. To account for accumulation of Rad21-AID-GFP ESCs in G2/M phase caused by failed sister chromatid cohesion, cHiC was performed on sorted G1 cells 3.5 hours post-auxin addition (Liu et al., 2021). By contrast, due to technical difficulties plating fixed cells on coverslips, FISH was performed on unsorted 2 hour-induced Rad21-AID-GFP ESCs where only moderate shifts in the G1:S:G2/M ratio were observed. See Figures 3 and 4.

**Figure S5.**
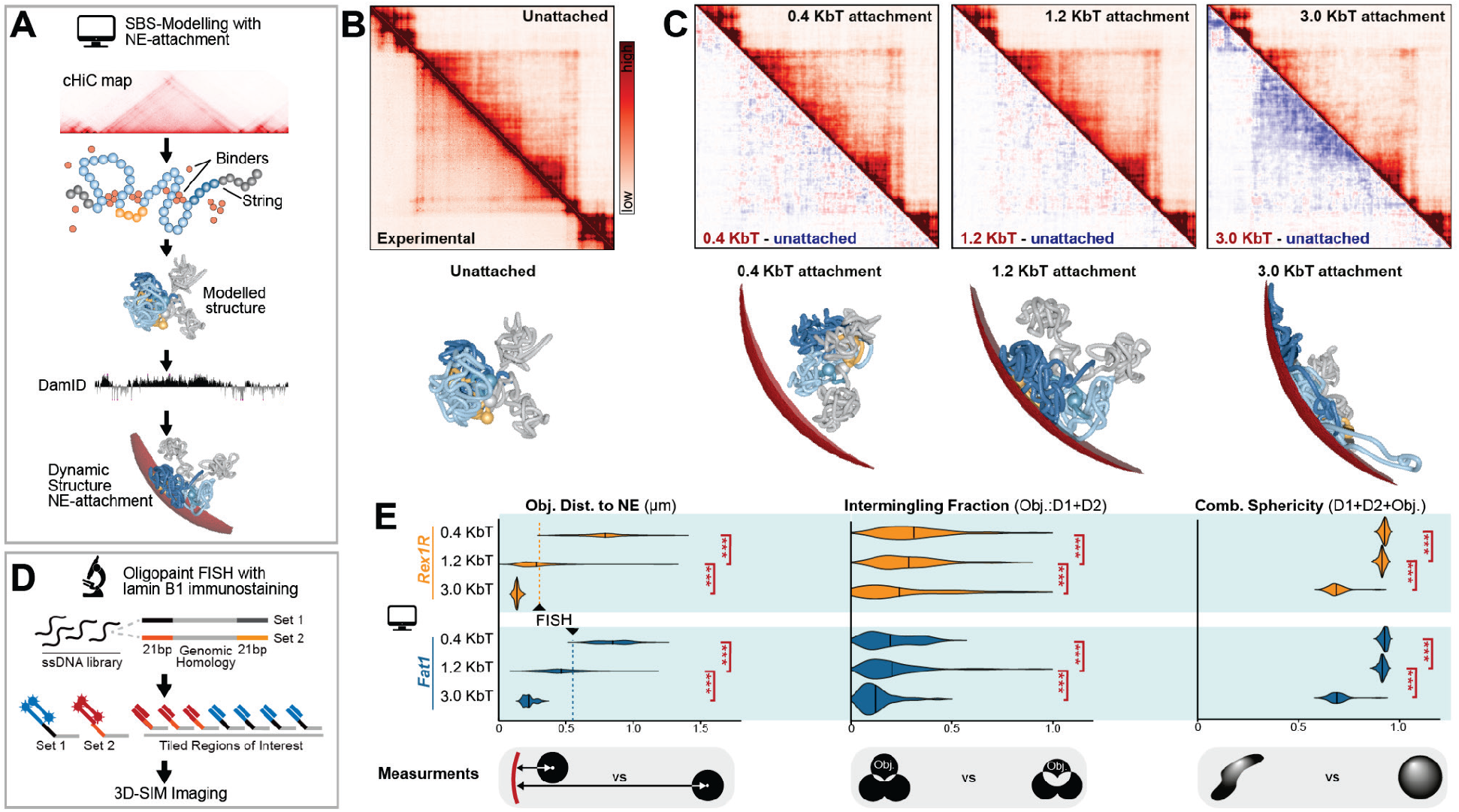
Summary of SBS Modelling with NE-attachment and Oligopaint FISH strategy. **A.** Schematic representation of the modified strings-and-binders (SBS) polymer model. cHiC contact maps were used to define PRISM-assigned chromatin binders. The chromatin polymer is then structured *in silco* through simulated DNA interactions created by the self-association between matching binders (Barbieri et al., 2012; Nicodemi and Prisco, 2009). Generated structures were subsequently dynamically-attached to a modelled NE with polymer affinities determined from sample-matched DamID (see STAR methods). **B and C.** Reconstructed contact maps from simulated limb structures before (B) and after (C) NE attachment with 0.4, 1.2 and 3.0 kTb interaction energies. Corresponding subtraction maps and representative structures are shown below. n=25 - 88 simulations. **D.** Oligopaint FISH 3D-SIM imaging strategy. A library of single stranded DNA oligos with genomic homology and overhangs allow multiplexed staining of multiple regions of interest. **E.** Quantification of object NE-distance (*left*), intermingling fraction (*middle*) and sphericity (*right*) for simulated limb structures following 0.4, 1.2 and 3.0 kTb NE-attachment. 1.2 kTb was selected for further analysis as it produced NE-proximities without deforming the structure’s intermingling or sphericity relative to FISH measurements.

**Figure S6.**
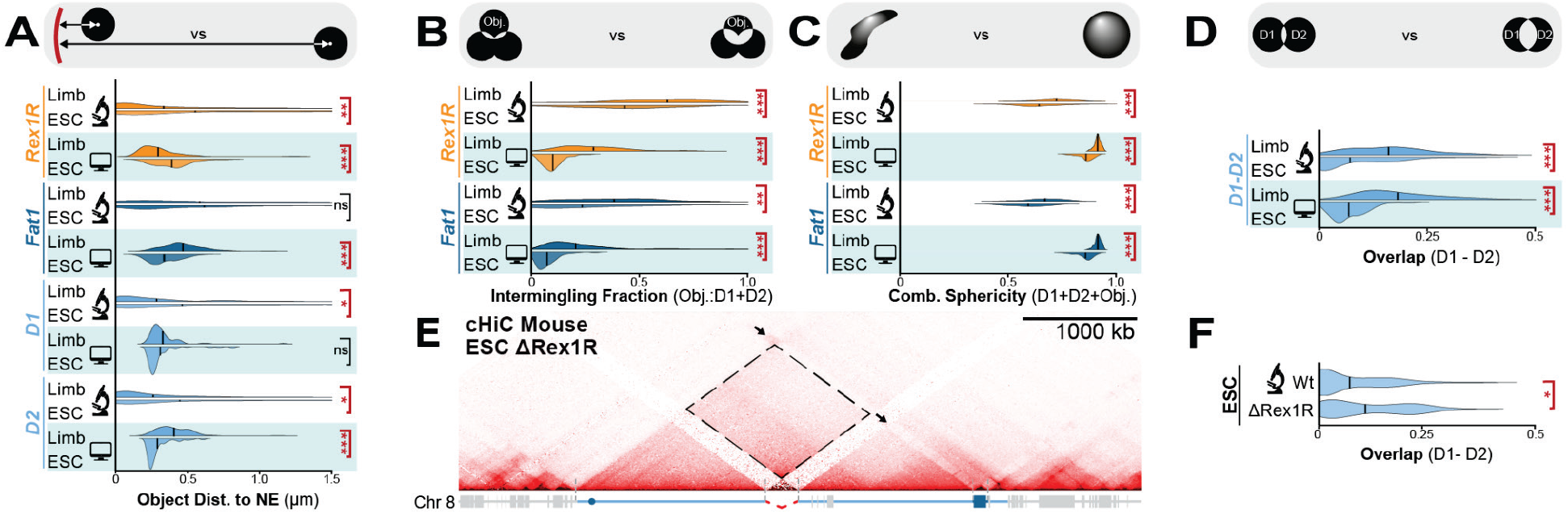
Comparison of simulated and observed *Rex1/Fat1* locus structures. **A-C.** Comparison of simulated 1.2 kTb NE-attachment model and experimental FISH data in wildtype E11.5 limbs and ESCs. Measurements are object NE-distance (A), intermingling fraction (B), and object sphericity with D1+D2 (C). **D.** Comparison of simulated and observed D1 and D2 overlap and combined sphericity in wt ESCs and E11.5 limbs. **E.** cHiC from ΔRex1R ESCs. Arrows indicate *Fat1’*s interaction with active chromatin (*upper*) and avoidance of heterochromatin (*lower*). Dotted rectangle displays gained interaction between inactive D1 and D2. **F.** Quantification of D1 and D2 overlap between wt and ΔRex1R ESCs. *** p<0.001, ** p<0.01, * p<0.05 and ns p>0.05 from Welch’s *t*-test comparisons between indicated samples. FISH; n=28-138 alleles of at least two biological replicates. Modelling; n= 25-106 simulations. See Figure 4.

**Figure S7.**
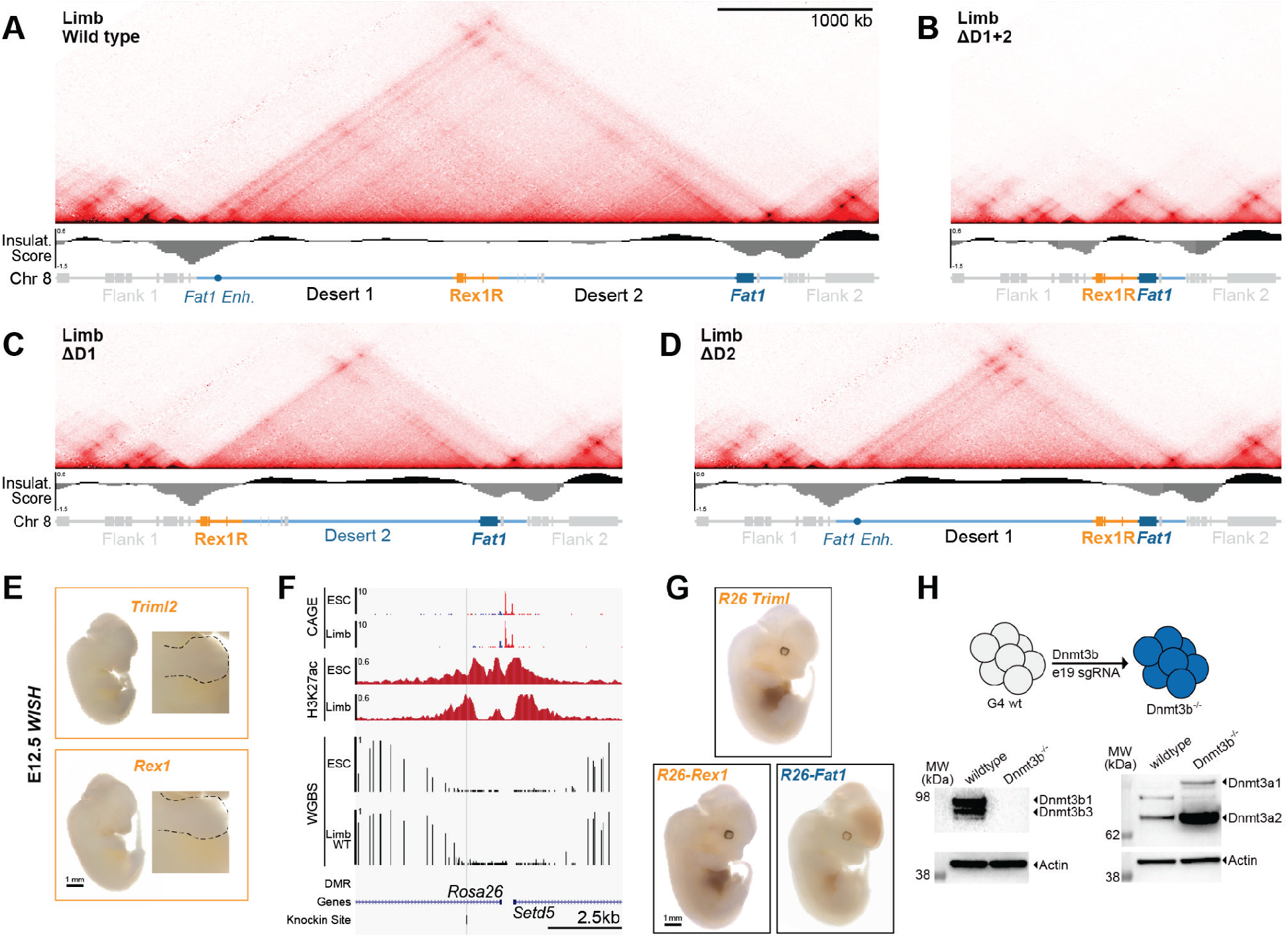
cHiC in limb deletion mutants, testing intrinsic promoter activities and generation of DNMT3A/B knockouts. **A-D.** cHiC with corresponding insulation scores from wt (A), ΔD1+2 (B), ΔD1 (C) and ΔD2 (D) limb buds. Note the *Rex1/Fat1* TAD and its boundaries remain intact even following combined D1 and D2 deletion. **E.** *Triml2* and *Rex1* WISH stainings in E12.5 embryos. **F.** Genome browser view of the *Rosa26* safe harbour locus with CAGE, H3K27ac ChIPseq and WGBS shown. Sensor integration site is indicated by the grey bar. **G.** Example lacZ stainings from E12.5 embryos with sensors with indicated promoters integrated at *Rosa26.* **H.** Strategy for *Dnmt3b* knockout in ESC clones with western blot confirmation shown below. DNMT3A increases following loss of DNMT3B.

## STAR★METHODS

Detailed methods are provided in the online version of this paper and include the following:

- KEY RESOURCES TABLE
- RESOURCE AVAILABILITY
- EXPERIMENTAL MODEL AND SUBJECT DETAILS
- METHOD DETAILS

- Plasmid Construction
- CRISPR-mediated genome editing
- Enhancer Reporter Line Generation
- Auxin induced CTCF and Rad21 depletion
- Western Blot
- Tetraploid morula complementation
- Whole mount in situ hybridisation
- LacZ staining in embryos
- RNA-seq
- Sample collection for DamID-seq, ChIP-seq, ATAC-seq, cHiC and FISH
- DamID-seq
- ATAC-seq
- ChIP-seq
- ChIPmentation
- WGBS
- Capture HiC
- HiC
- Oligopaint fluorescence in situ hybridisation with 3D-SIM imaging
- QUANTIFICATION AND STATISTICAL ANALYSIS

RNA-seq differential expression analysis
Single cell RNA-seq
DamID-seq analysis
ATAC-seq analysis
ChIP-seq analysis
Enhancer prediction
Enhancer conservation analysis
cHiC and HiC analysis
Gene co-regulation in TADs analysis
WGBS processing
Differentially methylated region (DMR) calling
SBS-polymer modelling with NE-attachment
OligoPaint FISH image analyses
Statistical methods

## ACKNOWLEDGEMENTS

We thank Guillaume Andrey and the entire Mundlos lab for input and discussions; Elphege Nora for kindly providing Rad21- and CTCF-AID-GFP ESCs; Katja Zill for mouse line maintenance; Ute Fischer and Asita Steige for genotyping and cloning assistance; Carola Dietrich for HiC sample processing; Norbert Brieske for WISH assistance; J. Fiedler and K. Macura (MPIMG transgenic unit) for aggregations and Myriam Hochradel for cHiC processing (MPIMG sequencing core facility). We thank the Montpellier Ressources Imagerie facility (BioCampus Montpellier, Centre National de la Recherche Scientifique (CNRS), INSERM, University of Montpellier). This work was funded by EMBO and the Wellcome Trust (ALTF1554-2016 and 206475/Z/17/Z; to M.I.R) as well as the Deutsche Forschungsgemeinschaft (KR3985/7-3 and MU 880/16-1 to S.M.).

## AUTHOR CONTRIBUTIONS

M.I.R conceived the study with input from S.M.; M.I.R, A.R.R., K.C. and P.R. produced and validated transgenics; Q.S. and F.B. designed Oligopaint probes, A.R.R., Q.S., F.B. performed FISH and image acquisition in the lab of G.C.; Q.S. performed FISH image analysis; A.M.C., S.B. and A.E. performed SBS-modelling in the lab of M.N., M.I.R., A.R.R. and D.M.I. performed cHiC and HiC which R.S. processed; M.I.R., A.R.R. and V.L. performed and processed DamID-seq; A.R.R., M.S and M.P performed ChIP-seq; A.R.R. and C.P. performed ATAC-seq; A.R.R. performed WISH; M.I.R. and A.R.R. performed and processed RNA-seq, M.I.R. processed ChIP-seq and ATAC-seq data; D.H. performed gene co-regulation analyses; A.L.M. and S.H. generated and processed DNMT3B knockout lines and WGBS data in the lab of A.M.; T.Z. performed evolutionary analysis; L.W. managed aggregations; C.A.P.-M. replotted scRNA-seq data; P.G. provided opossum embryos. M.I.R and A.R.R. wrote the manuscript with input from all authors.

## SUPPLEMENTAL INFORMATION

Supplemental Information includes 7 figures and 5 tables.

## STAR METHODS

### KEY RESOURCES TABLE

| REAGENT or RESOURCE | SOURCE | IDENTIFIER |
| --- | --- | --- |
| Antibodies |  |  |
| rabbit anti-H3K27ac | Diagenode | C15410174 |
| rabbit anti-H3K4me1 | Diagenode | C15410037 |
| rabbit anti-H3K27me3 | Merck Millipore | 07-449 |
| rabbit anti-H3K4me3 | Merck Millipore | 07-473 |
| rabbit anti-lamin B1 | Abcam | ab160486 |
| donkey anti-rabbit IgG-Atto647 | Sigma | 40839 |
| Anti-DIG-AP, 150 U | Roche Diagnostics | 11093274910 |
| rabbit anti-DNMT3A | Abcam | ab188470 lot GR224165-2 |
| rabbit antiDNMT3B | Cell Signaling | cs48488, lot 1 |
| Bacterial strains |  |  |
| One Shot TOP 10 Chemically Competent Cells E.c. | Thermo Fisher | C404006 |
| Chemicals, Peptides, and Recombinant Proteins |  |  |
| Advantage cDNA polymerase | Clontech | 639105 |
| Agencourt AMPure XP magnetic beads | Beckman Coulter | A63880 |
| Auxin | Abcam | ab14642 |
| BCIP , 3ml (150 mg) | Roche Diagnostics | 11383221001 |
| Biotin-14-dATP-50 nmol | Thermo Fisher Scientific | 19524016 |
| BM-Purple , AP-Substrat | Roche Diagnostics | 11442074001 |
| cOmplete, Mini, EDTA-free Protease Inhibitor Coctail | SIGMA-ALDRICH | 4693159001 |
| Covaris micro TUBE AFA Fiber Pre-Slit Snap-Cap tubes | Covaris | SKU - 520045 |
| DMEM, high glucose, no glutamine | Thermo Fisher | 11960085 |
| DNA Pol. Large Fragm. (Klenow) | New England Biolabs | M0210L |
| Dnase,recombinant,RNase-free (10000 U) | Roche Diagnostics | 4716728001 |
| DpnI, recombinant | New England Biolabs | R0176L |
| DpnII, recombinant | New England Biolabs | R0543S |
| Dynabeads MyOne Streptavidin T1-10 mL | Thermo Fisher Scientific | 65602 |
| ESGRO(LIF) | Millipore | ESG1107 |
| Formamide deionized for molecular biology | PanReac AppliChem | APP A2156,1000 |
| FuGENE® HD Transfection Reagent | Promega | E2311 |
| Gelatin 2% solution from bovine skin cell | Sigma Aldrich | G1393 |
| Heparin sodium salt | SIGMA-ALDRICH | H3149 |
| Hygromycin B (50mg/ml) | Thermo Fisher | 10687010 |
| Cot-1 DNA | Invitrogen Life Technologies | 18440-016 |
| Knockout DMEM-500 ml | Thermo Fisher | 10829018 |
| L-glutamine (200mM) | Lonza | 882027-12 |
| Lent-X concentrator | Takara | 631232 |
| Library Efficiency DH5a Competent Cells | Thermo Fisher | 18263012 |
| Lipofectamine 2000 Transfection Reagent | Thermo Fisher Scientific | 11668019 |
| MEM Non Essential Amino Acids Solution | Thermo Fisher | 11140068 |
| NBT, 3ml (300 mg) | Roche Diagnostics | 11383213001 |
| NEBNext® High-Fidelity 2X PCR Master Mix | New England Biolabs | M0541S |
| NEBNext® Multiplex Oligos for Illumina | New England Biolabs | E7335, E7500 |
| NEBNext® Quick Ligation Reaction Buffer (5X) | New England Biolabs | B6058S |
| NEBNext® Ultra II Q5® Master Mix | New England Biolabs | M0544L |
| Opti-MEM I Reduced Serum Medium, GlutaMAX Supplement | Thermo Fisher | 51985026 |
| Penicillin/Streptomycin | Fisher Bioreagents | 10003927 |
| Proteinase K | Roche Diagnostics | 1000144 |
| Puromycin | SIGMA-ALDRICH | P8833 |
| Recombinant Human/Mouse FGF-8b Isoform | R&D Systems | #423-F8-025/CF |
| Recombinant Mouse Wnt-3a protein | R&D Systems | #1324-WN-010/CF |
| Ribonuclease A from bovine pancreas, Type 1-A, RNase A | SIGMA | R4875 |
| Rnase Inhibitor (2000 U) | Roche Diagnostics | 3335399001 |
| Roti-Phenol/ Chloroform/ Isoamylalcohol | Carl Roth | A156.2 |
| SP6-RNA Polymerase ( 1000 U) | Roche Diagnostics | 10810274001 |
| SYBR Green I | Thermo Fisher | S7563 |
| T4 DNA Ligase | New England Biolabs | M0202L |
| T4 DNA Polymerase | New England Biolabs | M0203L |
| T4 Polynucleotide Kinase NK | New England BioLabs | M0201 |
| T7-RNA Polymerase ( 1000 U) | Roche Diagnostics | 10881767001 |
| Tagment DNA Buffer | Illumnia | 15027866 |
| Tagment DNA Enzyme 1 (TDE1) | Illumnia | 15027865 |
| tRNA from Baker's Yeast | SIGMA | R6750 |
| Trypsin-EDTA (0.05%), phenol red | Thermo Fisher | 25300096 |
| Water for Injection (WFI) for cell culture | Thermo Fisher | A1287303 |
| X-beta-Gal min 99 %, BioScience-Grade | Carl Roth | 2315.3 |
| Critical commercial kits |  |  |
| 0.45 µm2 low protein-binding PES syringe filter | Millipore | SLHP003RS |
| Accel-NGS Methyl-seq DNA library kit | Zymo | DL-ILMMS-12 |
| Agencourt AMPure XP beads | Beckman Coulter | A63881 |
| Dig-RNA-labeling Mix | Roche Diagnostics | 11277073910 |
| DNA Clean & Concentrator-5 kit | Zymo | D4013 |
| Dneasy Blood & Tissue Kit(50) | QIAGEN | 69504 |
| EZ DNA Methylation-Gold Kit | Zymo | D5005 |
| FISH Tag DNA Kit | Invitrogen Life Technologies | F32951 |
| iDeal ChIP-seq kit | Diagenode | C01010051 |
| KAPA HyperPrep kit for NGS DNA Library Prep | Roche | 7962363001 |
| MinElute PCR Purification Kit | QIAGEN | 28004 |
| MinElute Reaction Clean up kit | QIAGEN | 28206 |
| MycoAlert Assay Control Set | Lonza | LT07-518 |
| MycoAlert detection kit | Lonza | LT07-118 |
| NEBNext Multiplex Oligos for Illumina kit | New England Biolabs | E7500 |
| PureLink Genomic DNA Mini Kit | Thermo Fisher | K182002 |
| Quick Ligation™ Kit | New England Biolabs | M2200S |
| Rneasy Mini Kit | QIAGEN | 74104 |
| Vectashield | Vector laboratories | H-1000 |
| Zymo DNA Clean & Concentrator-5 kit | Zymo | D4013 |
| Deposited data |  |  |
| Raw and processed sequencing data | This study | GEO: GSEXXXXXX |
| HiC in mouse ESCs and Cortical Neurons | Bonev et al., 2017 | GEO: GSE96107 |
| HiC in mouse E11.5 limb buds | Kraft et al., 2019 | GEO: GSE116794 |
| ChIP-seq for CTCF, Rad21 and H3K9me3 in mouse ESCs and E11.5 limb buds | Kraft et al., 2019 | GEO: GSE116794 |
| ChIP-seq for H3K4me1, H3K4me3, H3K27ac and H3K27me3 in E11.5 limb buds | Andrey et al., 2017 | GEO: GSE84795 |
| ATAC-seq in mouse ESCs | Bauer et al., 2021 | GEO: GSE157448 |
| Fantom5 CAGE Expression datasets | Lizio et al., 2015 | https://fantom.gsc.riken.jp/5/data/ |
| DamID in mouse E11.5 limb cells | Allou et al., 2021 | GEO: GSE137335 |
| HiC in activated mouse B cells | Vian et al., 2018 | GEO: GSE98119 |
| HiC in mouse cardiomyocytes | Rosa-Garrido et al., 2017 | GEO: GSM2544836 |
| HiC in mouse olfactory cells | Monahan et al., 2019 | GEO: GSE112153 |
| HiC in mouse inner cell mass and 8-cell embryos | Du et al., 2017 | GEO: GSE82185 |
| HiC and H3K27ac & CTCF ChIP-seq in human ESCs and cardiomyocytes | Zhang, et al 2019 | GEO: GSE116862 |
| HiC in 48 hr hpf Zebrafish | Yang et al., 2020 | GEO: GSE134055 |
| HiC in axolotl AL-1 cells | Schloissnig et al., 2021 | SRA: PRJNA645452 |
| HiC in xenopus brain | Niu et al., 2021 | SRA: PRJNA606649 |
| HiC in pig embryonic fibroblasts | Li et al., 2020 | GEO: GSE153452 |
| DamID in human RPE and ESCs | van Schaik et al., 2020 | 4D nucleome |
| ChIP-seq for H3K36me3 in mouse ESCs | Encode | GEO: GSE31039 |
| scRNA-seq in gastrulating E6.5-8.5 mouse embryos | Pijuan-Sala et al., 2019 | ArrayExpress: E-MTAB-6967 |
| scRNA-seq in E9.5-E12.5 mouse embryos | Cao et al., 2019 | GEO: GSE119945 |
| scRNA-seq in E9.5-E14.5 mouse placentas | Marsh et al., 2020 | GEO: GSE152248 |
| Experimental models: Cell lines |  |  |
| G4 ESCs (XY, 129/Sv x C57BL/6 F1 hybrid | Georg et al. 2007 | N/A |
| CTCF-AID-GFP E14 ESCs | Elphege et al. 2017 | N/A |
| Rad21-AID-GFP E14 ESCs | Liu et al. 2020 | N/A |
| *mutant ESC lines are listed in Table 1 | This study | N/A |
| 293FT | Thermofisher | R70007 |
| Experimental models: Organisms/strains |  |  |
| Wildtype and mutant mice derived from G4 ESCs | This study | N/A |
| Opossums (*Monodelphis domestica*) | Naturkunde Museum, Berlin | N/A |
| Chicken (*Gallus Gallus*) | Valo Biomedica | N/A |
| Oligonucleotides |  |  |
| Rex1/Fat1 cHiC library | This study | mm10, chr8: 39022300-48000000 |
| DamID oligos and primers see Table S2 | Vogel et al, 2007 | N/A |
| WISH probe primers see table Table S2 | This study | N/A |
| Genotyping primers see table Table S2 | This study | N/A |
| Cloning primers see table Table S2 | This study | N/A |
| OligoPAINT probes see Table S3 | This study | N/A |
| Recombinant DNA |  |  |
| pLGW-Dam-V5-Lamin B1 (Mm) | Steensel Lab | N/A |
| pLGW-V5-Dam | Steensel Lab | N/A |
| pMD2.G | Bird Lab | N/A |
| psPAX2 | Bird Lab | N/A |
| BAC for *Fat1R* | CHORI/BACPAC | RP23-451E23 |
| pX459 pSpCas9(BB)-2A-Puro vector | Addgene | #62988 |
| Fat1 promoter 302bp | This study | chr8: 44935221 - 44935522 |
| Rex1 core promoter 100bp | This study | chr8: 43306912 - 43307011 |
| Rex1 midi promoter 602bp | This study | chr8: 43306912 - 43307513 |
| Rex1 full promoter 1219bp | This study | chr8: 43306912 - 43308130 |
| Triml promoter 427bp | This study | chr8: 43180161 - 43180587 |
| Fat1 enhancer | This study | chr8: 41591354 - 41594915 |
| Knockin donor vectors & corresponding pX459 sgRNAs see Table S1 | This study | N/A |
| Software and Algorithms |  |  |
| CRISPR design | https://www.benchling.com | N/A |
| R | https://www.r-project.org | N/A |
| MACS2.0 | https://github.com/taoliu/MACS | N/A |
| Bowtie2 | Langmead and Salzberg, 2012 | N/A |
| Samtools | http://samtools.sourceforge.net | N/A |
| HiCUP v0.5.9 | Wingett et al., 2015 | N/A |
| Juicer | Durand et al., 2016 | N/A |
| HiGlass | Kerpedjiev et al., 2018 | N/A |
| UCSC genome browser | https://genome.ucsc.edu | N/A |
| WashU browser | https://epigenomegateway.wustl.edu | N/A |
| Other |  |  |
| FISH and SBS-modelling statistics summary see Table S4 | This study | N/A |
| List of bridging species for conservation analysis see Table S6 | This study | N/A |

### RESOURCE AVAILABILITY

#### Lead contact

Further information and requests for resources and reagents should be directed to and will be fulfilled by the Lead Contacts Michael I. Robson and Stefan Mundlos.

#### Data and code availability

The sequencing data generated in this study will become available at the Gene Expression Omnibus.

### EXPERIMENTAL MODEL AND SUBJECT DETAILS

Mouse G4 ESCs (XY, 129S6/SvEvTac x C57BL/6Ncr F1 hybrid) were grown as described previously on a mitomycin-inactivated CD1 mouse embryonic fibroblast feeder monolayer on gelatinised dishes at 37°C, 7.5% CO2 (Andrey and Spielmann, 2017; George et al., 2007). CTCF-AID-GFP and *Rad21-AID-GFP* E14 ESCs were cultured feeder-free on gelatinised dishes at 37°C, 7.5% CO2. All ESCs were cultured in ESC medium containing knockout DMEM with 4,5 mg/ml glucose and sodium pyruvate supplemented with 15% FCS, 10 mM Glutamine, 1x penicillin/streptomycin, 1x non-essential amino acids, 1x nucleosides, 0.1 mM beta-Mercaptoethanol and 1000 U/ml LIF. Medium was changed every day while G4-cells were split every 2-3 days or were frozen at 1x 10^6^ cells/cryovial in ESC medium containing 20% FCS and 10% DMSO. ESCs and feeder cells were tested for Mycoplasma contamination using the MycoAlert detection kit and MycoAlert Assay Control Set.

E11.5 forelimb cells were isolated from C57BL/6 embryonic limbs through trypsinization, filtration (100 µm) and centrifugation. When cultured *in vitro*, forelimb cells were grown on gelatine-coated plates at 37°C in 7.5% CO2 for up to 48 h in limb medium (DMEM/F12, 10% FCS, 1% penicillin/streptomycin, 4 mM L-Glutamine) supplemented with 250 ng/ml Recombinant Mouse Wnt-3a protein and 150 ng/ml Recombinant Human/Mouse FGF-8b Isoform.

Mutant embryos and mutant live animals were produced through tetraploid or diploid aggregation, respectively (Artus and Hadjantonakis, 2011). Female mice of the CD1 strain were used as foster mothers. Mutant lines were established and maintained by crossing with wildtype C57Bl.6/J animals. All mice were housed in a centrally controlled environment with a 12 h light and 12 h dark cycle, temperature of 20–22.2 °C, and humidity of 30–50%. Bedding, food and water were routinely changed. All animal procedures were conducted as approved by the local authorities (LAGeSo Berlin) under the license numbers G0176/19, G0247/13 and G0243/18.

HH22 and HH24 Chicken embryos were extracted from fertilised chicken eggs (Valo Biomedia) incubated at 37.8°C, 45% humidity.

Embryonic stages of opossum originated from the breeding colony of *Monodelphis domestica* maintained under permit ZH104 (issued by the local authority, LAGeSo) in the animal care facility of the Museum für Naturkunde, Berlin. All opossums were housed in a centrally controlled environment with a reversed 12 h dark and 12 h light cycle, temperature of 24–26 °C, and humidity of 60-65%. Bedding, food and water were routinely changed. Females were euthanized using an overdose of Isoflurane under T0198/13 (issued by LAGeSo) according to national and international standards. Samples were taken immediately after death was confirmed.

### METHOD DETAILS

#### Plasmid Construction

SgRNAs were designed at desired structural variant breakpoints or knockin sites using the Benchling design tool (https://www.benchling.com/). Complementary sgRNA oligos were subsequently annealed, phosphorylated, and cloned into the BbsI site of dephosphorylated pX459 pSpCas9(BB)-2A-Puro vector (Addgene; #62988). For insertion of lacZ sensors, asymmetric homology arms surrounding insertion sites were first synthesised with a multiple cloning site that bisected, and so inactivated, the sgRNA. Once homology arms were cloned into a vector, the lacZ sensor insert harbouring the *β-globin* minimal promoter and polyA terminator were subsequently inserted by restriction digest (Symmons et al., 2014). For testing alternative promoters, the *β-globin* promoter was substituted for synthesized or PCR-amplified *Rex1, Triml1/2,* or *Fat1* promoters through restriction cloning. The bidirectional *Triml1/2* promoter was inserted to enable lacZ transcription from the *Triml2* orientation. For enhancer lacZ reporter experiments, the mouse *Fat1-* enh sequence was PCR-amplified and inserted into a phosphoglycerate kinase (PGK) promoter targeting vector containing FRT sites for insertion into C2 ESCs. A list of sgRNAs, corresponding homology constructs and resulting mutant ESCs can be found in Table S1. Cloned enhancer and promoter sequences can be found in the Key Resources Table. All plasmids are available on Addgene.

#### CRISPR-mediated genome editing

CRISPR was subsequently performed as described previously (Kraft et al., 2015). Briefly, 300,000 G4 ESCs (George et al., 2007) were seeded on CD1 feeders 16 h prior to transfection. For structural variants, ESCs were transfected with 4 μg of both sgRNAs targeting each breakpoint using FuGENE HD according to manufacturer’s instructions. For site-specific knockins, ESCs were transfected with 8 μg of the sgRNA and 4 μg of the homology construct. After 24 h, transfected cells were transferred onto puromycin-resistant DR4 feeders and treated with puromycin for 48 h. ESCs were grown for a further 4-6 days after which colonies were picked and transferred to CD1 feeders in 96-well plates. Plates were subsequently split into triplicates after 2-3 days, two for freezing and one for DNA harvesting. Following lysis and genotyping, selected clones were expanded from frozen plates after which genotypes were reconfirmed. Potential structural variant and knockin ESC clones were first identified by PCR-detection of unique deletion breakpoints or site-specific insertion breakpoints, respectively. Desired homozygous or heterozygous copy number were then determined by qPCR. All cell lines and corresponding genotyping primers can be found in Tables S1 and S2.

#### Enhancer Reporter Line Generation

The flippase (FLP)-flippase recognition target (FRT) system was used to introduce enhance-LacZ reporter constructs into C2 ESCs. This modified ESC line contains a phosphoglycerate kinase neomycin selection cassette flanked by FRT sites and a promoter- and ATG-less hygromycin cassette targeted downstream of the *Col1A1* locus (Beard et al., 2006). 800,000 C2 ESCs were seeded onto a feeder-coated 6-well plate and transfected with 9 μ g of targeting construct, 3 μg FLP-encoding vector, 1 μ l Lipofectamine LTX Plus reagent (Thermo Fisher Scientific), 20 μl Lipofectamine LTX in a to a final OPtiMEM volume 250 μl. After 24 h, transfected C2 cells were transferred onto hygromycin-resistant DR4 feeders and treated with hygromycin B (final concentration 150 μg/ml) in ES growth medium for 5-10 days. Colonies were then picked and transferred to CD1 feeders in 96-well plates. Plates were subsequently split into triplicates after 2-3 days, two for freezing and one for DNA harvesting. Following lysis and genotyping, selected clones were expanded from frozen plates after which genotypes were reconfirmed. Genetically modified C2 ESCs were used to produce embryos through diploid aggregation, and genotyping confirmed the presence of the desired mutations in the cells and later in the embryos. Enhancer reporter cell lines and corresponding genotyping primers can be found in Tables S1 and S2.

#### Auxin induced CTCF and Rad21 depletion

Available CTCF-AID-GFP and Rad21-AID-GFP ESCs E14 ESCs were treated with 500 µM auxin for 48 h and between 1-6 h, respectively (Liu et al., 2021; Nora et al., 2017). Successful depletion was confirmed through lost GFP signal by FACS. For CTCF-AID-GFP ESCs, bulk cell populations were plated on coverslips for FISH or directly fixed for cHiC. For cHiC on Rad21-AID-GFP ESCs, auxin-treated G1 cells were isolated by FACS following fixation and lysis for cHic and subsequent DAPI staining. For FISH on Rad21-AID-GFP ESCs, depleted cells were plated on coverslips following 2 h auxin-treatment where only modest changes to cell-cycle had occurred.

#### Western Blot

2 million mESCs were collected and then washed twice in PBS. The cell pellet was then resuspended in cell lysis buffer (2 5mM HEPES pH7.6, 5 mM MgCl2, 25 mM KCl, 0.05 mM EDTA, 10% Glycerol, 0.1% IGEPAL, 1X Roche protease inhibitor, 1mM DTT). Nuclei were pelleted from the cell lysate by centrifugation for 5 minutes at 1500 rpm. The nuclei were then washed once (10mM HEPES pH7.6, 3 mM MgCl2, 100 mM KCl, 0.01 mM EDTA, 10% glycerol, 1X Roche protease inhibitor, 1 mM DTT) and centrifuged at 3000g for 5 minutes. Nuclei were then resuspended in 150 µl RIPA Buffer and vortexed for 20 minutes at 4°C. This mixture was spun at 12,000 rpm for 15 minutes and the supernatant was collected for blotting. Western blots were performed with anti-Dnmt3a (1:2000) and anti-Dnmt3b (1:1000) and imaged using HRP chemiluminescence.

#### Tetraploid morula complementation

Mutant ESCs were seeded on CD1 feeders, grown for 2 days and then subjected to diploid or tetraploid aggregation, as previously described (Artus and Hadjantonakis, 2011). CD1 female mice were used as foster mothers. Genotypes of resulting embryos or animals was determined by genotyping PCR as performed in originating ESCs.

#### Whole mount *in situ* hybridisation

mRNAs were detected in embryos by WISH using digoxigenin-labelled antisense RNA probes transcribed from cloned mouse, opossum and chicken genomic sequences (PCR DIG Probe Synthesis Kit, Roche). Whole embryos were fixed overnight in 4% PFA/PBS, washed in PBS-Tween (PBST; 0.1% Tween) and then dehydrated for at least 10 min each in 25%, 50% and 75% methanol/PBST. Embryos were finally stored at −20°C in 100% methanol. For staining, embryos were rehydrated on ice in reversed methanol/PBST steps, washed in PBST, bleached in 6% H_2_O_2_/PBST for 1 h on ice. Following washing in PBST, embryos were then treated with 10 μg/ml proteinase K/PBST for 3 min, incubated in glycine/PBST, washed in PBST, and finally refixed for 20 min in 4% PFA/PBS, 0.2% glutaraldehyde, and 0.1% Tween 20. Following washing in PBST, embryos were incubated at 68°C in L1 buffer (50% deionized formamide, 5× saline sodium citrate, 1% SDS, 0.1% Tween 20 in diethyl pyrocarbonate, pH 4.5) for 10 min. Embryos were then incubated for 2 h at 68°C in hybridisation buffer 1 (L1 with 0.1% transfer RNA and 0.05% heparin). Afterwards, embryos were incubated overnight at 68 °C in hybridisation buffer 2 (hybridisation buffer 1 with 0.1% transfer RNA and 0.05% heparin and 1/500 digoxigenin probe). After overnight hybridisation, unbound probe was removed by 3 x 30 minute washing steps at 68°C in L1, L2 (50% deionized formamide, 2 × saline sodium citrate pH 4.5, 0.1% Tween 20 in diethyl pyrocarbonate, pH 4.5), and L3 (2 × saline sodium citrate pH 4.5, 0.1% Tween 20 in diethyl pyrocarbonate, pH 4.5). Subsequently, embryos were treated for 1 h with RNase solution (0.1 M NaCl, 0.01 M Tris pH 7.5, 0.2% Tween 20, 100 μg/ml RNase A in H_2_O), followed by washing in Tris-buffered saline, 0.1% Tween 20 (TBST 1) (140 mM NaCl, 2.7 mM KCl, 25 mM Tris-HCl, 1% Tween 20, pH 7.5). Embryos were then blocked for 2 h at room temperature in blocking solution (TBST 1 with 2% fetal bovine serum and 0.2% bovine serum albumin (BSA)), followed by incubation at 4 °C overnight in blocking solution containing 1:5,000 anti-digoxigenin-alkaline phosphatase. After overnight incubation, unbound antibody was removed by 6 × 30 min washings steps at room temperature with TBST 2 (TBST with 0.1% Tween 20 and 0.05% levamisole/tetramisole) and left overnight at 4 °C. At the next day, embryo staining was initiated by 3x 20 min washing steps in alkaline phosphatase buffer (0.02 M NaCl, 0.05 M MgCl_2_, 0.1% Tween 20, 0.1 M Tris-HCl and 0.05% levamisole/tetramisole in H_2_O) 3 × 20 min, followed by staining with BM Purple AP Substrate (Roche). At least three embryos were analysed from each mutant genotype. The stained embryos or their limb buds were imaged using a ZEISS SteREO Discovery.V12 with cold light source CL9000 microscope and Leica DFC420 digital camera. The sequences of primers used to generate *Triml2, Rex1, Fat1* are listed in Table S2.

#### LacZ staining in embryos

Whole-mount lacZ reporter staining was performed as previously described with minor adjustments (Lobe et al., 1999). E11.5 mouse embryos were dissected in cold PBS, fixed in 4% paraformaldehyde (PFA) in PBS on ice for 20 min and washed three times in lacZ buffer (2 mM MgCl2, 0.01% sodium deoxycholate, 0.02% Nonidet P-40 in PBS). Embryos were then incubated in staining solution (0.5 mg ml−1 X-gal, 5 mM potassium ferrocyanide, 5 mM potassium ferricyanide in lacZ buffer) at 37°C for a few hours to overnight until desired staining was achieved. Following staining, embryos were washed in lacZ buffer and imaged using a ZEISS SteREO Discovery.V12 with cold light source CL9000 microscope and Leica DFC420 digital camera. Embryos were stored at 4°C in 4% PFA in PBS.

#### RNA-seq

Isolated ESCs were trypsinized, heavily feeder depleted, centrifuged and snap frozen. E11.5 forelimb buds were microdissected from wildtype and mutant embryos in cold PBS and immediately snap-frozen for storage at −80°C. Total RNAs were extracted using the RNeasy Mini Kit according to the manufacturer’s instructions. Samples were poly-A enriched, prepared into libraries using the Kapa HyperPrep Kit, and sequenced on a Novaseq2 with 75 bp or 100 bp paired-end reads. RNA-seq experiments were performed at least in duplicates.

#### Sample collection for DamID-seq, ChIP-seq, ATAC-seq, cHiC and FISH

ESCs were trypsinized, heavily feeder depleted and pelleted by centrifugation. Chicken, opossum and mouse limb buds were microdissected from embryos in cold PBS. Isolated limbs were then trypsinised 5 minutes at 37°C with continuous agitation with a P1000 pipette until no visible clumps remained. Limb cell suspensions were then passed through a 40 µm filter, centrifuged at 250 g for 5 min. Supernatants were then removed from isolated ESCs or limb cells which could then be used for downstream applications.

#### DamID-seq

##### Lentiviral preparation and treatment

DamID was performed as described previously (Robson and Schirmer, 2016). Briefly, lentiviruses encoding the Dam methylase alone (pLgw V5-EcoDam) or fused to lamin B1 (pLgw-EcoDam-V5-Lamin) were generated in 293FT cells. Here, ∼6 million 293FT cells were transfected with 2.8 μg pMD2.G, 4.6 μg psPAX2, and 7.5 μg of pLgw V5-EcoDam or pLgw-EcoDam-V5-Lamin with 36 μl lipofectamine 2000 in 3 ml Optimem. After 16 h, 293FT media was replaced. Virus-containing supernatants were subsequently aspirated after 48 h and 72 h. Viral supernatants were then cleared of cellular debris by 10 min centrifugation at 3,500 rpm and subsequent filtration through a 0.45 μm^2^ low protein-binding PES syringe filter. Viral supernatants were finally purified using the Lent-X concentrator as per manufacturer’s instructions and resuspended in Optimem. If not used immediately, aliquots were frozen at -80°C.

To perform DamID, ESCs and cultured E11.5 limb cells were transduced with DamID lentiviruses and harvested 72 or 48 h later, respectively. Specifically, 1,5x10^5^ ESCs were plated feeder-free onto gelatinized 6 well 1 h prior to transduction with DamID lentiviruses. Transduction was then performed overnight after which virus-containing media was removed and cells were plated with feeders in 6 cm plates. After 48 h, contaminating feeders were removed by further feeder-depletion and pure ESCs were isolated by centrifugation. By contrast, isolated E11.5 limb bud cells were directly plated and transduced after 1 h. Virus-containing media was removed 24 h later after which cells were isolated after an additional 48 h.

##### DamID library processing

DamID sample processing was then performed as described previously (Robson and Schirmer, 2016). Briefly, DNA was extracted from cells using the DNeasy tissue lysis kit as per manufacturer’s instructions. 2.5 μg of extracted DNA was then digested by *DpnI* and, following heat inactivation of *DpnI*, was ligated to the DamID adaptor duplex (dsAdR) generated from the oligonucleotides AdRt (5’-CTAATACGACTCACATAGGGCAGCGTGGTCGCGGCCGA-GGA-3’) and AdRb (5’-TCCTCGGCCG-3’) after which DNA was further digested by *DpnII*. To amplify DNA sequences methylated by the Dam methylase, 5 μl of *DpnII* digested material was then subjected to PCR in the supplied buffer in the presence of the 1.25 μM Adr-PCR primer (5’-GGTCGCGGCCGAGGATC-3’), 0.2 mM dNTPs and 1X of the Advantage cDNA polymerase. PCR was performed as previously described after which amplified DNA was purified, processed into NGS libraries using the KAPA HyperPrep kit and analyzed for quality by Bioanalyzer analysis. standard protocols. DamID-seq samples were sequenced 75 or 100 bp paired-end reads and each experiment was performed in duplicates for sequencing.

#### ATAC-seq

ATAC-seq was performed as described previously (Buenrostro et al., 2015). Briefly, 1x10^5^ isolated E11.5 limb cells were employed per biological replicate. Cells were washed in cold PBS, lysed in fresh lysis buffer (10mM TrisCl pH7.4, 10mM NaCl, 3mM MgCl2, 0.1% (v/v) Igepal CA-630) for 2 min on ice, and finally pelleted for 10 min at 500 x g and 4°C. Following supernatant aspiration, nuclei-containing pellets were subjected to transposition using Tn5 Transposase for 30 min at 37° C. Resulting DNA was then purified using MinElute Reaction Clean up kit, eluted in 11 µl of elution buffer and stored in -20° C, if not immediately processed further. Barcoded adapters were added to the transposed fragments by PCR. To avoid saturation in our PCR, we initially performed 5 cycles and extracted a 5 µl aliquot for qPCR to identify the number of cycles required without overamplification. Nextera qPCR primers were used for the amplification. The remaining 45 µl of the PCR reaction were then amplified for the desired number of cycles which never exceeded 12. Finally, samples were purified on AMPure XP beads and eluted in 20 µl. Concentration was measured with Qubit and the quality of the samples was estimated by Bioanalyzer analysis. ATAC-seq samples were sequenced yielding for 50 million 75 bp paired-end reads and each experiment was performed in duplicate.

#### ChIP-seq

ChIP-seq was performed using the iDeal ChIP-seq kit for histones with several modifications. Briefly, ESCs were fixed in 1% paraformaldehyde (PFA)/10% FCS/PBS for 10 min with rotation at room temperature. Fixation was stopped by glycine after which cells were pelleted by centrifugation (8 min, 250 x g, 4°C). Cells were lysed in Lysis buffer (50 mM Tris, pH 7.5; 150 mM NaCl; 5 mM EDTA; 0.5% NP-40; 1.15% Triton X-100; protease inhibitors) for 10 min on ice. Nuclei were resuspended in sonication buffer (10 mM Tris–HCl, pH 8.0; 100 mM NaCl; 1 mM EDTA; 0.5 mM EGTA; 0,1% Na-deoxycholate; 0.5% N-lauroylsarcosine; protease inhibitors). Chromatin was sheared using a Bioruptor until reaching a fragment size of 200–500 base pairs. Afterwards, samples were processed with the iDeal ChIP-seq kit according to the manufacturer’s instructions. For each Histone ChIP 5 μg chromatin was used in combination with antibodies against H3K4me1 (1 µg) H3K4me3 (1 µg), H3K27ac (1 µg) and H3K27me3 (1 µg). Libraries were prepared for sequencing using the KAPA HyperPrep kit and their quality confirmed by Bioanalyzer analysis. ChIP-seq libraries were finally sequenced at 100 bp paired-end reads with all samples analyzed in biological duplicates.

#### ChIPmentation

For chicken embryonic limb buds, ChIPmentation libraries were prepared as previously described (Schmidl et al., 2015). Briefly, dissociated limb cells were filtered through a 70um MACS® SmartStrainer before fixation with 1% MeOH-free formaldehyde in PBS on ice for 10 minutes. Fixation was quenched using glycine, and the pellet was collected after centrifugation (3000rpm, 5 min, 4°C. Cells were then lysed in lysis buffer (10mM Tris pH 8.0, 100mM NaCl, 1mM EDTA pH 8.0, 0.5mM EGTA, 0.1% Sodium deoxycholate, 0.5% N-lauroylsarcosine) on ice, before shearing with a Covaris E220 for a fragment distribution of 200-700bp. Sheared chromatin was incubated with appropriate histone antibodies overnight at 4C. Antibodies were bound for immunoprecipitation with Dynabeads™ Protein G. Tn5-mediated ”tagmentation” of pull-downed chromatin was incubated at 37°C for 5min. Chromatin was de-crosslinked with Proteinase K at 65°C overnight. DNA was then purified using the MinElute Reaction Cleanup kit.

Nextera indexing primers (single-indexed) were added during library amplification. The number of PCR cycles for each library was estimated using Ct values as determined by qPCR (where number of cycles = rounded up Ct value +1). After amplification, DNA was cleaned up with AmPure XP beads, and then checked on a TapeStation D5000 HS for size distribution. Size selection was then carried out accordingly, with either a left-sided selection or a double-sided selection. The concentration of final eluted DNA was measured using Qubit HS and checked again on a TapeStation D5000HS. All libraries were sequenced on a Novaseq2 using 100bp paired-end reads. The same histone antibodies used for traditional ChIP-seq were also used here for ChIPmentation.

#### WGBS

Genomic DNA was extracted from ESCs and E11.5 limb buds using the PureLink Genomic DNA Mini Kit following manufacturer’s instructions. gDNA was then sheared in Covaris micro TUBE AFA Fiber Pre-Slit Snap-Cap tubes. Next, the sheared gDNA was purified with the Zymo DNA Clean & Concentrator according to manufacturer’s instructions. Purified DNA was then bisulfite converted using the EZ DNA Methylation-Gold Kit, and WGBS libraries were processed using the Accel-NGS Methyl-seq DNA library kit following manufacturer’s recommendations for each. Libraries were prepared and cleaned using Agencourt AMPure XP beads. The absence of adapters from the final libraries was verified using the Agilent TapeStation. WGBS libraries were sequenced on the NovaSeq6000 yielding 150 base pair paired-end reads.

#### Capture HiC

##### SureSelect design

The cHiC SureSelect library was designed over the genomic interval (mm10, chr8:x-y) using the SureDesign tool from Agilent.

##### Fixation

Disassociated ESCs and limb cells were transferred to a 50-ml falcon tube and complemented with 10% FCS/PBS. 37% formaldehyde was added to a final concentration of 2% and cells were fixed for 10 min at room temperature. Crosslinking was quenched by adding glycine (final concentration; 125 mM). Fixed cells were washed twice with cold PBS and lysed using fresh lysis buffer (10 mM Tris, pH 7.5, 10 mM NaCl, 5 mM MgCl2, 0.1 mM EGTA with protease inhibitor) to isolate nuclei. Cell lysis was assessed microscopically after 10-min incubation in ice. Nuclei were centrifuged for 5 min at 480g, washed once with PBS and snap frozen in liquid N2.

##### cHiC library preparation and sequencing

3C libraries were prepared from fixed nuclei as described previously (Kragesteen et al, 2018). Briefly, lysis buffer was removed by centrifugation at 400 g for 5 min at 4 °C, followed by supernatant aspiration, snap-freezing, and pellet storage at − 80 °C. Later, nuclei pellets were thawed on ice, resuspended in 520 μl 1× DpnII buffer, and then incubated with 7.4 μl 20% SDS shaking at 900 rpm. at 37 °C for 1 h. Next, 75 μl 20% Triton X-100 was added and the pellet was left shaking at 900 rpm at 37°C for 1 h. A 15-μl aliquot was taken as a control for undigested chromatin (stored at − 20°C). The chromatin was digested using 40 μl 10 U/μl DpnII buffer shaking at 900 rpm at 37°C for 6 h; 40 μl of DpnII was added and samples were incubated overnight, shaking at 900 rpm. at 37°C. On day three, 20 μl DpnII buffer was added to the samples followed by shaking for an additional 5 h at 900 rpm. at 37 °C. DpnII subsequently was inactivated at 65 °C for 25 min and a 50-μl aliquot was taken to test digestion efficiency (stored at − 20 °C). Next, digested chromatin was diluted in 5.1 ml H2O, 700 μl 10× ligation buffer, 5 μl 30 U/μl T4 DNA ligase and incubated at 16°C for 4 h while rotating. Ligated samples were incubated for a further 30 min at room temperature. Chimeric chromatin products and test aliquots were de-cross-linked overnight by adding 30 μl and 5 μl proteinase K, respectively, and incubated at 65 °C overnight. On the fourth day, 30 μl or 5 μl of 10 mg ml−1 RNase was added to the samples and aliquots, respectively, and incubated for 45 min at 37°C. Next, chromatin was precipitated by adding 1 volume phenol-chloroform to the samples and aliquots, vigorously shaking them, followed by centrifugation at 4,000 rpm at room temperature for 15 min. To precipitate aliquot chromatin, 1 volume 100% ethanol and 0.1 volume 3M NaAc, pH 5.6 was added and the aliquots placed at -80°C for 30 min. DNA was then precipitated by centrifugation at 5,000 rpm. for 45 min at 4°C followed by washing with 70% ethanol, and resuspension in 20 μl with 10 mM Tris-HCl, pH 7.5. To precipitate samples, extracted sample aqueous phases were mixed with 7 ml H2O, 1 ml 3M NaAc, pH 5.6, and 35 ml 100% ethanol. Following incubation at −20°C for at least 3 h, precipitated chromatin was isolated by centrifugation at 5,000 rpm for 45 min at 4 °C. The chromatin pellet was washed with 70% ethanol and further centrifuged at 5,000 rpm for 15 min at 4 °C. Finally, 3C library chromatin pellets were dried at room temperature and resuspended in 10 mM Tris-HCl, pH 7.5. To check the 3C library, 600 ng were loaded on a 1% gel together with the undigested and digested aliquots. The 3C library was then sheared using a Covaris sonicator (duty cycle: 10%; intensity: 5; cycles per burst: 200; time: 6 cycles of 60 s each; set mode: frequency sweeping; temperature: 4– 7 °C). Adaptors were added to the sheared DNA and amplified according to the manufacturer’s instructions for Illumina sequencing (Agilent). The library was hybridised to the custom designed SureSelect beads and indexed for sequencing (75–100 bp paired-end) following the manufacturer’s instructions (Agilent).

#### HiC

HiC libraries were prepared as described in a previously published in situ protocol (Melo et al., 2020; Rao et al., 2014). Briefly, ∼1 million cells were fixed in 2% formaldehyde, lysed, and digested overnight with DpnII enzyme. Digested DNA ends were marked with biotin-14-dATP and ligated overnight using T4 DNA ligase. Formaldehyde crosslinking was reversed by incubation in 5 M NaCl for 2 h at 68°C, followed by ethanol precipitation. A S-Series 220 Covaris was used to shear the DNA to fragments of 300–600 bp for library preparation, and biotin-filled DNA fragments were pulled down using Dynabeads MyOne Streptavidin T1 beads. DNA ends were subsequently repaired using T4 DNA polymerase and the Klenow fragment of DNA polymerase I and phosphorylated with T4 Polynucleotide Kinase NK. DNA was further prepared for sequencing by ligating adaptors to DNA fragments, using the NEBNext Multiplex Oligos for Illumina kit. Indexes were added via PCR amplification (4–8 cycles) using the NEBNext Ultra II Q5 Master Mix. PCR purification and size selection were carried out using Agencourt AMPure XP beads. Libraries were sequenced on a x platform yielding x bp paired-end reads. For each sample, the HiC library was created by pooling a total of four technical replicates generated from two different cell isolations cultures in order to ensure higher complexity of the sequencing library.

#### Oligopaint fluorescence *in situ* hybridisation with 3D-SIM imaging

##### Oligopaint library assembly

Oligopaint libraries were constructed as described by previously (Beliveau et al., 2015); see the Oligopaints website (https://oligopaints.hms.harvard.edu) for further details. Libraries were ordered from CustomArray in the 92K Oligo pool format. The mm10 coordinates, size, number, density of oligonucleotides and primers used for the libraries are listed in Tables S3. Oligopaint oligos were identified using the archived mm10 ‘balance’ BED files, which consist of 35–41-mer genomic sequences throughout the regions of interest (Beliveau et al., 2018). BED files can be retrieved from the Oligopaints website. Each library contains a universal primer pair followed by a specific primer pair hooked to genomic sequences (119-125 mer oligonucleotides). Oligopaint libraries were produced by emulsion PCR amplification from oligonucleotide pools followed by a ‘two-step PCR’ procedure and the lambda exonuclease method described by Beliveau et al. (Beliveau et al., 2015). The two-step PCR leads to the addition of a specific binding sequence for signal amplification with a secondary oligonucleotide (Sec1-Alexa 488 for green probes or Sec6-Atto 565 for red probes) containing two additional fluorophores. Consequently, each probe carries three fluorophores in total. This strategy allows for the 2-color imaging between different combinations of the oligopaint probes. All oligonucleotides used for Oligopaint production were purchased from Integrated DNA Technologies. Oligonucleotide primer sequences (5′→3′) used for this approach are listed in Table S3.

##### BAC probe preparation

The BAC probe corresponding to the Fat1 gene was labeled with the Alexa Fluor 555 using the FISH Tag DNA Kit.

##### FISH and immunostaining

FISH was performed as described previously (Szabo et al., 2020). Briefly, 1,5-2 x10^5^ isolated ESCs or E11.5 limb cells were plated from single-cell suspensions onto 0.01% poly-lysine coated coverslips (170 ± 5 µm) for 2 h. Cells were fixed for 10 min in PBS/4% PFA, washed three times in PBS, incubated for 10 min in PBS/0.5% Triton X-100, washed three times in PBS, incubated for 10 min in 0.1 M of HCl and washed twice in 2× SSC/0.1% Tween 20 (2× SSCT). Cells were then incubated in 50% formamide/2× SSCT (20 min at room temperature followed by 20 min at 60 °C). Hybridisation solution was made with 20 µl of FISH hybridisation buffer (50% formamide, 10% dextran sulfate, 2× SSC and salmon sperm DNA (final concentration 0.5 mg/ml)), 0.8 µl of RNase A (10 mg/ml) and Oligopaint probes (primary and secondary probes at 1–3 µM final concentration). When required, co-hybridization of Oligopaints with the *Fat1* BAC probe was performed using 25 ng of BAC probe together with a 50x excess of mouse Cot-1 DNA. Hybridisation solution was deposited on coverslips that were then sealed on glass slides with rubber cement. Slides were placed on a heating block immersed in a water bath for 3 min at 80 °C for denaturation. Probe hybridisation was performed overnight at 42 °C in a dark and humid chamber. Coverslips were removed from glass slides and washed for 15 min in 2× SSCT at 60 °C, 10 min in 2× SSCT at room temperature, 10 min in 0.2× SSC and in PBS. Cells were then washed in PBS/0.1% Tween 20 (PBT) and incubated for 1 h in PBT/2%BSA. Primary antibody (ant-lamin B1, 1:1,000 dilution in PBT/2% BSA) incubation was performed overnight at 4 °C between coverslips and glass slides in a humid and dark chamber. Cells were washed four times in PBT and secondary antibody (anti-rabbit-IgG-Atto 647, 1:100 dilution in PBT/2% BSA) incubation was performed for 1 h at room temperature between coverslips and glass slides in a dark and humid chamber. Last, cells were washed in PBT, stained with DAPI (final concentration at 1 µg/ml in PBS) and washed at least 3 times for 5 min each in PBS. Coverslips were mounted on slides with VECTASHIELD and sealed with nail polish.

##### Image acquisition

3D-SIM imaging was carried out with a DeltaVision OMX V4 microscope equipped with an ×100/1.4 numerical aperture (NA) Plan Super Apochromat oil immersion objective (Olympus) and electron-multiplying charge-coupled device (Evolve 512B; Photometrics) camera for a pixel size of 80 nm. Diode lasers at 405, 488, 561 and 647 nm were used with the standard corresponding emission filters. *Z*-stacks (*z*-step of 125 nm) were acquired using 5 phases and 3 angles per image plane. Raw images were reconstructed using SoftWorx v.6.5 (GE Healthcare Systems) using channel-specific optical transfer functions (pixel size of reconstructed images = 40 nm). TetraSpeck beads (200 nm) (T7280, Thermo Fisher Scientific) were used to calibrate alignment parameters between the different channels. The quality of reconstructed images was assessed using the SIMcheck plugin of ImageJ v.1.52i (Ball et al., 2015).

### QUANTIFICATION AND STATISTICAL ANALYSIS

#### RNA-seq differential expression analysis

Single-end, 100 bp reads from Illumina sequencing were mapped to the reference genome (mm10) using the STAR mapper (splice junctions based on RefSeq; options: --alignIntronMin20 -- alignIntronMax500000 --outFilterMismatchNmax 10). Differential gene expression was ascertained using the DESeq2 package (Love et al., 2014). The cut-off for significantly altered gene expression was an adjusted P value of 0.05.

#### Single cell RNA-seq

The expression of Triml2, Rex1, and Fat1 genes was investigated in three sc-RNAseq datasets of early mammal development, whole placenta (Marsh and Blelloch, 2020), whole embryo gastrulation (Pijuan-Sala et al., 2019), and whole embryo organogenesis (Cao et al., 2019). For visualization, we used the originally reported Uniform Manifold Approximation and Projection (UMAP) embeddings for the whole placenta and the gastrulation datasets and the t-Distributed Stochastic Neighbor Embedding (tSNE) for the organogenesis dataset. Likewise, we used the reported cell type definitions for visualization. For the whole placenta dataset, we used the “integrated_snn_res.0.6” cell variable to color cell types. UMI counts for *Triml2*, *Rex1*, and *Fat1* were plotted for all datasets in the range 0 to >2.

#### DamID-seq analysis

Raw reads from DamID-seq experiments were mapped to the mouse mm10 reference genome using the alignment tool BWA-MEM (v.0.7.12) (Li and Durbin, 2009). The counts of mapped reads overlapping a DpnII (GATC) restriction fragment side were normalized by reads per kilobase, divided by the length of the fragment, per million mapped reads (RPKM). Based on these normalized counts the log2 fold change between the Dam–Lamin B1 transduced samples and the respective Dam-only-encoding samples was calculated.

#### ATAC-seq analysis

Raw sequencing fastq files were processed using cutadapt (Martin, 2011) for adapter trimming, Bowtie2 (Langmead and Salzberg, 2012) for mapping, SAMtools (Li et al., 2009) for filtering, sorting and removing duplicates, and deepTools (Ramirez et al., 2016) for generating coverage tracks.

#### ChIP-seq analysis

Raw sequencing fastq files were processed using STAR (Dobin et al., 2013) for mapping, SAMtools (Li et al., 2009) for filtering, sorting and removing duplicates, and deepTools (Ramirez et al., 2016) for generating coverage tracks.

#### Enhancer prediction

Enhancers were predicted using a series of established tools for ATAC-seq peak prediction and enhancer / promoter prediction. First, Genrich (not published, https://github.com/jsh58/Genrich/) was used to predict ATAC-seq peaks. We filtered for those that overlap a enhancer predicted by CRUP (Ramisch et al., 2019) and do not overlap an annotated TSS (UCSC) or a promoter predicted by eHMM (Zehnder et al., 2019).

#### Enhancer conservation analysis

ATAC-seq peaks and predicted enhancers were projected between mouse, opossum and chicken using a stepped pairwise sequence alignment approach in multiple bridging species (Baranasic et al., 2021). The basic concept of the approach is indicated schematically below.

##### Genomic coordinate projection schematic illustration

**Left**. An example genomic location X is projected between observed (e.g. mouse) and target species (e.g. opossum) using the direct alignments (grey rectangles) and the alignments via a bridging species (e.g. human, blue and red rectangles). Projections are indicated as a black X in the respective species). Dashed lines connect pairwise sequence alignments. The projected locations of X in observed species are indicated in grey (direct alignments) and black (via bridging species). **Right.** Example graph comprising 13 species (nodes). For any genomic location, the shortest path through the species graph yields the combination of species which maximizes projection accuracy.

For a genomic region with conserved synteny, any non-alignable coordinate can be approximately projected from one genome to another by interpolating its relative position between two alignable anchor points. The accuracy of such interpolations correlates with the distance to an anchor point. Therefore, projections between species with large evolutionary distances tend to be inaccurate due to a low anchor point density. Including so-called bridging species increases the anchor point density and thus improve projection accuracy. The optimal choice and combination of bridging species may vary from one genomic location to another. This presents a shortest path problem in a graph where every node is a species and the weighted edges between nodes correspond to a scoring function that represents the distances of genomic locations to their anchor points (|x - a|). The scoring function exponentially decreases with increasing distances |x - a|. The shortest path problem is solved using Dijkstra’s Shortest Path Algorithm (Dijkstra, 1959). The used sets of bridging species are given in Table S6.

Projected elements from ATAC-seq peaks were then classified into directly (DC), indirectly (IC) or not conserved (NC) according to the following criteria: DC elements overlap a direct sequence alignment between the reference and the target species. IC elements do not overlap a direct alignment, but are projected with a score > 0.99, i.e. either overlapping or in direct vicinity to a multi-species anchor. A score of > 0.99 means that the sum of the distances from the element and its intermediate projections to their respective anchor points is < 150 bp throughout the optimal bridging species path. The remaining peaks are classified as non-conserved (NC).

#### cHiC and HiC analysis

##### cHiC analysis

Raw fastq files had read lengths of 75bp and 100bp, respectively. In a preprocessing step, fastq files with 100 bp read length were trimmed to 75bp to achieve comparable initial read lengths for all samples. Afterwards, fastq files were processed with the HiCUP pipeline v0.8.1 (no size selection, Nofill: 1, Format: Sanger) for mapping, filtering and deduplication steps (Wingett et al., 2015). The pipeline was set up with Bowtie 2.4.2 for mapping short reads to reference genome mm10 (Langmead and Salzberg, 2012). If replicates were available, they were merged after the processing with the HiCUP pipeline. Binned and KR normalized cHiC maps (Knight and Ruiz, 2012; Rao et al., 2014) were generated using Juicer tools v1.19.02 (Durand et al., 2016). Only read pairs for region chr8:39,030,001-48,000,000 and with MAPQ≥30 were considered for the generation of cHiC maps.

Additional to the original cHiC maps, custom reference genomes were derived from mm10 for the three deletions (ΔD1, ΔD2, ΔD1+2), considering the respective deletions and cHiC data was processed correspondingly. cHiC and HiC maps were displayed as heatmaps in which very high values were truncated to improve the visualization.

##### HiC analysis

Fastq files were processed with the Juicer pipeline v1.5.6 (Durand et al., 2016) (CPU version) using bwa v0.7.17 (Li and Durbin, 2010) for mapping short reads to the reference genomes mm10 (mouse), hg19 (human), galGal6 (chicken), monDom5 (opossum), susScr11.1 (pig), and AmexG_v6.0-DD (axolotl), respectively. Replicates were merged after the mapping, filtering and deduplication steps of the Juicer pipeline. Juicer tools v1.7.5 (Durand et al., 2016) were used to generate binned and KR normalized HiC maps from read pairs with MAPQ≥30.

For compartment analysis, hic-files were converted at 100kb bin size to the cool format using hic2cool (v0.8.2) (https://github.com/4dn-dcic/hic2cool) and balanced using cooler (v0.8.5) (Abdennur and Mirny, 2020). Afterwards, compartment analysis was performed using cooltools (v0.3.0) (https://github.com/open2c/cooltools) and using the GC content as reference track.

TADs were identified by applying TopDom v.0.0.228 on 50-kb binned and KR-normalized maps using a window size of 10 (Shin et al., 2016).

#### Gene co-regulation in TADs analysis

To calculate gene-expression correlations, we downloaded FANTOM stage 5’ CAGE TPM data (https://fantom.gsc.riken.jp/5/data/). We discarded samples annotated as belonging to ‘reference’ ‘whole body’ or similar samples, and also excluded testis and related tissues from the analysis.

We also removed all libraries with fewer than 1 million reads, and all peaks with less than 32 reads across all samples. Overlapping each peak with the Gencode M23 annotation, we assigned peaks to genes if they overlapped a Gencode exon for that gene, or were less than 200bp upstream of a TSS. Peaks not overlapping a gene were discarded, and the counts for all of a genes’ peaks were summed.

Since the FANTOM data contained the resulting gene x sample count matrix was then normalized as per as per (Alam et al., 2020) – normalized counts-per-million for each sample. As many of the sample in the FANTOM CAGE data were highly correlated (due e.g. to being replicates or adjacent time points), we performed hierarchical clustering on the 829 remaining datasets, and then merged libraries with a pearson correlation of 0.95 or greater, resulting in a final 349 metasamples. Co-expression between two genes was then defined as pearson correlation across these 349 metasamples.

To identify housekeeping genes (Figure S1B), we replicated the procedure used by FANTOM previously (Consortium et al., 2014). Here, the 2D density of median and maximum normalized expression over all samples is first plotted, and then setting a cutoff on median expression that separated ubiquitous from non-ubiquitous genes. To assess the relationship between co-expression and linear gene distance separation or TAD co-occupancy and co-expression we next identified TADs in ESCs, E11.5 limb buds and cortical neurons (Bonev et al., 2017; Kraft et al., 2019). Plotting co-expression as a function of distance revealed, as expected, a strong relationship between linear proximity in the genome and co-expression. Since genes sharing TADs are necessarily more likely to be closely spaced, we plotted (log10) linear distance against co-expression separately for pairs either sharing or not sharing a TAD, pooling gene pairs with similar linear distance in a moving average over 2000 points (Fig S3D). Mean Corr. Values were calculated by averaging correlations for all gene pairs within a TAD (Figure 1D).

#### WGBS processing

Raw reads were subjected to adapter and quality trimming using cutadapt (version 2.4; parameters: --quality-cutoff 20 --overlap 5 --minimum-length 25; Illumina TruSeq adapter clipped from both reads), followed by trimming of 10 nucleotides from the 5’ end of the first read, 15 nucleotides from the 5’ end of the second read and 5 nucleotides from the 3’ end of both reads (Kechin et al., 2017). The trimmed reads were aligned to the mouse genome (mm10) using BSMAP (version 2.90; parameters: -v 0.1 -s 16 -q 20 -w 100 -S 1 -u -R) (Xi and Li, 2009). Duplicates were removed using the ‘MarkDuplicates’ command from GATK (version 4.1.4.1; --VALIDATION_STRINGENCY=LENIENT --REMOVE_DUPLICATES=true) (McKenna et al., 2010). Methylation rates were called using mcall from the MOABS package (version 1.3.2; default parameters) (Sun et al., 2014). All analyses were restricted to autosomes and only CpGs covered by at least 10 reads and at most 150 reads were considered for downstream analyses.

#### Differentially methylated region (DMR) calling

DMRs were called using metilene (version 0.2-8; parameters: -m 10 -d 0.2 -c 1 -f 1) (Juhling et al., 2016) using two replicates per condition and filtered for a *Q*-value < 0.05.

#### SBS-polymer modelling with NE-attachment

We simulated the 3D structure of the *Fat1/Rex1* locus in ESC and E11.5 limb buds using a Strings and Binders Switch (SBS) polymer model that incorporates NE-attachment as described below (Barbieri et al., 2012; Chiariello et al., 2016; Nicodemi and Prisco, 2009).

##### Polymer model

Briefly, the SBS polymer model simulates a chromatin filament as a string with *N* beads, possessing potential binding sites for specific interacting molecules (binders). The binder concentration*c* and bead-binder interaction energies *E*_*int*_ control the system’s state through a coil-globule transition occurring when they are above a threshold (Barbieri et al., 2012; Chiariello et al., 2016). The type and location of binding sites specific for different regions of the *Rex1/Fat1* locus were inferred from ESC or E11.5 limb cHi-C data using PRISMR (mm10 chr8: 40300000 - 46200000; 20 Kb resolution) (Bianco et al., 2018). This machine-learning based algorithm returns the minimal arrangement of binding sites to fit the input. As output, the best polymer modelling the *Fat1/Rex1* locus was generated with 13 distinct types of binding sites in each condition. From these polymers, we obtain a set of 3D structures representing chromatin conformations in ESC and E11.5 limb through standard Molecular Dynamics simulations (see below).

##### Details of Molecular Dynamics simulations

In order to build an ensemble of 3D structures representing the Rex1 locus in E11.5 limb and ESC cell lines, we perform extensive Molecular Dynamics (MD) simulations (Chiariello et al., 2016). For simplicity, bead and binders have the same diameter *σ* = 1 and mass *m* = 1 in dimensionless units. A standard truncated Lennard-Jones (LJ) potential models the hard-core repulsion between the objects. By contrast, interaction between beads and binders is modelled with an attractive LJ potential with distance cutoff ranging from *R*_*int*_ = 1.3*σ* to *R*_*int*_= 1.5*σ* and an interaction intensity, given by the minimum of the LJ potential, within the range of *E*_*int*_ = 3.1 − 8.2*K_B_T*. An additional non-specific, weaker interaction (in the *E*_*int*_ = 2 − 3*K_B_T* range) is set among binders and the polymer. Consecutive beads of the polymer are linked by FENE bonds (Kremer and Grest, 1990) with standard parameters (length *R*_0_ = 1.6*σ* and spring constant *K*_*FENE*_ = 30*K_B_T*/*σ*^2^). Beads and binders move through Brownian dynamics according to the standard Langevin equation (Allen and Tildesley, 1989) with temperature *T* = 1, a friction coefficient *ζ* = 0.5 and an integration time step *Δt* = 0.012 (dimensionless units). The polymer is first initialized as a Self-Avoiding-Walk and the binders are randomly located in the simulation box, then the system is equilibrated up to approximately 10^8^ timesteps. From each model, we perform up to 10^2^ independent simulations in which polymer configurations are sampled every 5*10^5^ timestep once equilibrium is reached. Simulations are performed with the LAMMPS package (Plimpton, 1995).

##### Modelling the nuclear envelope

To model the NE, we introduce a spherical wall of radius *R* within the simulation box. Polymer beads can attractively interact with NE though a short range, truncated LJ potential with affinity *E*_*int*_ ranging from 0.0*K_B_T* to 10*K_B_T* and cutoff distance *r*_*cutoff*_ = 2.5*σ*. Among the NE-bead interaction energies tested, the structures obtained immediately after the NE-polymer adsorption (around 1.2*K_B_T*) generated structural measurements that most closely matched those observed by FISH (Figure S5). Alternatively, beads interact with NE only through a purely repulsive LJ potential. The NE sphere radius is set to *R* = 40*σ*. In order to define the interaction state (repulsive or attractive) of each polymer bead with NE, we employ DamID data for each wild or mutant ESC/limb sample. Briefly, we compute the average DamID signal in each 20kb window and evaluate its sign. Polymer beads associated with an average positive DamID signal are classified as attractively interacting with NE. Conversely, beads associated with a negative signal experience only a repulsive interaction. In this way, regions enriched with DamID tend to attach to the NE in the model. In our simulations, the NE is introduced after the SBS (polymer+binders) system is equilibrated, as described in the previous section. Then, in order to ensure the complete interaction of the polymer with the NE, the system is equilibrated up to other 7*10^7^ timesteps.

##### Quantification of measurements

Pairwise distance distributions are extracted from the population of 3D polymer structures as previously described (Chiariello et al., 2020; Conte et al., 2020). For each pair of objects, we first compute the center of mass of the polymer beads belonging to that object, then we evaluate the distance between the centers of mass. This distance is then averaged over the last 20 frames of each simulation. In order to map dimensionless length scale into physical units we compare pairwise distances measured by FISH. In total, we compare six different probe pairs (D1-D2, Fl1-Fl2, Rex1R-D1, Rex1R-D2, Rex1R-Fl1, Rex1R-Fl2) both in E11.5 limb and ESCs, for each pair we equalize the model and experimental median and then average over the different probe pairs. The resulting length scale mapping factor is *σ* = 44nm.

Distances from NE shown in Figure S5E and S6A are estimated by computing: 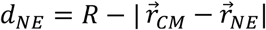, where *R* is the model NE radius, 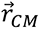 is the position of the center of mass of the object and 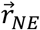 is the position of the NE center. Physical distances are then obtained using the mapping factor *σ* previously calculated from the comparison with pairwise FISH distances.

Pairwise overlaps between two objects shown in Figures S6D and S6F are obtained by using the following expression: *overlap*_12_ = *A*_12_/(*A*_1_+*A*_2_ – *A*_12_) where *A*_1_ and *A*_2_ are the surfaces of 2D projections associated to object 1 and object 2 respectively and *A*_12_ is their common area. For simplicity, 2D projections are approximated as circles whose radii *R*_1_ and *R*_2_ are estimated as gyration radii from the projected coordinates, so 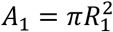 and 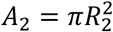. In this way, overlapping areas can be easily estimated using standard geometric relations. Indeed, given the distance *d* between the centers of the projected objects and supposing, without loss generality, *R*_2_ > *R*_1_, we have a partial overlap if *R*_2_ − *R*_1_ < *d* < *R*_1_ + *R*_2_. In this case: 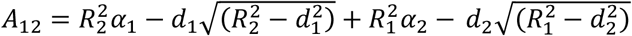, where 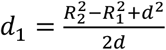 and 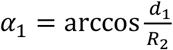 (analogous relations hold for *d*_2_ and *α*_2_). If *d* < *R*_1_ + *R*_2_, we impose *A*_12_ = 0, i.e. objects are well separated in space; finally, if *d* ≤ *R*_2_ − *R*_1_, we set 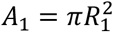, i.e. object 1 is completely contained within object 2. Three body overlaps shown in Figures S5E and S6B involving Rex1R or Fat1 with D1+D2, are defined as: *overlap*_123_ = *A*_12_/(*A*_1_ + *A*_2_ + *A*_3_ − *A*_12_ − *A*_13_ − *A*_23_), where object 1 can be Rex1R or Fat1. As for 3D distances, overlap values are averaged over the last 20 frames of each simulation. Analogously, a geometric mapping factor of 1.2 is found when comparing with pairwise experimental medians.

Sphericity is defined using the standard formula: 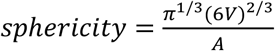, where *A* and *V* are area and volume of the object respectively. Area and volume are estimated from the coordinates of the polymer beads belonging to the region under consideration by means of a 3D convex hull approximation, computed with the Python package scipy.spatial. Sphericity measurements can be viewed in Figures S5E and S6C.

Contact maps are computed as previously described (Chiariello et al., 2016; Conte et al., 2020). We first measure the distance *r*_*ij*_ between any two beads *i* and *j*. If the distance is lower than threshold (7.5*σ* in Figure S5B, C), the beads are in contact. For each considered condition (without NE and with NE at different interaction energies), aggregated matrices are obtained over the different independent simulations. Visual and quantitative comparisons reveal a general good agreement between model and cHi-C data in both cell lines (Pearson *r* = 0.90 and distance-corrected (Bianco et al., 2018) Pearson *r*^′^ = 0.72 in HL, *r* = 0.91 and *r*^′^ = 0.64 in ESC, genomic distances > 100*kb*). Subtraction matrices *D* are defined as the simple bin-wise difference *D_ij_* = 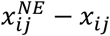, where 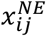 and *x*_*ij*_ are the entries of the contact maps with and without NE respectively.

##### Models of mutants

Polymer models of deletions in HL are simulated as described in (Bianco et al., 2018). Basically, we implement in-silico mutations on the polymer model trained on WT data by deleting the portion corresponding to the deleted chromatin regions in experiments. Specifically, polymer model for ΔD1 has *N* = 2130 beads (i.e. without the region corresponding to D1); analogously, polymer model for ΔD2 has *N* = 2190 beads (i.e. without the region corresponding to D2); finally, polymer model for ΔD1+2 has *N* = 1370 beads and it is much shorter as it carries the deletion of both D1 and D2. For each mutation, a population of 3D polymer structures is then obtained through independent MD simulation performed as described above. DamID data specific for each mutation is integrated in the model to simulate NE. Distances and overlap distributions are generated using mapping coefficients estimated from the WT models.

##### Polymer graphics

Polymer 3D snapshots shown in Figures 4 and S5 are representative single molecule structures taken from real MD simulations. Regions corresponding to Fl1, D1, Rex1R, D2, Fat1, Fl2 are differently colored. A slice of the simulated NE is rendered as a thick spherical wall colored as in FISH imaging. To clarify the relationship between the polymer and NE, each image is presented from the same point-of-view through a geometrically calibrated 3D rotation matrix. For visual purposes, polymers are shown in a coarse-grained version of a smooth third-order polynomial spline passing through bead coordinates.

See Table S4 for a summary of statistical measurements from polymer modelling.

#### Oligopaint FISH image analyses

Image analysis was performed using Fiji and MATLAB (R2018-2019 and image processing toolbox). For *overlap intermingling fraction* and *combined sphericity* measurements, z-stacks of regions of interest (ROIs) of 3×3 μm^2^ surrounding FISH signals were extracted and smoothed using a 3D Gaussian filter (sigma = 0.5 pixel). FISH channels were then segmented in 3D using automatic Otsu’s method. Only ROIs containing 1 FISH segmented object per channel (or at least 1 object for the combined D1+D2 FISH) larger than 0.04 μm^3^ were kept for further analyses. *Object intermingling fraction* of *Rex1R* or *Fat1* with D1+D2 (Figures 4D and S6) was obtained by dividing the overlapping volume between *Rex1R* or *Fat1* and D1+D2 by the volume of *Rex1R* or *Fat1*. *Overlap* fractions (Figure 5B and S6) correspond to the Jaccard Index between the two segmented FISH objects. For *combined sphericity* calculation, FISH segmented objects from the two channels were merged into one, and only ROIs containing 1 merged object were considered for the analysis. *Combined sphericity* ψ was defined as as 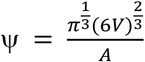 where V is the volume of the segmented object and A its surface area. For distance to lamin analysis, z-stacks of ROIs surrounding individual nuclei were extracted and smoothed using a 3D Gaussian filter (sigma = 0.5 pixel). FISH channels were segmented using a threshold value corresponding to 20% of the maximum pixel intensity. For a given FISH channel, only nuclei containing 2 segmented FISH objects larger than 0.04 μm^3^ were kept for further analysis. For each FISH object, an ROI surrounding its maximum and minimum z-coordinates was extracted and the lamin channel was segmented using Otsu’s method. Lamin segmented objects smaller than 0.02 μm^3^ were discarded and Lamin segmented channel was processed using the MATLAB imfill function. 3D Euclidean distance transform of the segmented Lamin channel was calculated using the MATLAB bwdistsc function and distance to the centroid of the FISH segmented object was extracted.

See Table S4 for a summary of statistical measurements from FISH analyses.

#### Statistical methods

All details of statistical analyses can be found in the figure legends and STAR methods.

## SUPPLEMENTARY TABLES

**Table S1.** List of mutant ESCs with sgRNAs, donor knockin plasmids and genotyping primers indicated. Breakpoint genotyping primers are used to detect deletions (del), site specific integration (ssi) or specific KI-construct (kic).

**Table S2.** List of oligonucleotides for DamID processing, promoter cloning and genotyping.

**Table S3**. Summary of Oligopaint library with regions of interest indicated and oligonucleotide primer sequences. See Figures 4, 5, S5 and S6.

**Table S4.** Summary of statistics for FISH and polymer modelling analyses. See Figures 4, 5, S5 and S6.

**Table S5.** Summary of species used evolutionary comparison. See Figure 2 and S3.

## REFERENCES

1. Abdennur, N., and Mirny, L.A. (2020). Cooler: scalable storage for Hi-C data and other genomically labeled arrays. Bioinformatics 36, 311–316.

2. Acemel, R.D., Maeso, I., and Gomez-Skarmeta, J.L. (2017). Topologically associated domains: a successful scaffold for the evolution of gene regulation in animals. Wiley Interdisciplinary Reviews-Developmental Biology 6.

3. Alam, T., Agrawal, S., Severin, J., Young, R.S., Andersson, R., Arner, E., Hasegawa, A., Lizio, M., Ramilowski, J.A., Abugessaisa, I., et al. (2020). Comparative transcriptomics of primary cells in vertebrates. Genome Res 30, 951–961.

4. Anderson, E., Devenney, P.S., Hill, R.E., and Lettice, L.A. (2014). Mapping the Shh long-range regulatory domain. Development 141, 3934–3943.

5. Andrey, G., Montavon, T., Mascrez, B., Gonzalez, F., Noordermeer, D., Leleu, M., Trono, D., Spitz, F., and Duboule, D. (2013). A switch between topological domains underlies HoxD genes collinearity in mouse limbs. Science 340, 1234167.

6. Andrey, G., and Mundlos, S. (2017). The three-dimensional genome: regulating gene expression during pluripotency and development. Development 144, 3646–3658.

7. Andrey, G., Schopflin, R., Jerkovic, I., Heinrich, V., Ibrahim, D.M., Paliou, C., Hochradel, M., Timmermann, B., Haas, S., Vingron, M., et al. (2017). Characterization of hundreds of regulatory landscapes in developing limbs reveals two regimes of chromatin folding. Genome Res 27, 223–233.

8. Andrey, G., and Spielmann, M. (2017). CRISPR/Cas9 Genome Editing in Embryonic Stem Cells. Methods Mol Biol 1468, 221–234.

9. Artus, J., and Hadjantonakis, A.K. (2011). Generation of chimeras by aggregation of embryonic stem cells with diploid or tetraploid mouse embryos. Methods Mol Biol 693, 37–56.

10. Ball, G., Demmerle, J., Kaufmann, R., Davis, I., Dobbie, I.M., and Schermelleh, L. (2015). SIMcheck: a Toolbox for Successful Super-resolution Structured Illumination Microscopy. Sci Rep 5, 15915.

11. Baranasic, D., Hörtenhuber, M., Balwierz, P., Zehnder, T., Mukarram, A.K., Nepal, C., Varnai, C., Hadzhiev, Y., Jimenez-Gonzalez, A., Li, N., et al. (2021). Integrated annotation and analysis of genomic features reveal new types of functional elements and large-scale epigenetic phenomena in the developing zebrafish. bioRxiv, 2021.2008.2009.454869.

12. Barbieri, M., Chotalia, M., Fraser, J., Lavitas, L.M., Dostie, J., Pombo, A., and Nicodemi, M. (2012). Complexity of chromatin folding is captured by the strings and binders switch model. Proc Natl Acad Sci U S A 109, 16173–16178.

13. Bauer, M., Vidal, E., Zorita, E., Uresin, N., Pinter, S.F., Filion, G.J., and Payer, B. (2021). Chromosome compartments on the inactive X guide TAD formation independently of transcription during X-reactivation. Nat Commun 12, 3499.

14. Beard, C., Hochedlinger, K., Plath, K., Wutz, A., and Jaenisch, R. (2006). Efficient method to generate single-copy transgenic mice by site-specific integration in embryonic stem cells. Genesis 44, 23–28.

15. Beliveau, B.J., Boettiger, A.N., Avendano, M.S., Jungmann, R., McCole, R.B., Joyce, E.F., Kim-Kiselak, C., Bantignies, F., Fonseka, C.Y., Erceg, J., et al. (2015). Single-molecule super-resolution imaging of chromosomes and in situ haplotype visualization using Oligopaint FISH probes. Nat Commun 6, 7147.

16. Beliveau, B.J., Kishi, J.Y., Nir, G., Sasaki, H.M., Saka, S.K., Nguyen, S.C., Wu, C.T., and Yin, P. (2018). OligoMiner provides a rapid, flexible environment for the design of genome-scale oligonucleotide in situ hybridization probes. Proc Natl Acad Sci U S A 115, E2183–E2192.

17. Bianco, S., Lupianez, D.G., Chiariello, A.M., Annunziatella, C., Kraft, K., Schopflin, R., Wittler, L., Andrey, G., Vingron, M., Pombo, A., et al. (2018). Polymer physics predicts the effects of structural variants on chromatin architecture. Nat Genet 50, 662–667.

18. Bonev, B., and Cavalli, G. (2016). Organization and function of the 3D genome. Nature reviews Genetics 17, 772.

19. Bonev, B., Cohen, N.M., Szabo, Q., Fritsch, L., Papadopoulos, G.L., Lubling, Y., Xu, X.L., Lv, X.D., Hugnot, J.P., Tanay, A., et al. (2017). Multiscale 3D Genome Rewiring during Mouse Neural Development. Cell 171, 557-+.

20. Bonora, G., Ramani, V., Singh, R., Fang, H., Jackson, D., Srivatsan, S., Qiu, R., Lee, C., Trapnell, C., Shendure, J., et al. (2020). Single-cell landscape of nuclear configuration and gene expression during stem cell differentiation and X inactivation. bioRxiv, 2020.2011.2020.390765.

21. Borgel, J., Guibert, S., Li, Y., Chiba, H., Schubeler, D., Sasaki, H., Forne, T., and Weber, M. (2010). Targets and dynamics of promoter DNA methylation during early mouse development. Nat Genet 42, 1093–1100.

22. Brueckner, L., Zhao, P.A., van Schaik, T., Leemans, C., Sima, J., Peric-Hupkes, D., Gilbert, D.M., and van Steensel, B. (2020). Local rewiring of genome-nuclear lamina interactions by transcription. Embo J 39, e103159.

23. Buenrostro, J.D., Wu, B., Chang, H.Y., and Greenleaf, W.J. (2015). ATAC-seq: A Method for Assaying Chromatin Accessibility Genome-Wide. Curr Protoc Mol Biol 109, 21 29 21–21 29 29.

24. Cao, J., Spielmann, M., Qiu, X., Huang, X., Ibrahim, D.M., Hill, A.J., Zhang, F., Mundlos, S., Christiansen, L., Steemers, F.J., et al. (2019). The single-cell transcriptional landscape of mammalian organogenesis. Nature 566, 496–502.

25. Chiariello, A.M., Annunziatella, C., Bianco, S., Esposito, A., and Nicodemi, M. (2016). Polymer physics of chromosome large-scale 3D organisation. Sci Rep 6, 29775.

26. Choi, I., Oh, J., Cho, B.N., Ahnn, J., Jung, Y.K., Han Kim, D., and Cho, C. (2004). Characterization and comparative genomic analysis of intronless Adams with testicular gene expression. Genomics 83, 636–646.

27. Chubb, J.R., Boyle, S., Perry, P., and Bickmore, W.A. (2002). Chromatin motion is constrained by association with nuclear compartments in human cells. Current Biology 12, 439–445.

28. Ciani, L., Patel, A., Allen, N.D., and ffrench-Constant, C. (2003). Mice lacking the giant protocadherin mFAT1 exhibit renal slit junction abnormalities and a partially penetrant cyclopia and anophthalmia phenotype. Mol Cell Biol 23, 3575–3582.

29. Consortium, F., the, R.P., Clst, Forrest, A.R., Kawaji, H., Rehli, M., Baillie, J.K., de Hoon, M.J., Haberle, V., Lassmann, T., et al. (2014). A promoter-level mammalian expression atlas. Nature 507, 462–470.

30. Deng, W., Rupon, J.W., Krivega, I., Breda, L., Motta, I., Jahn, K.S., Reik, A., Gregory, P.D., Rivella, S., Dean, A., et al. (2014). Reactivation of developmentally silenced globin genes by forced chromatin looping. Cell 158, 849–860.

31. Despang, A., Schöpflin, R., Franke, M., Ali, S., Jerkovic, I., Paliou, C., Chan, W.-L., Timmermann, B., Wittler, L., Vingron, M., et al. (2019). Functional dissection of TADs reveals non-essential and instructive roles in regulating gene expression. bioRxiv, 566562.

32. Dijkstra, E.W. (1959). A note on two problems in connexion with graphs. Numerische Mathematik 1, 269–271.

33. Dixon, J.R., Gorkin, D.U., and Ren, B. (2016). Chromatin Domains: The Unit of Chromosome Organization. Molecular Cell 62, 668–680.

34. Dixon, J.R., Selvaraj, S., Yue, F., Kim, A., Li, Y., Shen, Y., Hu, M., Liu, J.S., and Ren, B. (2012). Topological domains in mammalian genomes identified by analysis of chromatin interactions. Nature 485, 376–380.

35. Dobin, A., Davis, C.A., Schlesinger, F., Drenkow, J., Zaleski, C., Jha, S., Batut, P., Chaisson, M., and Gingeras, T.R. (2013). STAR: ultrafast universal RNA-seq aligner. Bioinformatics 29, 15–21.

36. Durand, N.C., Shamim, M.S., Machol, I., Rao, S.S., Huntley, M.H., Lander, E.S., and Aiden, E.L. (2016). Juicer Provides a One-Click System for Analyzing Loop-Resolution Hi-C Experiments. Cell Syst 3, 95–98.

37. Eden, E., Navon, R., Steinfeld, I., Lipson, D., and Yakhini, Z. (2009). GOrilla: a tool for discovery and visualization of enriched GO terms in ranked gene lists. BMC Bioinformatics 10, 48.

38. Falk, M., Feodorova, Y., Naumova, N., Imakaev, M., Lajoie, B.R., Leonhardt, H., Joffe, B., Dekker, J., Fudenberg, G., Solovei, I., et al. (2019). Heterochromatin drives compartmentalization of inverted and conventional nuclei. Nature 570, 395–399.

39. Finlan, L.E., Sproul, D., Thomson, I., Boyle, S., Kerr, E., Perry, P., Ylstra, B., Chubb, J.R., and Bickmore, W.A. (2008). Recruitment to the nuclear periphery can alter expression of genes in human cells. PLoS Genetics 4, e1000039.

40. Flavahan, W.A., Drier, Y., Liau, B.B., Gillespie, S.M., Venteicher, A.S., Stemmer-Rachamimov, A.O., Suva, M.L., and Bernstein, B.E. (2016). Insulator dysfunction and oncogene activation in IDH mutant gliomas. Nature 529, 110–114.

41. Fraser, J., Ferrai, C., Chiariello, A.M., Schueler, M., Rito, T., Laudanno, G., Barbieri, M., Moore, B.L., Kraemer, D.C., Aitken, S., et al. (2015). Hierarchical folding and reorganization of chromosomes are linked to transcriptional changes in cellular differentiation. Molecular systems biology 11, 852.

42. Fudenberg, G., Imakaev, M., Lu, C., Goloborodko, A., Abdennur, N., and Mirny, L.A. (2016). Formation of Chromosomal Domains by Loop Extrusion. Cell Rep 15, 2038–2049.

43. Fulco, C.P., Nasser, J., Jones, T.R., Munson, G., Bergman, D.T., Subramanian, V., Grossman, S.R., Anyoha, R., Doughty, B.R., Patwardhan, T.A., et al. (2019). Activity-by-contact model of enhancer-promoter regulation from thousands of CRISPR perturbations. Nat Genet 51, 1664–1669.

44. Furlong, E.E.M., and Levine, M. (2018). Developmental enhancers and chromosome topology. Science 361, 1341–1345.

45. George, S.H., Gertsenstein, M., Vintersten, K., Korets-Smith, E., Murphy, J., Stevens, M.E., Haigh, J.J., and Nagy, A. (2007). Developmental and adult phenotyping directly from mutant embryonic stem cells. Proc Natl Acad Sci U S A 104, 4455–4460.

46. Gjaltema, R.A.F., Schwämmle, T., Kautz, P., Robson, M., Schöpflin, R., Lustig, L.R., Brandenburg, L., Dunkel, I., Vechiatto, C., Ntini, E., et al. (2021). Distal and proximal cis-regulatory elements sense X-chromosomal dosage and developmental state at the *Xist* locus. bioRxiv, 2021.2003.2029.437476.

47. Gorkin, D.U., Barozzi, I., Zhao, Y., Zhang, Y., Huang, H., Lee, A.Y., Li, B., Chiou, J., Wildberg, A., Ding, B., et al. (2020). An atlas of dynamic chromatin landscapes in mouse fetal development. Nature 583, 744–751.

48. Grosswendt, S., Kretzmer, H., Smith, Z.D., Kumar, A.S., Hetzel, S., Wittler, L., Klages, S., Timmermann, B., Mukherji, S., and Meissner, A. (2020). Epigenetic regulator function through mouse gastrulation. Nature.

49. Gustafsson, M.G., Shao, L., Carlton, P.M., Wang, C.J., Golubovskaya, I.N., Cande, W.Z., Agard, D.A., and Sedat, J.W. (2008). Three-dimensional resolution doubling in wide-field fluorescence microscopy by structured illumination. Biophys J 94, 4957–4970.

50. Harmston, N., Ing-Simmons, E., Tan, G., Perry, M., Merkenschlager, M., and Lenhard, B. (2017). Topologically associating domains are ancient features that coincide with Metazoan clusters of extreme noncoding conservation. Nat Commun 8, 441.

51. Helmbacher, F. (2018). Tissue-specific activities of the Fat1 cadherin cooperate to control neuromuscular morphogenesis. PLoS Biol 16, e2004734.

52. Huang, P., Keller, C.A., Giardine, B., Grevet, J.D., Davies, J.O.J., Hughes, J.R., Kurita, R., Nakamura, Y., Hardison, R.C., and Blobel, G.A. (2017). Comparative analysis of three-dimensional chromosomal architecture identifies a novel fetal hemoglobin regulatory element. Genes Dev 31, 1704–1713.

53. Juhling, F., Kretzmer, H., Bernhart, S.H., Otto, C., Stadler, P.F., and Hoffmann, S. (2016). metilene: fast and sensitive calling of differentially methylated regions from bisulfite sequencing data. Genome Res 26, 256–262.

54. Kane, L., Williamson, I., Flyamer, I.M., Kumar, Y., Hill, R.E., Lettice, L.A., and Bickmore, W.A. (2021). Cohesin is required for long-range enhancer action. bioRxiv, 2021.2006.2024.449812.

55. Kechin, A., Boyarskikh, U., Kel, A., and Filipenko, M. (2017). cutPrimers: A New Tool for Accurate Cutting of Primers from Reads of Targeted Next Generation Sequencing. J Comput Biol 24, 1138–1143.

56. Kim, J.D., Faulk, C., and Kim, J. (2007). Retroposition and evolution of the DNA-binding motifs of YY1, YY2 and REX1. Nucleic Acids Res 35, 3442-3452.

57. Kim, J.D., Kim, H., Ekram, M.B., Yu, S., Faulk, C., and Kim, J. (2011). Rex1/Zfp42 as an epigenetic regulator for genomic imprinting. Hum Mol Genet 20, 1353–1362.

58. Klosen, P., Lapmanee, S., Schuster, C., Guardiola, B., Hicks, D., Pevet, P., and Felder-Schmittbuhl, M.P. (2019). MT1 and MT2 melatonin receptors are expressed in nonoverlapping neuronal populations. J Pineal Res 67, e12575.

59. Knight, P.A., and Ruiz, D. (2012). A fast algorithm for matrix balancing. IMA Journal of Numerical Analysis 33, 1029–1047.

60. Kraft, K., Geuer, S., Will, A.J., Chan, W.L., Paliou, C., Borschiwer, M., Harabula, I., Wittler, L., Franke, M., Ibrahim, D.M., et al. (2015). Deletions, Inversions, Duplications: Engineering of Structural Variants using CRISPR/Cas in Mice. Cell reports.

61. Kraft, K., Magg, A., Heinrich, V., Riemenschneider, C., Schopflin, R., Markowski, J., Ibrahim, D.M., Acuna-Hidalgo, R., Despang, A., Andrey, G., et al. (2019). Serial genomic inversions induce tissue-specific architectural stripes, gene misexpression and congenital malformations. Nat Cell Biol 21, 305–310.

62. Kragesteen, B.K., Spielmann, M., Paliou, C., Heinrich, V., Schopflin, R., Esposito, A., Annunziatella, C., Bianco, S., Chiariello, A.M., Jerkovic, I., et al. (2018). Dynamic 3D chromatin architecture contributes to enhancer specificity and limb morphogenesis. Nat Genet 50, 1463–1473.

63. Krefting, J., Andrade-Navarro, M.A., and Ibn-Salem, J. (2018). Evolutionary stability of topologically associating domains is associated with conserved gene regulation. BMC Biol 16, 87.

64. Kremer, K., and Grest, G.S. (1990). Dynamics of entangled linear polymer melts: A molecular-dynamics simulation. The Journal of Chemical Physics 92, 5057–5086.

65. Langmead, B., and Salzberg, S.L. (2012). Fast gapped-read alignment with Bowtie 2. Nat Methods 9, 357–359.

66. Laugsch, M., Bartusel, M., Rehimi, R., Alirzayeva, H., Karaolidou, A., Crispatzu, G., Zentis, P., Nikolic, M., Bleckwehl, T., Kolovos, P., et al. (2019). Modeling the Pathological Long-Range Regulatory Effects of Human Structural Variation with Patient-Specific hiPSCs. Cell Stem Cell 24, 736–752 e712.

67. Leemans, C., van der Zwalm, M.C.H., Brueckner, L., Comoglio, F., van Schaik, T., Pagie, L., van Arensbergen, J., and van Steensel, B. (2019). Promoter-Intrinsic and Local Chromatin Features Determine Gene Repression in LADs. Cell 177, 852–864 e814.

68. Li, H., and Durbin, R. (2009). Fast and accurate short read alignment with Burrows-Wheeler transform. Bioinformatics 25, 1754–1760.

69. Li, H., and Durbin, R. (2010). Fast and accurate long-read alignment with Burrows-Wheeler transform. Bioinformatics 26, 589–595.

70. Li, H., Handsaker, B., Wysoker, A., Fennell, T., Ruan, J., Homer, N., Marth, G., Abecasis, G., Durbin, R., and Genome Project Data Processing, S. (2009). The Sequence Alignment/Map format and SAMtools. Bioinformatics 25, 2078–2079.

71. Li, X., and Noll, M. (1994). Compatibility between enhancers and promoters determines the transcriptional specificity of gooseberry and gooseberry neuro in the Drosophila embryo. Embo J 13, 400–406.

72. Liu, N.Q., Maresca, M., van den Brand, T., Braccioli, L., Schijns, M., Teunissen, H., Bruneau, B.G., Nora, E.P., and de Wit, E. (2021). WAPL maintains a cohesin loading cycle to preserve cell-type-specific distal gene regulation. Nat Genet 53, 100–109.

73. Lizio, M., Harshbarger, J., Shimoji, H., Severin, J., Kasukawa, T., Sahin, S., Abugessaisa, I., Fukuda, S., Hori, F., Ishikawa-Kato, S., et al. (2015). Gateways to the FANTOM5 promoter level mammalian expression atlas. Genome Biol 16, 22.

74. Lobe, C.G., Koop, K.E., Kreppner, W., Lomeli, H., Gertsenstein, M., and Nagy, A. (1999). Z/AP, a double reporter for cre-mediated recombination. Dev Biol 208, 281–292.

75. Long, H.K., Prescott, S.L., and Wysocka, J. (2016). Ever-Changing Landscapes: Transcriptional Enhancers in Development and Evolution. Cell 167, 1170–1187.

76. Love, M.I., Huber, W., and Anders, S. (2014). Moderated estimation of fold change and dispersion for RNA-seq data with DESeq2. Genome Biol 15, 550.

77. Marinic, M., Aktas, T., Ruf, S., and Spitz, F. (2013). An integrated holo-enhancer unit defines tissue and gene specificity of the Fgf8 regulatory landscape. Dev Cell 24, 530–542.

78. Marsh, B., and Blelloch, R. (2020). Single nuclei RNA-seq of mouse placental labyrinth development. Elife 9.

79. Martin, M. (2011). Cutadapt removes adapter sequences from high-throughput sequencing reads. 2011 17, 3.

80. Masui, S., Ohtsuka, S., Yagi, R., Takahashi, K., Ko, M.S., and Niwa, H. (2008). Rex1/Zfp42 is dispensable for pluripotency in mouse ES cells. BMC Dev Biol 8, 45.

81. McKenna, A., Hanna, M., Banks, E., Sivachenko, A., Cibulskis, K., Kernytsky, A., Garimella, K., Altshuler, D., Gabriel, S., Daly, M., et al. (2010). The Genome Analysis Toolkit: a MapReduce framework for analyzing next-generation DNA sequencing data. Genome Res 20, 1297–1303.

82. Melo, U.S., Schopflin, R., Acuna-Hidalgo, R., Mensah, M.A., Fischer-Zirnsak, B., Holtgrewe, M., Klever, M.K., Turkmen, S., Heinrich, V., Pluym, I.D., et al. (2020). Hi-C Identifies Complex Genomic Rearrangements and TAD-Shuffling in Developmental Diseases. Am J Hum Genet 106, 872–884.

83. Merli, C., Bergstrom, D.E., Cygan, J.A., and Blackman, R.K. (1996). Promoter specificity mediates the independent regulation of neighboring genes. Genes Dev 10, 1260–1270.

84. Nasser, J., Bergman, D.T., Fulco, C.P., Guckelberger, P., Doughty, B.R., Patwardhan, T.A., Jones, T.R., Nguyen, T.H., Ulirsch, J.C., Lekschas, F., et al. (2021). Genome-wide enhancer maps link risk variants to disease genes. Nature 593, 238–243.

85. Nicodemi, M., and Prisco, A. (2009). Thermodynamic pathways to genome spatial organization in the cell nucleus. Biophys J 96, 2168–2177.

86. Noordermeer, D., Leleu, M., Splinter, E., Rougemont, J., De Laat, W., and Duboule, D. (2011). The dynamic architecture of Hox gene clusters. Science 334, 222–225.

87. Nora, E.P., Goloborodko, A., Valton, A.L., Gibcus, J.H., Uebersohn, A., Abdennur, N., Dekker, J., Mirny, L.A., and Bruneau, B.G. (2017). Targeted Degradation of CTCF Decouples Local Insulation of Chromosome Domains from Genomic Compartmentalization. Cell 169, 930–944 e922.

88. Nora, E.P., Lajoie, B.R., Schulz, E.G., Giorgetti, L., Okamoto, I., Servant, N., Piolot, T., van Berkum, N.L., Meisig, J., Sedat, J., et al. (2012). Spatial partitioning of the regulatory landscape of the X-inactivation centre. Nature 485, 381–385.

89. Nuebler, J., Fudenberg, G., Imakaev, M., Abdennur, N., and Mirny, L.A. (2018). Chromatin organization by an interplay of loop extrusion and compartmental segregation. Proc Natl Acad Sci U S A 115, E6697–E6706.

90. Okano, M., Bell, D.W., Haber, D.A., and Li, E. (1999). DNA methyltransferases Dnmt3a and Dnmt3b are essential for de novo methylation and mammalian development. Cell 99, 247–257.

91. Ou, H.D., Phan, S., Deerinck, T.J., Thor, A., Ellisman, M.H., and O’Shea, C.C. (2017). ChromEMT: Visualizing 3D chromatin structure and compaction in interphase and mitotic cells. Science 357.

92. Palstra, R.J., Tolhuis, B., Splinter, E., Nijmeijer, R., Grosveld, F., and de Laat, W. (2003). The beta-globin nuclear compartment in development and erythroid differentiation. Nature Genetics 35, 190–194.

93. Peng, Z., Gong, Y., and Liang, X. (2021). Role of FAT1 in health and disease. Oncol Lett 21, 398.

94. Pijuan-Sala, B., Griffiths, J.A., Guibentif, C., Hiscock, T.W., Jawaid, W., Calero-Nieto, F.J., Mulas, C., Ibarra-Soria, X., Tyser, R.C.V., Ho, D.L.L., et al. (2019). A single-cell molecular map of mouse gastrulation and early organogenesis. Nature 566, 490-+.

95. Ramirez, F., Ryan, D.P., Gruning, B., Bhardwaj, V., Kilpert, F., Richter, A.S., Heyne, S., Dundar, F., and Manke, T. (2016). deepTools2: a next generation web server for deep-sequencing data analysis. Nucleic Acids Res 44, W160–165.

96. Ramisch, A., Heinrich, V., Glaser, L.V., Fuchs, A., Yang, X., Benner, P., Schopflin, R., Li, N., Kinkley, S., Romer-Hillmann, A., et al. (2019). CRUP: a comprehensive framework to predict condition-specific regulatory units. Genome Biol 20, 227.

97. Rao, S.S., Huntley, M.H., Durand, N.C., Stamenova, E.K., Bochkov, I.D., Robinson, J.T., Sanborn, A.L., Machol, I., Omer, A.D., Lander, E.S., et al. (2014). A 3D map of the human genome at kilobase resolution reveals principles of chromatin looping. Cell 159, 1665–1680.

98. Rao, S.S.P., Huang, S.C., Glenn St Hilaire, B., Engreitz, J.M., Perez, E.M., Kieffer-Kwon, K.R., Sanborn, A.L., Johnstone, S.E., Bascom, G.D., Bochkov, I.D., et al. (2017). Cohesin Loss Eliminates All Loop Domains. Cell 171, 305–320 e324.

99. Real, F.M., Haas, S.A., Franchini, P., Xiong, P., Simakov, O., Kuhl, H., Schopflin, R., Heller, D., Moeinzadeh, M.H., Heinrich, V., et al. (2020). The mole genome reveals regulatory rearrangements associated with adaptive intersexuality. Science 370, 208–214.

100. Reddy, K.L., Zullo, J.M., Bertolino, E., and Singh, H. (2008). Transcriptional repression mediated by repositioning of genes to the nuclear lamina. Nature 452, 243–247.

101. Rennie, S., Dalby, M., van Duin, L., and Andersson, R. (2018). Transcriptional decomposition reveals active chromatin architectures and cell specific regulatory interactions. Nat Commun 9, 487.

102. Ribeiro, D.M., Rubinacci, S., Ramisch, A., Hofmeister, R.J., Dermitzakis, E.T., and Delaneau, O. (2021). The molecular basis, genetic control and pleiotropic effects of local gene co-expression. Nat Commun 12, 4842.

103. Robson, M.I., de Las Heras, J.I., Czapiewski, R., Le Thanh, P., Booth, D.G., Kelly, D.A., Webb, S., Kerr, A.R.W., and Schirmer, E.C. (2016). Tissue-Specific Gene Repositioning by Muscle Nuclear Membrane Proteins Enhances Repression of Critical Developmental Genes during Myogenesis. Mol Cell 62, 834–847.

104. Robson, M.I., de Las Heras, J.I., Czapiewski, R., Sivakumar, A., Kerr, A.R.W., and Schirmer, E.C. (2017). Constrained release of lamina-associated enhancers and genes from the nuclear envelope during T-cell activation facilitates their association in chromosome compartments. Genome Res 27, 1126–1138.

105. Robson, M.I., Ringel, A.R., and Mundlos, S. (2019). Regulatory Landscaping: How Enhancer-Promoter Communication Is Sculpted in 3D. Mol Cell 74, 1110–1122.

106. Robson, M.I., and Schirmer, E.C. (2016). The Application of DamID to Identify Peripheral Gene Sequences in Differentiated and Primary Cells. Methods Mol Biol 1411, 359–386.

107. Sadeqzadeh, E., de Bock, C.E., and Thorne, R.F. (2014). Sleeping giants: emerging roles for the fat cadherins in health and disease. Med Res Rev 34, 190–221.

108. Sanborn, A.L., Rao, S.S., Huang, S.C., Durand, N.C., Huntley, M.H., Jewett, A.I., Bochkov, I.D., Chinnappan, D., Cutkosky, A., Li, J., et al. (2015). Chromatin extrusion explains key features of loop and domain formation in wild-type and engineered genomes. Proc Natl Acad Sci U S A 112, E6456–6465.

109. Schermelleh, L., Carlton, P.M., Haase, S., Shao, L., Winoto, L., Kner, P., Burke, B., Cardoso, M.C., Agard, D.A., Gustafsson, M.G., et al. (2008). Subdiffraction multicolor imaging of the nuclear periphery with 3D structured illumination microscopy. Science 320, 1332–1336.

110. Schmidl, C., Rendeiro, A.F., Sheffield, N.C., and Bock, C. (2015). ChIPmentation: fast, robust, low-input ChIP-seq for histones and transcription factors. Nat Methods 12, 963–965.

111. Schwarzer, W., Abdennur, N., Goloborodko, A., Pekowska, A., Fudenberg, G., Loe-Mie, Y., Fonseca, N.A., Huber, W., C, H.H., Mirny, L., et al. (2017). Two independent modes of chromatin organization revealed by cohesin removal. Nature 551, 51–56.

112. Shen, Y., Yue, F., McCleary, D.F., Ye, Z., Edsall, L., Kuan, S., Wagner, U., Dixon, J., Lee, L., Lobanenkov, V.V., et al. (2012). A map of the cis-regulatory sequences in the mouse genome. Nature 488, 116–120.

113. Shima, Y., Sugino, K., Hempel, C.M., Shima, M., Taneja, P., Bullis, J.B., Mehta, S., Lois, C., and Nelson, S.B. (2016). A Mammalian enhancer trap resource for discovering and manipulating neuronal cell types. Elife 5, e13503.

114. Shin, H., Shi, Y., Dai, C., Tjong, H., Gong, K., Alber, F., and Zhou, X.J. (2016). TopDom: an efficient and deterministic method for identifying topological domains in genomes. Nucleic Acids Res 44, e70.

115. Sima, J., Chakraborty, A., Dileep, V., Michalski, M., Klein, K.N., Holcomb, N.P., Turner, J.L., Paulsen, M.T., Rivera-Mulia, J.C., Trevilla-Garcia, C., et al. (2019). Identifying cis Elements for Spatiotemporal Control of Mammalian DNA Replication. Cell 176, 816-+.

116. Soshnikova, N., and Duboule, D. (2009). Epigenetic temporal control of mouse Hox genes in vivo. Science 324, 1320–1323.

117. Spielmann, M., Lupianez, D.G., and Mundlos, S. (2018). Structural variation in the 3D genome. Nat Rev Genet 19, 453–467.

118. Sun, D., Xi, Y., Rodriguez, B., Park, H.J., Tong, P., Meong, M., Goodell, M.A., and Li, W. (2014). MOABS: model based analysis of bisulfite sequencing data. Genome Biol 15, R38.

119. Symmons, O., Uslu, V.V., Tsujimura, T., Ruf, S., Nassari, S., Schwarzer, W., Ettwiller, L., and Spitz, F. (2014). Functional and topological characteristics of mammalian regulatory domains. Genome Res 24, 390–400.

120. Szabo, Q., Donjon, A., Jerkovic, I., Papadopoulos, G.L., Cheutin, T., Bonev, B., Nora, E.P., Bruneau, B.G., Bantignies, F., and Cavalli, G. (2020). Regulation of single-cell genome organization into TADs and chromatin nanodomains. Nat Genet 52, 1151–1157.

121. Szabo, Q., Jost, D., Chang, J.M., Cattoni, D.I., Papadopoulos, G.L., Bonev, B., Sexton, T., Gurgo, J., Jacquier, C., Nollmann, M., et al. (2018). TADs are 3D structural units of higher-order chromosome organization in Drosophila. Sci Adv 4, eaar8082.

122. Takebayashi, S., Dileep, V., Ryba, T., Dennis, J.H., and Gilbert, D.M. (2012). Chromatin-interaction compartment switch at developmentally regulated chromosomal domains reveals an unusual principle of chromatin folding. P Natl Acad Sci USA 109, 12574–12579.

123. Therizols, P., Illingworth, R.S., Courilleau, C., Boyle, S., Wood, A.J., and Bickmore, W.A. (2014). Chromatin decondensation is sufficient to alter nuclear organization in embryonic stem cells. Science 346, 1238–1242.

124. van Arensbergen, J., van Steensel, B., and Bussemaker, H.J. (2014). In search of the determinants of enhancer-promoter interaction specificity. Trends Cell Biol 24, 695–702.

125. van Schaik, T., Vos, M., Peric-Hupkes, D., Hn Celie, P., and van Steensel, B. (2020). Cell cycle dynamics of lamina-associated DNA. EMBO Rep 21, e50636.

126. van Steensel, B., and Belmont, A.S. (2017). Lamina-Associated Domains: Links with Chromosome Architecture, Heterochromatin, and Gene Repression. Cell 169, 780–791.

127. Vogel, M.J., Peric-Hupkes, D., and van Steensel, B. (2007). Detection of in vivo protein-DNA interactions using DamID in mammalian cells. Nature protocols 2, 1467–1478.

128. Wingett, S., Ewels, P., Furlan-Magaril, M., Nagano, T., Schoenfelder, S., Fraser, P., and Andrews, S. (2015). HiCUP: pipeline for mapping and processing Hi-C data. F1000Res 4, 1310.

129. Wu, H.J., Landshammer, A., Stamenova, E.K., Bolondi, A., Kretzmer, H., Meissner, A., and Michor, F. (2021). Topological isolation of developmental regulators in mammalian genomes. Nat Commun 12, 4897.

130. Xi, Y., and Li, W. (2009). BSMAP: whole genome bisulfite sequence MAPping program. BMC Bioinformatics 10, 232.

131. Yagi, M., Kabata, M., Tanaka, A., Ukai, T., Ohta, S., Nakabayashi, K., Shimizu, M., Hata, K., Meissner, A., Yamamoto, T., et al. (2020). Identification of distinct loci for de novo DNA methylation by DNMT3A and DNMT3B during mammalian development. Nat Commun 11, 3199.

132. Zehnder, T., Benner, P., and Vingron, M. (2019). Predicting enhancers in mammalian genomes using supervised hidden Markov models. BMC Bioinformatics 20, 157.

133. Zhan, Y., Mariani, L., Barozzi, I., Schulz, E.G., Bluthgen, N., Stadler, M., Tiana, G., and Giorgetti, L. (2017). Reciprocal insulation analysis of Hi-C data shows that TADs represent a functionally but not structurally privileged scale in the hierarchical folding of chromosomes. Genome Res 27, 479–490.

134. Zhang, X., Pavlicev, M., Jones, H.N., and Muglia, L.J. (2020). Eutherian-Specific Gene TRIML2 Attenuates Inflammation in the Evolution of Placentation. Mol Biol Evol 37, 507–523.

135. Zhang, Y., Li, T., Preissl, S., Amaral, M.L., Grinstein, J.D., Farah, E.N., Destici, E., Qiu, Y., Hu, R., Lee, A.Y., et al. (2019). Transcriptionally active HERV-H retrotransposons demarcate topologically associating domains in human pluripotent stem cells. Nat Genet 51, 1380–1388.

136. Zuin, J., Roth, G., Zhan, Y., Cramard, J., Redolfi, J., Piskadlo, E., Mach, P., Kryzhanovska, M., Tihanyi, G., Kohler, H., et al. (2021). Nonlinear control of transcription through enhancer-promoter interactions. bioRxiv, 2021.2004.2022.440891.

